# Towards understanding nitrate-reducing Fe(II) oxidation in *Ferrigenium straubiae*: spotlight on old and new key protein candidates

**DOI:** 10.64898/2026.01.22.701106

**Authors:** Stefanie Becker, Daniel Straub, James Hemp, Andreas Kappler

## Abstract

3.

Autotrophic Fe(II) oxidation is a key microbial process linking iron cycling to global carbon and nitrogen turnover in anoxic and low-oxygen environments. While Fe(II) oxidation, and electron utilization by canonical denitrification or oxygen-reducing pathways have been proposed, the whole electron transport chain via the periplasm is unknown.

To address this gap, we compared the transcriptome of *Ferrigenium straubiae* strain KS (= KCTC 25982; = DSM 118991) cultivated under autotrophic conditions using Fe(II) as the electron donor and either nitrate or oxygen as the terminal electron acceptor. Additionally, we reanalyzed the genome using bioinformatic tools (FeGenie, MHCScan, Foldseek, AlphaFold, COFACTOR, and InterProScan) to classify known, putative, and previously uncharacterized proteins.

Using these analyses, we (i) identified a form II RubisCO-mediated, sedoheptulose-7-phosphate-forming transaldolase variant of the Calvin-Benson-Bassham cycle, (ii) revealed the metabolic possibility of a reversed oxidative TCA cycle and an alternative CO_2_ fixation route via 10-formyl-tetrahydrofolate, (iii) found novel structural features of the proposed Fe(II) oxidases (Cyc2 and MtoA/B: β-barrel outer membrane cytochromes), (iv) uncovered a nitric oxide reductase (eNOR-family) and two redox complexes specifically upregulated under nitrate-reducing Fe(II)-oxidizing conditions, including one comprising a sphaeroides heme protein homolog, and (v) reveal a cytochrome *c* rich gene area encoding ten heme proteins and (iv) proposed a conserved nonaheme cytochrome *c* to link Fe(II) oxidation to quinone reduction.

**Repositories:** GenBank whole genome sequence of strain KS: JAGRPI00000000

IMG Taxon ID of strain KS: 2878407288

NCBI BioProject ID of RNA-Seq data: PRJNA1399691

Strain availability: KCTC 25982 and DSM 118991

Normalized transcripts and Log2FoldChanges (Excel file), and supplementary information are available with the online version of this article.

## 4. Introduction

Bacterial nitrate-reducing Fe(II) oxidation is an environmental process coupling the geochemical nitrogen and iron cycle, taking place in soils, sediments, and aquatic habitats. Nitrate-reduction leads to climate gas emission, while iron speciation and mineralogy has a strong influence on ecosystem functioning. Both make nitrate-reducing Fe(II) oxidation curtail to the understanding of Earth’s present and future state. However, because the metabolic pathway of nitrate-reducing Fe(II) oxidation is poorly understood and incomplete, genomic data derived directly from nitrate polluted field sites cannot yet be reliable interpreted.

The first neutrophilic Fe(II)-oxidizing bacteria (NRFeOx) capable of coupling nitrate reduction to Fe(II) oxidation were described in 1996 (Hafenbradl *et al*. 1996, Straub *et al*. 1996). Since then, Fe(II) oxidation in the presence of nitrate by denitrifying microbial cultures has been frequently observed (Bryce *et al*. 2018). While autotrophic nitrate-reducing Fe(II)-oxidizers are able to oxidize Fe(II) under autotrophic conditions, other denitrifying bacterial cultures in which Fe(II) oxidation was observed require the addition of organic substrates. Based on genomic analysis, growth experiments, and kinetic modeling, these organotrophic cultures linked to Fe(II) oxidation have been classified as either mixotrophic NRFeOx or heterotrophic denitrifiers. The mixotrophs have both the genetic and physiological capacity to oxidize Fe(II), while the heterotrophs induce chemodenitrification (abiotic denitrification) through reactive nitrogen intermediates (e.g., NO_2_⁻, NO). Although both autotrophic and mixotrophic (enzymatic) nitrate-reducing Fe(II) oxidation have been investigated since, mostly driven by environmental and wastewater-related interests, fundamental understanding of this process is still limited. The diversity and detailed metabolic mechanisms of dissimilatory Fe(II) oxidation coupled to nitrate reduction remain largely unknown.

Typically, model microorganisms are used to elucidate such pathways. However, identifying Fe(II) oxidation-specific proteins in mixotrophic cultures is difficult, as electron flow originates from both organic substrates and Fe(II). Additionally, distinguishing between biotic and abiotic nitrate reduction complicates electron balancing and metabolic analysis.

A more tractable approach is to study a pure culture of an autotrophic NRFeOx bacterium, and later extend insights to more complex systems. Unfortunately, to our knowledge, no such pure culture exists that can be stably maintained using solely Fe(II) and nitrate as energy substrates over multiple transfers. Nevertheless, four autotrophic nitrate-reducing Fe(II)-oxidizing enrichment cultures—KS, BP, AG and HP— are currently available, the first three have been well-characterized both physiologically and genomically (Straub *et al*. 1996, Huang, Straub, Blackwell *et al*. 2021, Huang, Straub, Kappler *et al*. 2021, Jakus *et al*. 2021). All are dominated by autotrophic Fe(II)-oxidizers of the genus *Ferrigenium* (Betaproteobacterium), specifically *Ferrigenium straubiae* strain KS (*F. straubiae*), *Candidatus Ferrigenium bremense* strain BP, and *Candidatus Ferrigenium altingense* strain AG (Becker and Kappler 2025, 2025). Among these, culture KS represents the oldest and best-studied autotrophic NRFeOx enrichment culture to date. Further it is the only culture from which the main Fe(II) oxidizer has been successfully isolated (*Ferrigenium straubiae* strain KS: KCTC 25982; = DSM 118991) (Becker and Kappler 2025). Although this culture is composed of up to 98% *F. straubiae* under Fe(II)-oxidizing, denitrifying conditions (Tominski *et al*. 2018), the *F. straubiae* isolate itself grows only under Fe(II)-oxidizing, microoxic conditions (Becker and Kappler 2025). Given its high abundance in the NRFeOx enrichment and the availability of an isolate capable of Fe(II) oxidation under defined conditions, *F. straubiae* provides a unique opportunity for comparative transcriptomic analysis.

The first comprehensive analysis of the KS genome was conducted by He *et al*., followed by comparative transcriptomic studies by Huang *et al*., which aimed to elucidate the role of the flanking community (He *et al*. 2017, Huang, Straub, Blackwell *et al*. 2021). Their work raised several key questions regarding the physiology and metabolism of *F. straubiae*. What are the autotrophic CO_2_ fixation pathways? Which of the putative Fe(II) oxidases are functionally prioritized? How are electrons transferred from the Fe(II) oxidase to the quinone pool, particularly across the periplasmic space? Does *F. straubiae* possess a non-canonical NO reductase gene for using NO energetically and to overcome potential NO toxicity? What role does the flanking community play in supporting autotrophic NRFeOx activity? On what basis was it assumed that *F. straubiae* could utilize oxygen as a terminal electron acceptor under anoxic conditions, and why was ‘dark oxygen’ production invoked?

Since these studies were conducted, new findings have emerged regarding unusual CO_2_ fixation pathways, NO dismutase and reductase variants, and Fe(II)-oxidizing bacteria in general. In parallel, bioinformatic tools have advanced rapidly, creating an opportunity to gain new insights through a comprehensive reanalysis of the KS genome.

In this study, we investigated these fundamental questions by conducting the first comparative transcriptomic analysis of an isolate of a dominant Fe(II)-oxidizing bacterium from an autotrophic NRFeOx enrichment culture, i.e. *F. straubiae* (Figure 1).

**Figure 1:**
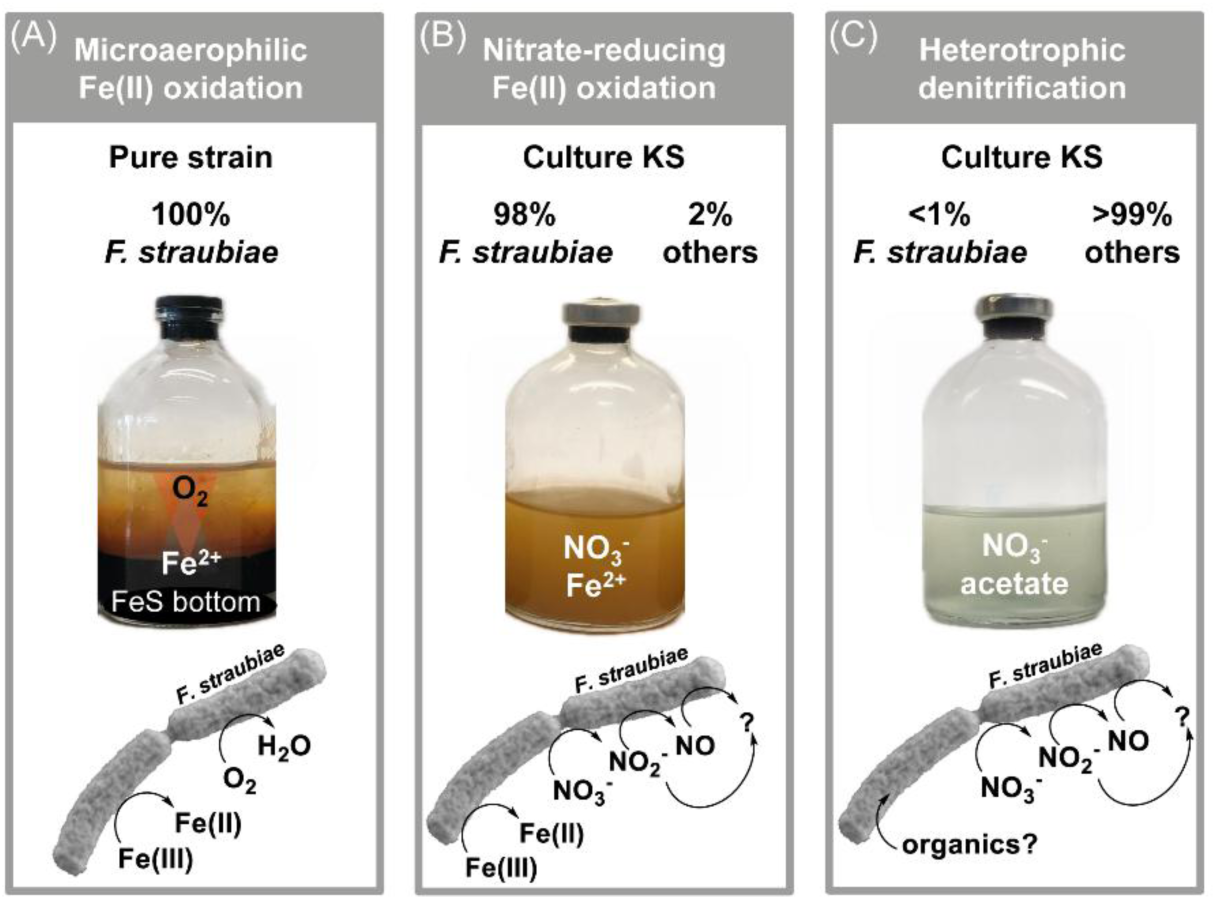
Comparative transcriptomic study using *F. straubiae* as an isolate under microaerophilic Fe(II) oxidation (A) compared to denitrifying conditions using culture KS (B) and (C). In this study we cultured condition (A) and (B) at 20°C [extracted and analyzed transcriptomes] and reanalyzed transcriptomic data of Huang *et al*. of condition (B) and (C) at 28°C (Huang, Straub, Blackwell *et al*. 2021).

Using recently developed bioinformatic tools, we evaluate the potential functions of newly identified proteins. While previous studies primarily focused on known proteins, our approach emphasizes uncharacterized proteins, as modern structural prediction tools have become more accessible and reliable, allowing us to derive valuable insights into their potential functions. Our findings place a spotlight on key candidate proteins that may play central roles in microbial nitrate-reducing Fe(II) oxidation.

## 5. Methods

### 5.1 Source of microorganism and medium preparation

Culture KS, isolated by Straub *et al*., has been maintained under autotrophic growth conditions in our laboratory for over two decades (Straub *et al*. 1996). Transfers are routinely performed with a 1% (v/v) inoculum into an anoxic, unfiltered mineral medium buffered with 22 mM bicarbonate and supplemented with 10 mM FeCl_2_ and 4 mM NaNO_3_. The final pH was adjusted to 6.9–7.2. Cultures were incubated at 28°C in the dark without agitation, with a headspace gas mixture of N_2_/CO_2_ (90:10). The 22 mM bicarbonate-buffered mineral medium was prepared according to Hegler *et al*. (Hegler *et al*. 2008), and anoxic media preparation according to Widdel and Bak (Widdel and Bak 1992). 7-vitamin solution, SL-10 trace elements, selenite-tungstate solution (11 µM Na_2_SeO_3_ and 12 µM Na_2_WO_4_), were prepared as 1000× stock solutions as described by Widdel and Bak (1992).

The strain *Ferrigenium straubiae* KS (hereafter referred to as *F. straubiae*) was obtained from culture KS, isolated and maintained as an active culture using agarose-stabilized Fe(II)-O_2_ gradient tubes [following the approach described by Becker and Kappler, find details in corresponding SI], over 3 years (Becker and Kappler 2025). For the transcriptomic study, we prepared 9 agarose-stabilized Fe(II)-O_2_ gradient cultures in serum bottles (gradient serum bottles), which were a scaled-up version of our standard gradient tubes, to obtain enough biomass and nucleic acids for the transcriptomic analysis. Each of these gradient serum bottles contained a 20-mL FeS plug, a 50-mL top layer, and a 30-mL headspace and was cultivated at 20°C.

### 5.2 Transcriptomics

#### Experimental design

To identify enzymatic pathways specific to autotrophic nitrate-reducing Fe(II) oxidation, we analyzed transcriptomes under four conditions. Please note that the corresponding raw data of the first two conditions were collected by Stefanie Becker, while the last two steam from Yu-Ming Huang and were collected and analyzed for a previous study of our laboratory (Huang, Straub, Blackwell *et al*. 2021). *F. straubiae* with NRFeOx flanking community = culture KS.

- **autotrophic microoxic Fe(II)-oxidizing conditions at 20°C (A_microoxic)**: Pure culture of *F. straubiae* grown in Fe(II)/O_2_ opposing gradient serum bottles
- **autotrophic denitrifying Fe(II)-oxidizing conditions at 20°C (A_denit_20°C)**: *F. straubiae* with NRFeOx flanking community grown with 10 mM Fe(II) and 4 mM nitrate
- **autotrophic denitrifying Fe(II)-oxidizing conditions at 28°C (A_denit_28°C)**: *F. straubiae* with NRFeOx flanking community grown with 10 mM Fe(II) and 4 mM nitrate
- **heterotrophic denitrifying conditions at 28°C (H_denit)**: *F. straubiae* with NRFeOx flanking community grown with 2 mM acetate and 4 mM nitrate

In contrast to Huang *et al*. (2021), the raw data (steaming from Yu Ming Huang and from us) were here processed by mapping the transcripts solely to the metagenome-assembled genome of *F. straubiae* KS instead of the whole metagenome, which improved the resolution for interpreting gene expression in *F. straubiae*.

#### Cultivation and biomass sampling

The **A_microox setup** was carried out using *F. straubiae* inoculated into six gradient serum bottles (Figure S1). Cultures were incubated for one month to obtain a high cell density (6.5 × 10⁶ cells mL⁻¹; cell viability was checked by fluorescence microscopy; exemplary micrographs are shown in Figure S3). The FeS layer at the bottom of the tubes provided a continuous source of Fe(II), making the gradient culture system more similar to a continuous than a batch cultivation method.

The **A_denit_20°C setup** was carried out using culture KS inoculated into twelve 100-mL serum bottles containing 50 mL liquid mineral medium. Cultures were harvested during the exponential growth phase (after 2.7 days), at that timepoint with 8.0 mM Fe(II) and 3.7 mM nitrate remaining in the cultures, and a cell density of 1.25 × 10⁷ cells mL⁻¹.

The **A_denit_28°C setup** was performed by Huang *et al*. (2021) in the same laboratory under identical conditions, except for the higher incubation temperature (8°C above the A_denit_20°C setup). Samples were collected on day 2, at that timepoint with 3.7 mM Fe(II) and 3.8 mM nitrate remaining in the cultures, and 7.95 × 10⁶ cells mL⁻¹.

The **H_denit setup** was performed by Huang *et al*. (2021). Samples were collected on day 5, at that timepoint with 3.68 mM acetate, 2.84 mM nitrate and 0.04 mM nitrite remaining in the cultures, and 5.29 × 10⁷ cells mL⁻¹. Prior to inoculation, the culture had been transferred twice in medium lacking 10 mM FeCl_2_, corresponding to approximately seven generations on heterotrophic medium.

In contrast to Huang *et al*. (2021) [setups: A_denit_28°C and H_denit], who harvested biomass by filtration and proceeded directly with RNA extraction, we harvested cells by centrifugation at 7,197 × *g* for 10 min at 25°C, and the resulting pellets were flash-frozen in liquid nitrogen and stored at –80 °C until further processing.

#### RNA extraction

RNA extraction for all setups was performed according to the Lueders *et al*. (2004) protocol with minor modifications (Lueders, Manefield, and Friedrich 2004). Both our study and that of Huang *et al*. (2021) [setups A_denit_28°C and H_denit] applied this method. In the latter study, cells were lysed by combining two 2 mL MP Biomedicals™ Lysing Matrix E reaction tubes into a single 15 mL Falcon tube containing filter pieces and collected biomass. Then, the tube was vortexed (see the original publication for details).

Each A_microox serum bottle yielded a 5 g pellet consisting of agar, Fe(III) minerals, and biomass (Figure S1). Following the manufacturer’s recommendations, we used MP Biomedicals™ TeenPrep™ Lysing Matrix E 15-mL reaction tubes for cell lysis. For the A_denit_20°C cultures, which produced approximately 0.25 g pellets, we used the smaller MP Biomedicals™ Lysing Matrix E 2-mL reaction tubes.

To disrupt the cells, we added 7.50 mL of chilled PB buffer (112.87 mM Na_2_HPO₄ and 7.12 mM NaH_2_PO₄) and 2.5 mL of TNS buffer (500 mM Tris-HCl, 100 mM NaCl, and 10% [wt/vol] SDS) to each TeenPrep™ sample, and 10-times less to the 2-mL reaction tube preps. Samples were bead-beaten twice for 30 s at 4.0 m/s using a FastPrep-24™ Classic instrument (MP Biomedicals™), with a 5 min cooling step on ice between runs.

All centrifugation steps were performed at 7,197 × g and 4°C. All sample transfers were conducted on ice. After an initial centrifugation for 10 min, the supernatant was distributed into sterile 2-mL reaction tubes (1 mL per tube) and subjected to phenol-chloroform-isoamyl alcohol and chloroform-isoamyl alcohol extractions, as described by Lueders *et al*. (2004). The aqueous phases corresponding to one culture were then combined in a new 15 mL tube and precipitated overnight using PEG solution (30% PEG 6000 [wt/vol] and 1.6 M NaCl) with a ratio of one-part sample and two-parts PEG solution.

From the ethanol washing step onward, all procedures were conducted under a clean bench. The ethanol was carefully removed with a filtered pipette tip, and the resulting pellet was air-dried at room temperature for approximately 10 minutes. The DNA/RNA pellets were dissolved in 35 μL of TE buffer solution (10 mM Tris-HCl and 1 mM EDTA at pH 7.4). The pellets corresponding to two cultures were then pooled and frozen at −80°C. The A_denit_20°C pellets were centrifuged after pooling the samples due to Fe(III) mineral contamination (Figure S1). The pellets were centrifuged at 20,817×g for 1 min. The supernatant was transferred to a fresh reaction tube.

#### RNA sequencing, mapping, and differential RNA abundance

RNA samples were further processed using the Zymo RNA Clean & Concentrator Kit following the manufacturer’s protocol for rigorous treatment. After that, RNA was quantified with a Qubit RNA HS Assay Kit (Thermo Fisher) and RNA integrity was checked by Agilent 2100 BioAnalyzer with RNA 6000 Pico kit (Agilent). Only two samples of A_microox and A_denit_20°C passed the quality control and were used for library preparation, which was performed with Illumina Stranded Total RNA Prep, Ligation with Ribo-Zero Plus Microbiome kit according to the manufacturer’s instructions. In brief, 100 ng of total RNA per sample were subjected to rRNA depletion, followed by cDNA library construction, adapter ligation and 17 cycles of barcoding PCR. Obtained libraries were quantified with Qubit 1x DNA HS Assay Kit (Thermo Fisher) and the fragment distribution was checked with the Agilent 2100 BioAnalyzer using High Sensitivity DNA Kit (Agilent). Libraries were subsequently pooled and sequenced on the Illumina MiSeq device using MiSeq Reagent Kit v3 (150 cycles) with a run mode 74,10,10,74. Data was demultiplexed using nf-core/demultiplex v1.4.1 (Ewels P *et al*. 2022) of the nf-core collection of workflows (Ewels PA *et al*. 2020), executed with Nextflow v24.10.2 (Di Tommaso *et al*. 2017) and utilizing reproducible software environments from the Bioconda (Grüning *et al*. 2018) and Biocontainers (Veiga Leprevost da *et al*. 2017) projects. 3,472,675 to 5,999,243 read pairs were generated for four samples (total 19,082,590).

The four samples sequenced in this study plus one heterotrophic (SRX9677098) and three autotrophic (SRX9677095, SRX9677096, SRX9677097) samples of Huang *et al*. (2021) (NCBI bioproject PRJNA682552, https://www.ncbi.nlm.nih.gov/bioproject/PRJNA682552) were included in the further processing (Huang, Straub, Blackwell *et al*. 2021).

Read quality control and mapping to the metagenome-assembled genome of *F. straubiae* KS and its annotation (IMG Taxon ID: 2878407288) was carried out using the pipeline nf-core/rnaseq v3.15.1 (Patel *et al*. 2024) of the nf-core collection (Ewels PA *et al*. 2020) with Nextflow v24.04.4 (Di Tommaso *et al*. 2017). Briefly, quality was assessed with FastQC v0.12.1 (Andrews [2017] 2010), reads trimmed by Trim Galore! v0.6.7 (Krueger [2017] 2021), rRNA removed by SortMeRNA v4.3.6 (Kopylova, Noé, and Touzet 2012) with the SILVA database (Quast *et al*. 2013), >80% reads mapped with STAR v2.7.10a (Dobin *et al*. 2013), and finally transcripts were quantified with RSEM v1.3.1 (Li B and Dewey 2011). More than 93% of all annotated CDS were detected (>0 TPM) per sample, however, the sample with treatment heterotrophic denitrification at 28°C (SRX9677098) of Huang *et al*. (2021) had only 49% detected (Huang, Straub, Blackwell *et al*. 2021). Differential gene expression analysis was conducted using qbic-pipelines/rnadeseq v2.4 (https://doi.org/10.5281/zenodo.12799992) (Wacker *et al*. 2024), which is based on nf-core template (Ewels PA *et al*. 2020). For differential expression analysis, the read quantification data from RSEM were processed with DESeq2 v1.40.2 (Love, Huber, and Anders 2014). The thresholds for differentially expressed genes were 2 for the log2 Fold Change and 0.05 for the adjusted *P*-value.

#### Quality of the Transcriptomic data

We note that the transcriptomic data are of limited quality due to the small number of samples, and therefore all transcriptomic results should be interpreted with caution. Nevertheless, the statistically significant expression patterns proved valuable for identifying novel proteins potentially involved in key metabolic processes under the respective growth conditions. Thus, the transcriptomic data were used in conjunction with our bioinformatic genome analysis to support the identification and functional interpretation of relevant proteins. Further, we would like to mention that the heterotrophic condition (setup) does not equal with the heterotrophic growth of *F. straubiae*, as it is presumably an obligate chemolithoautotrophic bacterium (Becker and Kappler 2025).

### 5.3 Bioinformatic analysis of genome and proteins

To better understand the activity of canonical metabolic pathways in *F. straubiae*, we linked transcriptomic data to KEGG annotations. This allowed us to perform pathway-level analyses on all genes assigned to the KEGG database.

In addition, we aimed to identify unclassified but potentially relevant proteins involved in Fe(II)-oxidizing metabolism by reanalyzing the *F. straubiae* genome for multicopper oxidases, multiheme cytochromes, and other Fe(II)-associated genes, with particular focus on their expression patterns. To functionally characterize targets that were highly expressed under any of the tested conditions, we analyzed their amino acid sequences for conserved motifs, known functional features, and evolutionarily coupled residues. We predicted three-dimensional structures using state-of-the-art modeling approaches, which enabled us to perform protein structure-based searches.

Genes encoding proteins of interest were frequently found to be co-localized, suggesting that they may form functional units. Therefore, we examined genomic neighborhoods and compared the corresponding transcriptomic profiles to support potential operon- or pathway-level organization.

**Multi-copper oxidase screening** was performed by the use of a custom python script, coded by Austin Chambers, Christopher Blanda, and Rene Hoover. It searches for specific protein sequence motifs as described in Gräff *et al*. (Gräff *et al*. 2020).

**Fe-related and potentially related novel protein screening** was performed by FeGenie and MHCScan (Garber, Nealson, and Merino 2024) using the Middle Author Bioinformatics server (omix.midauthorbio.com). MHCScan searches for the same genes as FeGenie (Garber *et al*. 2020) and additionally for clusters of multi heme cytochromes translocated to the periplasmic space.

**Protein homology analysis** was carried out by phylogenetic tree generated by Geneious Prime v2019.2.3 using Muscle (Edgar 2022).

**Protein 3D structure prediction** was performed using MODELLER v10.0 (Sali *et al*. 1995), I-TASSER v5.1(Zheng W *et al*. 2021, Zhou X *et al*. 2022, Zheng W *et al*. 2025), and AlphaFold 3 (Abramson *et al*. 2024) which were run via the MPI Bioinformatics Toolkit server (Zimmermann *et al*. 2018, Gabler *et al*. 2020), I-TASSER and AlphaFold dedicated servers, respectively. However, only the AlphaFold-derived structures are presented here. MODELLER and I-TASSER were used for comparative purposes, and in all cases, the models generated by the different tools were highly similar within the scope of this study.

**Signal peptide identification and cleavage sites** were predicted by SignalP v6.0 (Teufel *et al*. 2022, Nielsen *et al*. 2024).

**Transmembrane helices identification** were predicted by DeepTMHMM - 1.0 (Hallgren *et al*. 2022).

**Protein function prediction** was performed by **sequence-based database searching** using HHpred (Hildebrand *et al*. 2009) on the MPI Bioinformatics Toolkit server and by **structure-based database searching** using FoldSeek (Kempen van *et al*. 2024) on the Foldseek Search Server. Further we searched for protein signatures (motifs) and co-factors. **Protein sequence motifs** (InterPro’s member database signatures) were identified by InterProScan run on the EMBL-EBI server (Jones *et al*. 2014, Blum *et al*. 2025). **Co-factors and binding sites** were predicted by COFACTOR using the I-TASSER server (Roy, Yang, and Zhang 2012). Searches were performed within January to October 2025. Potential variations in substrate specify was predicted from **evolutionary sequence covariations** by searching for evolutionary coupled amino acids using EVcouplings via its respective server (https://v2.evcouplings.org/ ) (Hopf *et al*. 2019).

**Functional units** (proteins that share a common function) were predicted by evaluating co-expression patterns based on transcriptome data, in combination with **genomic neighborhood analysis** using cblaster (Gilchrist *et al*. 2021) and clinker (Gilchrist and Chooi 2021) searches. These analyses were used to assess whether genes are located in proximity by chance or if conserved gene clusters in other species indicate that they form a functional unit. Both cblaster and clinker search were performed using the CompArative GEne Cluster Analysis Toolbox (CAGECAT) server (Release: 1.0).

**Gene neighborhoods** were initially analyzed using the metagenome-assembled genome of *F. straubiae* KS (IMG Taxon ID: 2878407288). Subsequently, a genome derived from a pure isolate of *F. straubiae* KS was published (Taxon ID: 8132338324) (Becker and Kappler 2025). Assemblies from pure isolates are considered more accurate; therefore, all gene neighborhood analyses were validated against the isolate genome as well. No differences were observed.

## 6. Results and Discussion

The results and discussion section issue first CO_2_ fixation, using Fe(II) as the primary electron donor supporting autotrophic growth, followed by an analysis of the generation, regeneration and utilization of its key substrates—CO_2_, NADPH, ferredoxin, and ATP (section 6.11). We then trace the electron flow from Fe(II) oxidation (section 6.2), across the periplasm, into the quinone pool (sections 6.3 and 6.4), and finally towards the terminal electron acceptors (nitrogen species and oxygen [sections 6.5 and 6.6]). Finally, we address iron-related genes involved in transcription regulation, iron acquisition and storage (section 6.7).

This framework allowed us to explore how energy and quinone demands can be met under both nitrate-reducing and microaerophilic Fe(II)-oxidizing conditions. A Volcano plot based on the most important comparison [autotrophic nitrate-reducing Fe(II) oxidation (20°C) verses autotrophic microaerophilic Fe(II) oxidation (20°C)] to identify differences in O_2_ respiration and dentification under autotrophic conditions can be found in the supplemental information Figure S1. By seven subplots (within Figure S2) we highlighted all genes which were analyzed in the upon mentioned sections.

### 6.1 Autotrophic CO_2_ fixation pathways in *F. straubiae*

Seven autotrophic CO_2_ fixation pathways have been described in autotrophic microorganisms; the reductive pentose phosphate cycle (Calvin-Benson-Bassham; CBB cycle); the reverse tricarboxylic acid (rTCA) cycle including a variant—reversed oxidative TCA cycle (roTCA)—, the 3-hydroxypropionate (3HP) bi-cycle, the 4-hydroxybutyrate/3-hydroxypropionate (4HB/3HP) cycle, the dicarboxylate/4-hydroxybutyrate (DC/4HB) cycle, the reductive acetyl-CoA (Wood-Ljungdahl) pathway, and the reductive glycine pathway (Bar-Even, Noor, and Milo 2012, Mall *et al*. 2018, Sánchez-Andrea *et al*. 2020, Steffens *et al*. 2021, Mitic *et al*. 2023). Metagenomic analysis revealed that *F. straubiae* employs a non-canonical CBB cycle, has the genetic potential for a roTCA cycle (rTCA variant), and has an incomplete reductive glycine pathway (Figure 2).

**Figure 2:**
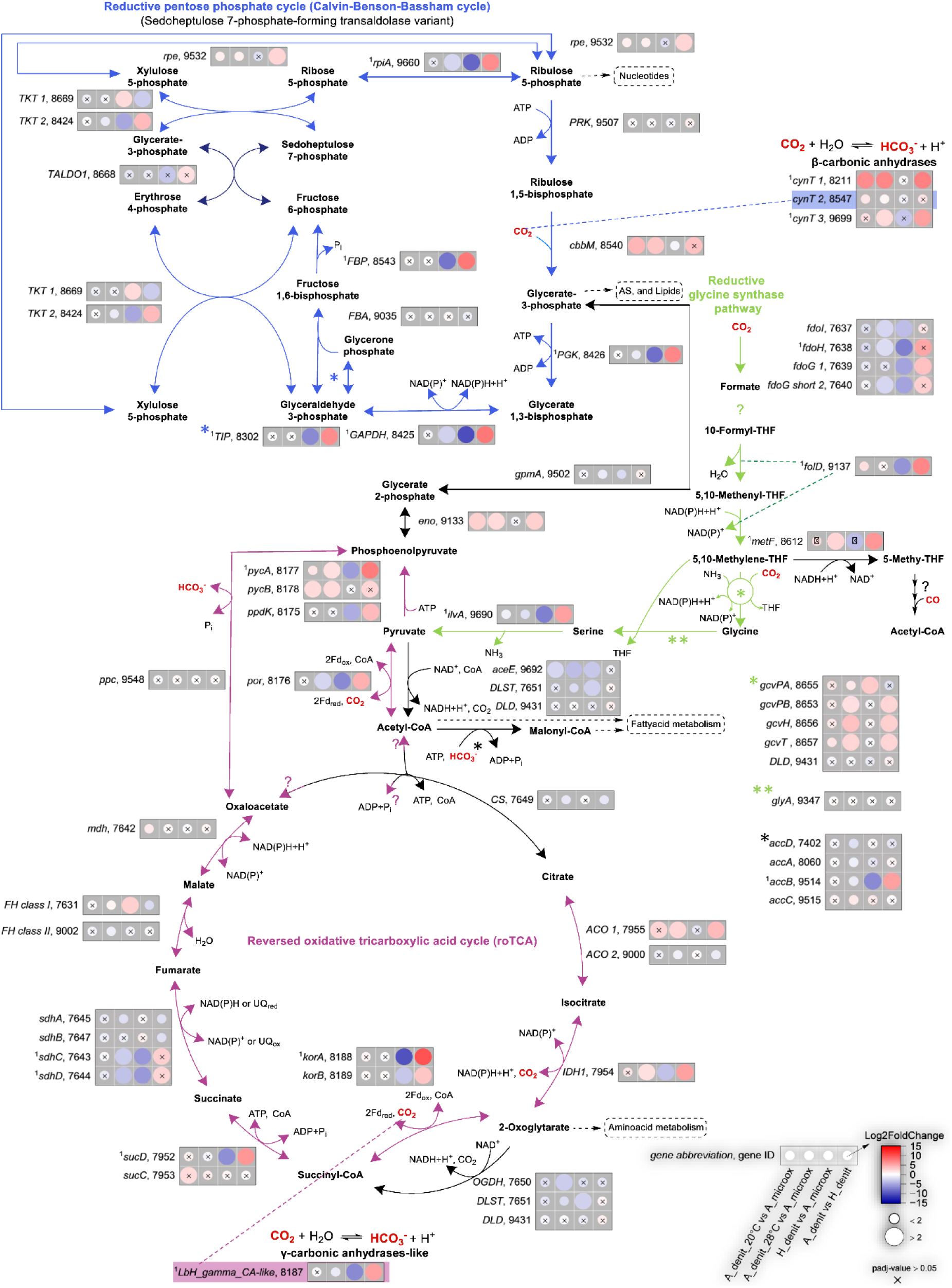
Autotrophic CO_2_ fixation gene expression in *F. straubiae*. CBB cycle (blue, the characteristic transaldolase variant reaction is indicated by dark blue) and reductive oxidative TCA cycle (purple) are complete pathways. Question marks at the arrows indicate that the reverse reaction lacks experimental evidence and thus is hypothetical. The glycine synthase pathway (green) is indicated, however, based on current knowledge it is incomplete as the formate tetrahydrofolate ligase is not encoded in the genome of *F. straubiae*. The log_2_ fold changes of normalized transcript count (Log2FoldChange) are shown as colored dots. The labels follow the structure: gene name, last four numbers of IMG Gene ID. All IMG Gene IDs share the prefix 287840XXXX. Connected rows indicate several subunits of an enzyme complex while not connected bubble-plots correspond to independent enzymes which, in some cases, catalyze the same reaction. The four conditions were: autotrophic denitrifying Fe(II) oxidation at 28°C (A_denit_28°C, n=3), heterotrophic denitrification at 28°C (H_denit, n=1), autotrophic denitrifying Fe(II) oxidation at 20°C (A_denit_20°C, n=2) and autotrophic microaerophilic Fe(II) oxidation at 20°C (A_microox, n=2). *F. straubiae* was grown as community member of culture KS in all nitrate containing conditions. We found paralogues for *ACO*s and *TKT*s and named them *ACO 1*, *ACO* 2, *TKT* 1 and *TKT* 2. Asterisks have been used when there was no space for the corresponding transcripts of a reaction. ^1^no transcript detected in H_denit. Gene abbreviation in table S7.

#### 6.1.1 CBB cycle—RubisCO is most dominant under denitrifying-autotrophic conditions

Literature describing chemolithoautotrophic, RubisCO-mediated CO_2_ fixation is limited (Frolov *et al*. 2019, Nishihara, Kato, and Ohkuma 2025). Accordingly, we did not find previous reports of the specific modification of the CBB cycle observed in *F. straubiae*, which we identify here as a form II RubisCO-mediated transaldolase variant (specifically, a sedoheptulose-7-phosphate-forming transaldolase variant) (Figure 2).

Unlike the canonical CBB cycle, this variant bypasses the requirement for fructose-1,6-bisphosphate aldolase by converting fructose-6-phosphate and erythrose-4-phosphate into glycerate-3-phosphate and sedoheptulose-7-phosphate (Figure 2, dark blue reaction). Explanatory illustrations of Cbb cycle variants, including stoichiometric considerations, are provided by both Ohta and Hundson (Ohta 2022, Hudson 2024).

Consistent with the microaerophilic lifestyle of *F. straubiae*, its RubisCO was identified as a bacterial Form II (gene: *cbbM*; IMG ID: 2878408540), which is commonly associated with microaerophilic bacteria (He *et al*. 2016a). The *cbbM* gene encodes the large subunit of the homomultimeric Form II RubisCO, whereas other forms additionally include a small subunit (Tabita *et al*. 2008, Varaljay *et al*. 2016). In contrast to Form I RubisCO, Form II possess lower affinity for CO_2_ and are not optimized for functioning under high O_2_ partial pressures (Badger and Bek 2008). As a result, organisms that rely solely on Form II RubisCO are typically strictly microaerophilic or anaerobic, where oxygen levels are sufficiently low to minimize the oxygenase side reaction. In agreement with this pattern, Form II RubisCO is also present in other microaerophilic and anaerobic Fe(II)-oxidizing bacteria, including *Sideroxydans lithotrophicus* ES-1, *Gallionella capsiferriformans* ES-2, *Acidithiobacillus ferrooxidans*, *Mariprofundus ferrooxydans* PV-1, and *Thiobacillus denitrificans* (Beller *et al*. 2006, Emerson *et al*. 2013).

RubisCO and phosphoribulokinase (PRK) are unique to the CBB cycle (key enzymes), whereas the other CBB enzymes also function in the pentose phosphate pathway or glycolysis. Therefore, expression of this key enzymes strongly indicates autotrophic CO_2_ fixation. Notably, RubisCO and PRK transcripts were detected under all conditions tested (Figure 2), suggesting that growth under "*heterotrophic*" conditions does not necessarily depend solely on heterotrophic carbon assimilation metabolism in *F. straubiae*. Despite the constitutive expression of cbbM under all conditions, the highest transcript levels were observed under denitrifying-autotrophic conditions (SI-Table 3). Interestingly no significant difference was observed between the microoxic-autotrophic and heterotrophic conditions.

This underscores the importance of RubisCO, which catalyzes the initial fixation of CO_2_ into organic molecules and thus acts as a key control point for autotrophic growth in *F. straubiae*. CO_2_ fixation is energetically costly, and RubisCO’s kinetic limitations necessitates high cellular abundance of this enzyme, which adds the burden of additional protein synthesis. Thus, the control of *cbb* gene expression is vital to balance the overall cost of operating the CBB cycle with the cell’s energetic state. In bacteria, this regulation is commonly mediated by the LysR-type transcription factor CbbR. In *F. straubiae* the gene encoding CbbR (IMG ID: 2878408539) is located directly upstream of its only *cbb* operon, in the opposite orientation (Figure 3A, which aligns with the common observation in bacteria (Dangel and Tabita 2015).

**Figure 3:**
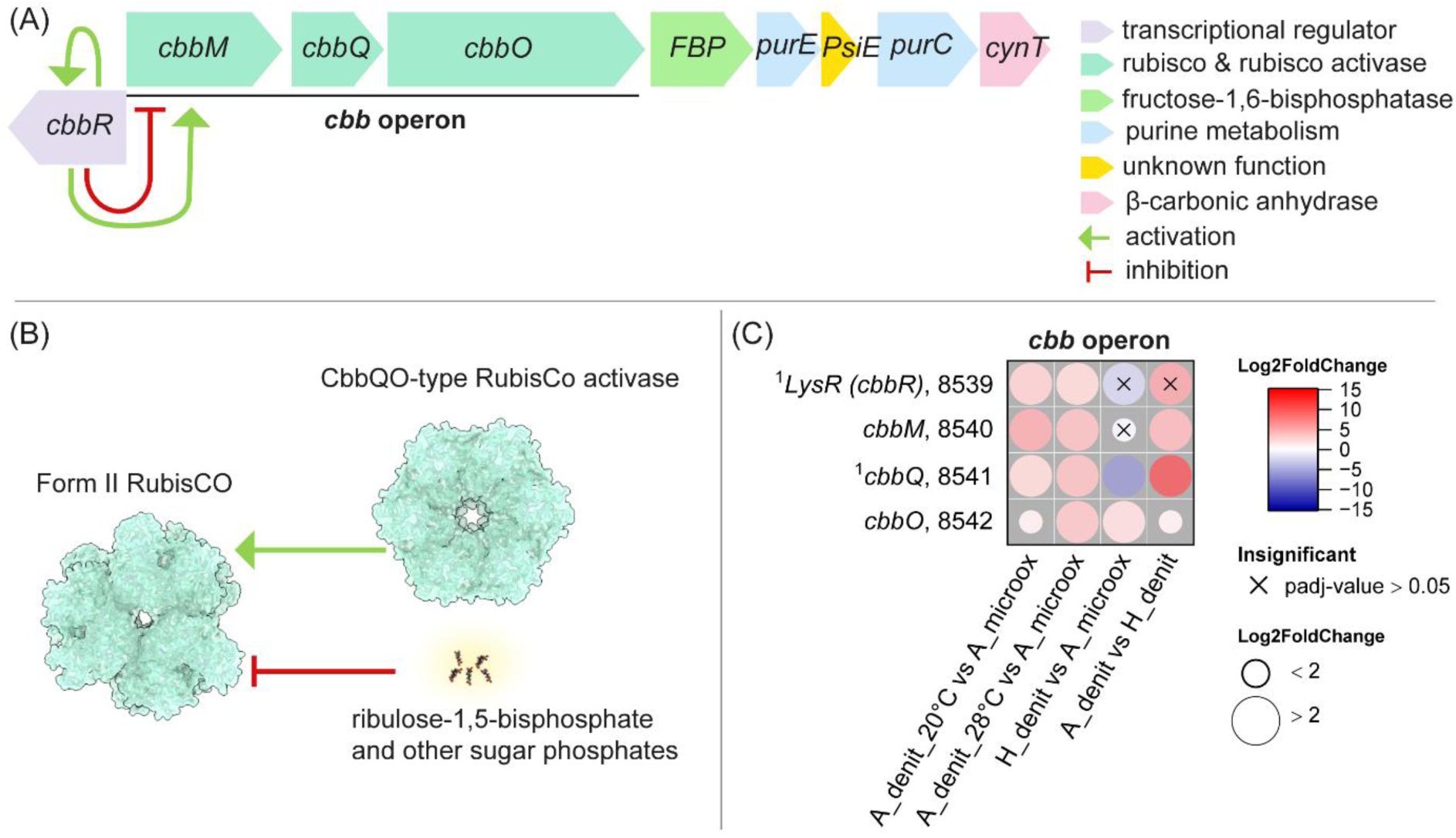
(A) CBB cycle regulation on the transcriptional level by CbbR (the product of *cbbR*). CbbR is a transcriptional regulator which interacts with other regulatory factors causing either induction (green arrow) or suppression (red symbol) of target genes. (B) RubisCO regulation on the protein level by sugar phosphate (inactivation), and by the CbbQO complex due to reactivation. The RubisCO complex is represented by the *Gallionellaceae* sp. Form II RubisCO (https://doi.org/10.2210/pdb5C2C/pdb). The CbbQO complex is represented by the *Acidithiobacillus ferrooxidans* CbbQO-type Rubisco activase (https://doi.org/10.2210/pdb6L1Q/pdb). (C) The fold change of normalized counts (Log2FoldChange) of *cbb* operon and *cbbR* transcripts. Gene abbreviation in able S7.

Further the *cbb* operon of *F. straubiae* has a very classical structure of cbbM (encoding Form II RubisCO) followed by cbbQ (IMG ID: 2878408541), and cbbO (IMG ID: 2878408542) which cluster with other *cbb* metabolic genes [Table 3 of Badger and Bek] (Badger and Bek 2008). The genes identified here as cbbRMQO were not annotated as those by the IMG pipeline, for details see SI-Text 2. CbbR is the "*master*" regulator playing an essential role in *cbb* gene transcription (Figure 3A) while CbbQ and CbbO regulate the CBB cycle on the protein level (Figure 3B) (Dangel and Tabita 2015, Böhnke and Perner 2017). CbbR is a LysR-type transcription factor and as such, it can act both as an activator and repressor of *cbb* genes [Maddocks and Oyston plus references therein] (Maddocks and Oyston 2008), and autoinduces its own transcription (Axler-DiPerte, Miller, and Darwin 2006). Therefore, and due to its interaction with other transcriptional regulators, LysR-mediated regulation can be highly complex (Joshi *et al*. 2013, Dangel, Luther, and Tabita 2014). Transcripts for CbbR could not be detected under heterotrophic conditions, and transcript abundance under autotrophic conditions was low; thus, the differences between conditions were not statistically significant (*P* > 0.05, Figure 2: RubisCO operon).

On the protein level, RubisCO is inhibited by its substrate RuBP and other sugar phosphates (Tsai *et al*., 2015). Removal of these inhibitors from the active site is essential to proceed the CBB cycle. Its reactivation is facilitated by CbbQ (an AAA+ ATPase) and CbbO (a von Willebrand factor type A domain-containing protein)(Tsai *et al*. 2015). CbbQ and CbbO form a RubisCO activase complex (Figure 3B) and thus the presence of both are required for activation of RubisCO form II (Tsai *et al*. 2020). However, a significant difference between both autotrophic and the heterotrophic condition was observed in the expression of cbbQ, which was found to be upregulated under both microoxic and nitrate-reducing autotrophic conditions and not detected under heterotrophic conditions (Figure 3C, SI-Table 3). Expression levels were found to be marginally elevated in the denitrifying-autotrophic conditions in comparison to the microoxic-autotrophic condition.

The CbbO transcript was detected under all conditions, but was significantly higher under denitrifying conditions compared to the microoxic-autotrophic conditions.

In conclusion, the CBB cycle plays a significant role in CO_2_ fixation under autotrophic conditions. Notably, RubisCO was expressed at higher levels under denitrifying-autotrophic conditions, possibly indicating the presence of a condition-specific factor that affects its activity and requires increased enzyme abundance compared to microoxic-autotrophic conditions. This finding is counterintuitive, as one would generally expect higher RubisCO levels under conditions where oxygen is present [as both O_2_ and CO_2_ compete for the active site of RubisCO].

Furthermore, RubisCO was constitutively expressed, and transcripts were detected under heterotrophic conditions. However, our data suggest that RubisCO may have been less effectively reactivated by the CbbQO activase complex under these conditions. Nevertheless, the expression pattern indicates that the autotrophic metabolism remains important in *F. straubiae* even in the presence of acetate.

Given that acetate is not a growth-supporting carbon source for *F. straubiae* (Becker and Kappler, 2025), two possibilities must be considered: either the flanking community feed *F. straubiae* with other organic compounds, or *F. straubiae* relies primarily on its autotrophic metabolism even under heterotrophic conditions.

Notably, we identified two transketolase paralogs (TKT1 and TKT2) with contrasting expression profiles: TKT2 was upregulated under autotrophic conditions, whereas TKT1 was upregulated during heterotrophic conditions. This reciprocal pattern suggests functional divergence, with TKT2 likely supporting CO_2_ fixation through the CBB cycle, and TKT1 functioning within the pentose phosphate pathway when organic carbon is available.

#### 6.1.2 Reversed oxidative tricarboxylic acid (roTCA) cycle does not function as a complete cycle under autotrophic conditions

In addition to the CBB cycle, *F. straubiae* possesses the genetic potential of a roTCA cycle which is metabolically connected via 2,3-bisphosphoglycerate-dependent phosphoglycerate mutase (GpmA) and enolase (Eno) (Figure 2). The reverse TCA cycle (rTCA) is conventionally considered as incomplete in absence of the ATP citrate lyase, an enzyme that cleaves citrate into acetyl-CoA and oxaloacetate—an essential step in the canonical reversed direction of TCA cycle. Indeed, the *F. straubiae* genome lacks the ATP citrate lyase. However, it has been demonstrated by Mall *et al*. (2018) that citrate synthase (CS)—an enzyme typically associated with the forward (oxidative) TCA cycle— can, under specific conditions, facilitate the reverse direction resulting in a reversed oxidative TCA cycle. Although citrate synthase exhibits unfavorable kinetics in the reverse direction, previous studies suggest that under high CO_2_ concentrations and elevated levels of citrate synthase expression (>7% of total protein), citrate cleavage in the reverse direction may be feasible (Steffens *et al*. 2021)*et al*.. Additionally, the presence of 2-oxoglutarate:ferredoxin oxidoreductase (OGOR) further supports the potential for reverse flux. OGOR is a thiamine pyrophosphate and [4Fe-4S] cluster-dependent enzyme from the reductive TCA cycle. It is a functional orthologue of NAD-dependent dehydrogenase from the TCA cycle, however it can catalyze both the oxidation of 2-oxoglutarate to succinyl-CoA and, in the reverse direction, fixes CO_2_ to succinyl-CoA, forming 2-oxoglutarate and CoA.

*F. straubiae* possesses the alpha and beta subunits of OGOR which form a heterotetrameric (αβ)_2_ complex based on structural analysis of different species (Chen *et al*. 2019) (corresponding genes are *korA* and *korB* [alpha-**k**etoglutarate **o**xido**r**educase]).

Therefore, our genome analysis suggests that *F. straubiae* may, in principle, be capable of running a roTCA cycle (see Figure 2, where citrate cleavage is indicated with question marks). Based on the high OGOR (*korA/B*) transcript abundancy determined in this study under autotrophic conditions, it supports the involvement of the roTCA cycle in CO_2_ fixation.

The findings that CS expression was highest under heterotrophic conditions while *korA/B* were down regulated, suggest forward (oxidative) reaction of the TCA under this condition. As the CS transcript level under autotrophic conditions was not strikingly higher (even lower), the results further imply that the CS concentration might be insufficient to support the reverse flux under autotrophic growth. Taken together, these observations suggest that although *F. straubiae* possesses the genetic potential for roTCA activity, the cycle might not operate in full under the tested autotrophic conditions. Instead, the reversed TCA pathway may function in a linear fashion, working in parallel with and fed by the CBB cycle. Nevertheless, our transcriptomic data are not sufficient to strictly argue whether the reverse direction does operate as a full cycle (as roTCA) or if it stops at citrate (incomplete rTCA). To ultimately track the roTCA further investigations are needed [experimental design and protocols are provided by Steffens *et al*.] (Steffens *et al*. 2022).

#### 6.1.3 Reductive glycine synthase pathway—potential for a novel alternative CO_2_ fixation in F. straubiae?

The reductive glycine pathway is theoretically an autotrophic CO_2_ fixation pathway. However, its role in autotrophic growth is controversial (Bar-Even, Noor, and Milo 2012, Claassens 2021, Mitic *et al*. 2023). The genome of *F. straubiae* does not encode the gene for formate tetrahydrofolate ligase which is the enzyme required for the second step in the reductive glycine synthase pathway: ligation of tetrahydrofolate (THF) and formate under the use of ATP. However, we detected transcripts of all other enzymes involved in the serine aminase variant (Mitic *et al*. 2023) of the reductive glycine pathway under autotrophic conditions which strongly indicates the presence of all the intermediates. This made us wonder is there an unknown (novel) functional orthologue to formate tetrahydrofolate ligase forming 10-formyl-THF? Or an alternative novel pathway could also result in the formation of 10-formyl-THF? Is it possible that *F. straubiae* uses a third, partially unknown CO_2_ fixation pathway? Taken together, these data motivate targeted metabolite and isotope-tracing experiments (e.g., LC-MS quantification of 10-formyl-THF and ^13^CO_2_ flux analysis) to test whether *F. straubiae* forms 10-formyl-THF via an alternative C1-assimilation route.

#### 6.1.4 Regeneration of ATP, NAD(P)H, reduced ferredoxin and supply of inorganic carbon

For a better understanding of the overall metabolism of *F. straubiae*, we also focused on the intermediates of the autotrophic CO_2_ fixating metabolic pathways, which are ATP, NADH, NADPH, Ferredoxin, CO_2_ and bicarbonate (HCO_3_^-^). The following sub-section focusses on carbonic anhydrases catalyzing the reversible hydration of CO_2_ to carbonic acid (HCO_3_^-^) while the regeneration of ATP and the reducing agents will be discussed in sub-section ’ATP and reducing equivalent regeneration by the use of the proton motive force’.

##### Carbonic anhydrases the enzyme involved in CO_2_ supply

The spontaneous diffusion of CO_2_ through the outer membrane and its conversion to HCO_3_^-^inside the cell are insufficient to meet the microorganism’s metabolic needs. This issue is typically addressed by carbonic anhydrases, which catalyze the reversible hydration of CO_2_ to HCO₃⁻. This can occur either externally, to improve CO_2_ uptake, or intracellularly, to either trap CO_2_ as HCO_3_^-^ inside the cell and to increase local CO_2_ concentrations [functioning in reverse direction](Smith and Ferry 2000). Bacterial extracellular carbonic anhydrases, are typically of the class α-carbonic anhydrases (α-CA), and can be identified by their peptide signal for transport through the membrane where they fulfil their physiological function (Supuran and Capasso 2017). β- and γ-carbonic anhydrases (β- and γ-CA) are generally located in the cytoplasm, where they supply CO_2_ to enzymes such as RubisCO, and are responsible for maintaining pH homeostasis and performing other intracellular functions (Capasso and Supuran and 2014, Supuran and Capasso and 2016). RubisCO exhibits relatively slow catalytic kinetics and low affinity for CO_2_, necessitating high local concentrations of CO_2_ for efficient function. Consequently, carbonic anhydrase plays a pivotal role in the CBB cycle by accelerating the conversion of bicarbonate to CO_2_, thereby increasing the supply of CO_2_ to RubisCO (Flamholz *et al*. 2019).

We found three β-CAs in the genome of *F. straubiae*, all of which are located in the cytoplasm according to SignalP predictions, but we could not identify any α-CAs (see Figure 2: ’β-carbonic anhydrases’). The absence of α-CAs and the lack of β-CAs in the periplasm are common observations for bacteria (Supuran and Capasso and 2016). Of the three β-CAs identified, two share 47% similarity and only 23–21% similarity with the third β-CA (IMG ID: 9699, encoded by cynT 3). The two similar β-CAs (IMG IDs 8547 and 8211, encoded by *cynT 1* and *cynT 2*, respectively) are identified as clade B of the β-CA class by InterProScan. *cynT 1* is adjacent to nitrate reductase-encoding genes, while *cynT 2* is adjacent to genes of the CBB cycle. The function of the second β-CA (*cynT 2*) is clearly in supplying CO_2_ to RubisCO; however, the physiological function of the first β-CA (*cynT 1*) is unknown. Notably, *cynT1* is located next to denitrification genes and was most upregulated in autotrophic denitrifying conditions among the three β-CAs (Figure 2: ’β-carbonic anhydrases’). This suggests a role in nitrate-reducing Fe(II) oxidation. We discuss this possibility further in the section *’Could one of the carbonic anhydrases be an isoenzyme functioning as nitrous anhydrase?’*.

Additionally, one γ-carbonic anhydrase-like protein (InterPro family IPR047324) is encoded directly next to the 2-oxoglutarate ferredoxin oxidoreductase (IMG ID: 8187). This protein is significantly and comparably strongly up-regulated under autotrophic conditions compared to heterotrophic conditions (Figure 2: ’γ-carbonic anhydrase-like’). This indicates its role in the reversed TCA cycle, whereby it supplies the 2-oxoglutarate ferredoxin oxidoreductase with CO_2_.

##### 4.1.5.2 ATP and reducing equivalent regeneration for CO_2_ fixation

To power CO_2_ fixation, the CCB and the reverse TCA cycle require a supply of ATP, NADH, NADPH and reduced ferredoxin (Fd). The regeneration of these is, as far as we know, coupled to the use of charge due to the separation of protons (proton motive force, pmf) and sodium ions across the inner membrane (Figure 4) (Bar-Even *et al*. 2012, Bar-Even, Noor, and Milo 2012, Martin and Thauer 2017).

**Figure 4:**
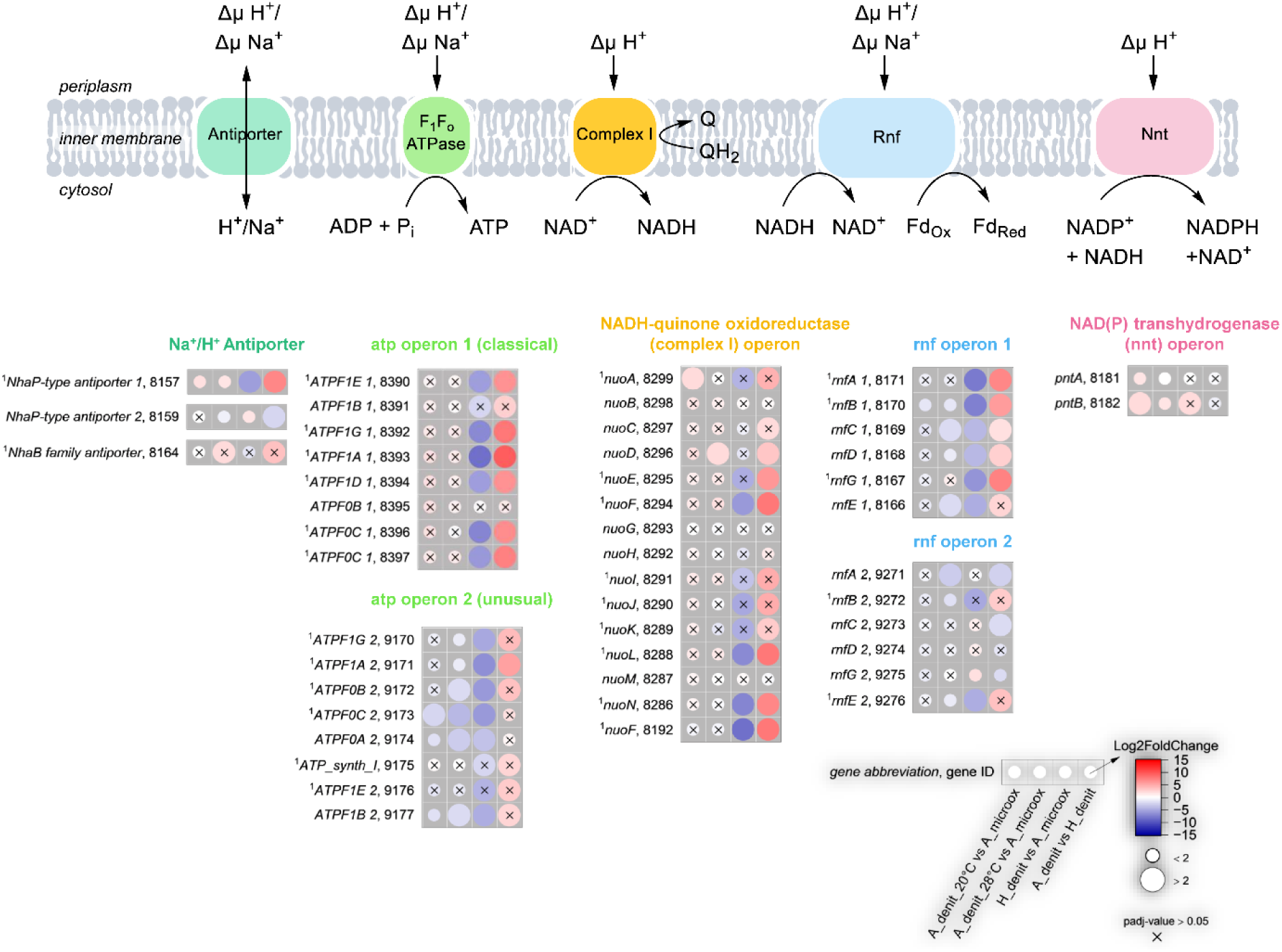
Regeneration of ATP, NADH, NADPH and reduced Ferredoxin. Only the reactions in the direction of the regeneration of the substrates of the CO_2_ fixation metabolic pathways are shown. The log_2_ fold changes of normalized transcript count (Log2FoldChange) are shown as colored dots. The labels follow the structure: gene name, last four numbers of IMG Gene ID. All IMG Gene IDs share the prefix 287840XXXX. Connected rows indicate several subunits of an enzyme while not connected bubble-plots correspond to different enzymes which, in some cases, catalyze the same reaction. The four conditions were: autotrophic denitrifying Fe(II) oxidation at 28°C (A_denit_28°C, n=3), heterotrophic denitrification at 28°C (H_denit, n=1), autotrophic denitrifying Fe(II) oxidation at 20°C (A_denit_20°C, n=2) and autotrophic microaerophilic Fe(II) oxidation at 20°C (A_microox, n=2). *F. straubiae* was grown as community member of culture KS in all nitrate containing conditions. ^1^no transcript detected in H_denit. Gene abbreviation in table S7.

We identified all the proteins commonly suggested to meet the ATP and reducing equivalent demands of cellular metabolism in the genome of *F. straubiae*. These are the Na^+^/H^+^ antiporter, the F1Fo ATPase, the NADH-quinone oxidoreductase (complex I), the Fd^2−:^NAD^+^-oxidoreductase (Rnf complex), and the NAD(P) transhydrogenase (Nnt) (Weber and Senior 2003, Martin and Thauer 2017, Kuhns *et al*. 2020). These proteins catalyze reversible reactions; however, Figure 4 only shows the electron flux towards regenerating substrates of the CO_2_ fixation pathways. We propose that under autotrophic conditions, the whole cell’s NADH demand is met by reverse electron transport via complex I, as the reducing potential of Fe(II) is insufficient to reduce NADH directly. To make the reduction of NADH thermodynamically feasible it requires energy steaming from the pmf. NADH can then be used to reduce ferredoxin (via Rnf) and NADPH (via Nnt), again consuming the pmf. *F. straubiae* possesses two *atp* operons that encode two F_1_F_o_-type ATP synthases. The first operon is classical and encodes the first eight ATP synthase subunits: a, b, c, alpha, beta, gamma, delta, epsilon. The second operon lacks the gene that encodes the delta subunit, which is substituted by a gene that encodes an ATP synthase protein (ATP_synth_I, InterPro family IPR005598). We hypothesize that the second operon might encode Na⁺-translocating ATPases/ATP synthases (N-ATPases), as these operons are always present alongside operons that encode typical H⁺-transporting F-type ATPases (Dibrova, Galperin, and Mulkidjanian 2010). One characteristic of N-ATPases is the absence of the delta subunit. A third indicator of N-ATPases is the presence of a Glu residue in the middle of both transmembrane helices of the c-subunit. However, this does not hold true for the c-subunit encoded by the second ATP operon of *F. straubiae*. This observation challenges our hypothesis and raises questions about whether it is H⁺- or Na⁺-specific.

*F. straubiae* has two operons that encode the six-subunit Rnf complex. Notably, the B subunit of the Rnf complex varies in length among different bacteria due to the presence of different numbers of 4Fe-4S clusters in the C-terminus of the protein (Kuhns *et al*. 2020). Both RnfB subunits in *F. straubiae* have two 4Fe-4S cluster domains (identified by InterProScan), making them rather small. For comparison, *Acetobacterium woodii* has six predicted 4Fe-4S clusters. The implications of these variants are not yet understood.

Comparison of the normalized transcriptional levels of the two *rnf* operons (Figure 4) suggests that *rnf* operon 1 is more strongly expressed under autotrophic conditions, while the expression pattern of *rnf* operon 2 is less distinct and may reflect a function relevant across all tested conditions.

Notable differences in the expression of the genes encoding Na^+^/H^+^ antiporter, F_1_F_o_ ATPase, complex I, Rnf complex and Nnt for denitrifying-autotrophic and microoxic-autotrophic conditions where not observed, indicating that both metabolisms are similar for the regeneration of ATP and reducing equivalents.

### 6.2 First electron acceptor in the electron transfer chain — the Fe(II) oxidases

Fe(II) oxidation by *F. straubiae* has been proposed to involve one or more of the following putative Fe(II) oxidases: two *c*-type cytochromes, MtoAB_KS_ and Cyc2_KS_, and the multi copper oxidase MofAC_KS_ (He *et al*. 2016b) (Figure 5A, main characteristics of Fe(II) oxidases summarized in Table 1).

**Figure 5:**
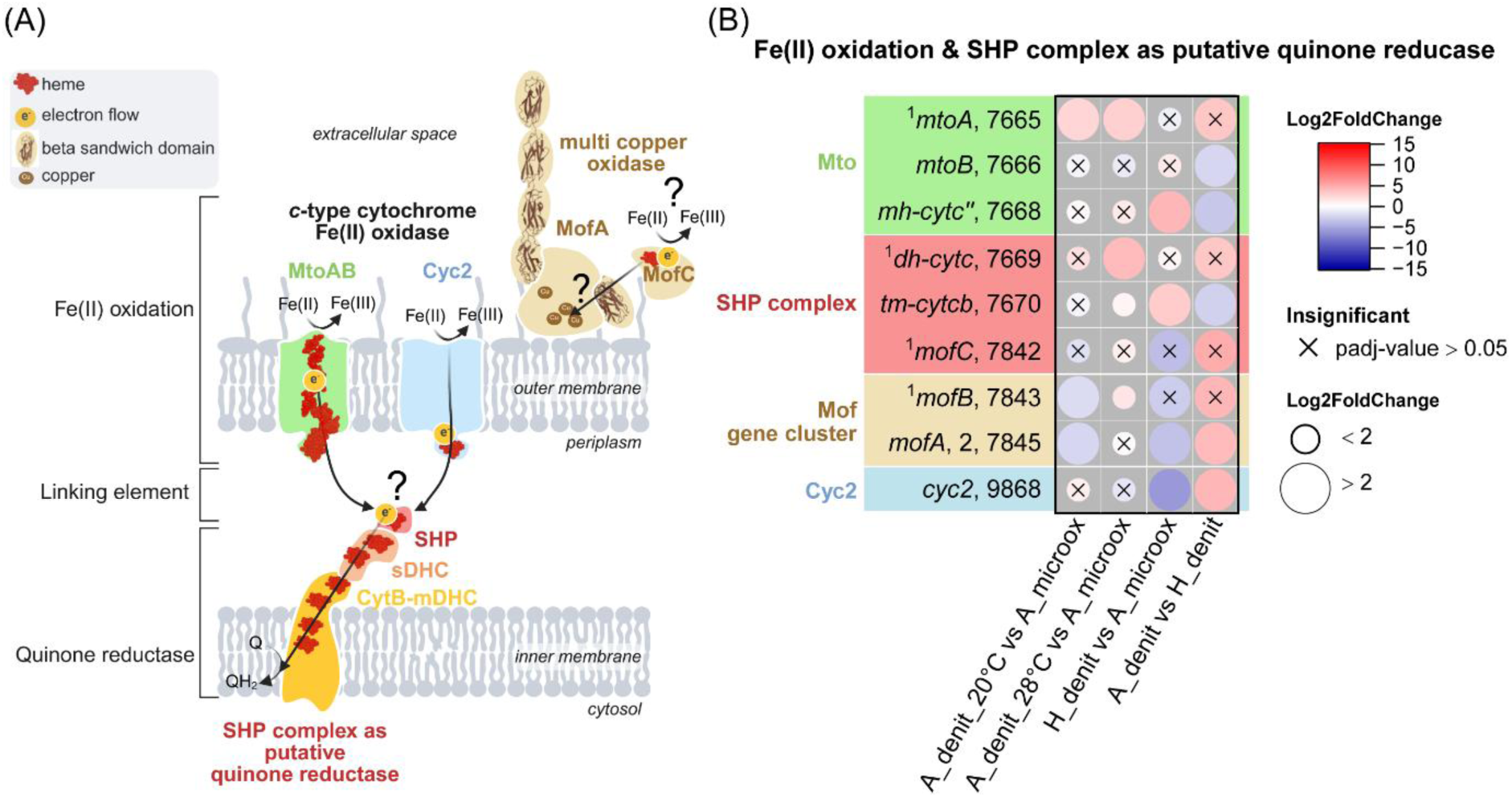
Fe(II) oxidation-related gene expression in *F. straubiae* under denitrifying and microoxic conditions. (A) Putative model of Fe(II) oxidation and downstream flow of the electron in *F. straubiae*. The role of MofAC is hypothetical (He *et al*. 2017) and could also function as Mn-oxidase. (B) The log_2_ fold changes of normalized transcript count (Log2FoldChange) are shown as colored dots. The labels follow the structure: gene name, last four numbers of IMG Gene ID. All IMG Gene IDs share the prefix 287840XXXX. The four conditions were: autotrophic denitrifying Fe(II) oxidation at 28°C (A_denit_28°C, n=3), heterotrophic denitrification at 28°C (H_denit, n=1), autotrophic denitrifying Fe(II) oxidation at 20°C (A_denit_20°C, n=2) and autotrophic microaerophilic Fe(II) oxidation at 20°C (A_microox, n=2). *F. straubiae* was grown as community member of culture KS in all nitrate-containing conditions. ^1^no transcript detected in H_denit. Gene abbreviation in table S7.

**Table 1:** Putative Fe(II) oxidases (MtoA_KS_ and Cyc2_KS_) and Mn-oxidase homologue MofA_KS_ which was hypothesizes to potentially oxidize Fe(II) as well (He *et al*. 2017).

| Characteristics | MtoA<sub>KS</sub> | Cyc2<sub>KS</sub> | MofA<sub>KS</sub> |
| --- | --- | --- | --- |
| IMG ID | 2878407665 | 2878409868 | 2878407845 |
| Homologs | MtrA<sub>S</sub>, *oneidensis*<br>PioA<sub>R</sub>, *ferrooxidans* | Cyc2<sub>At</sub>, *ferrooxidans* | OmpB<sub>G</sub>, *sulfurreducens* MofA<sub>L</sub>, *discophora* |
| Cellular location | Outer membrane | Outer membrane | Extracellular space |
| Electron transfer into the cell | yes | yes | might reduce oxygen outside |
| Metalloprotein | c-type cytochrome<br>multiheme cytochrome<br>→10 hemes | c-type cytochrome<br>→1 heme | multicopper oxidase<br>→4-5 Cu |
| Gene neighborhood | MtoB: 2878407666 (outer membrane protein, β-barrel)<br>Chaperone: 2878407667<br>Putative 'quinone reductase complex':<br>2878407668-2878407670<br>sphaeroides heme protein: 2878407668 | Fe-S-cluster containing protein:<br>2878409869<br>adenylylsulfate kinase:<br>2878409870 | MofB: 2878407843 (chaperon)<br>MofC: 2878407842 (cytochrome c) |
| Structural features | MtoA and B form a complex, multiple hemes spanning a barrel across the outer membrane | B-barrel, one heme domain | 8 beta sandwich motifs potentially involved in mineral attachment |

First, the results of our transcriptome study focusing on these three proteins will be discussed. Then we show the results of in-situ analysis of Cyc2KS and MtoAKS protein sequences and 3D structure predictions to gain functional insights.

Although MtoA is discussed here as a potential Fe(II) oxidase, as proposed by He *et al*., its involvement in Fe(II) oxidation remains less supported than that of Cyc2 and the MtoA/B system, for which homologs with experimental evidence for Fe(II) oxidation have been described. Analysis on MofA_KS_ protein sequence and 3D structure prediction are in the supplemental information (SI-Text 2). Further we show the negative results of searching the genome for an alternative multi-copper Fe(II) oxidase listing all multi-copper oxidases in the supplemental information (SI-Text 3).

#### 6.2.1 Cyc2 is the most dominant Fe(II) oxidase in F. straubiae

MtoA_KS_ was present solely in the Fe(II)-oxidizing conditions and was up-regulated 5.66 times (*P* value 0.001) in the presence of Fe(II)Cl_2_ (denitrifying condition) compared to the presence of FeS (microoxic condition) (SI-Table 1 and 2: *’Normalized transcripts of genes encoding putative Fe(II) oxidases and Log2FoldChange of Fe(II) oxidases within one condition compared to each other’*). This finding supports the complex role of MtoAB in Fe(II) oxidation. Unlike MtoA, Cyc2 transcripts were present in all four conditions being the highest expressed putative Fe(II) oxidase and specifically up-regulated for Fe(II)-containing conditions while not different for conditions with either Fe(II)Cl_2_ (denitrifying condition) or FeS (microoxic condition) as Fe^2+^ source. This indicates that *cyc2* is constitutively expressed enhancing *F. straubiae’s* ability for fast adaptation to a Fe(II)-containing environment.

#### 6.2.2 Sequence analysis of MtoA outer membrane cytochrome reveals His-Met heme iron ligation as it is more prevalent in photoferrotophic homologues

Based on the experimentally solved structure of MtrAB (6R2Q) (Edwards *et al*. 2020), a homolog to MtoAB_KS_, the architecture of the protein complex of MtoAB_KS_ is as follows: MtoA_KS_ is located inside of the β-barrel protein MtoB_KS_ (Figure 6A).

**Figure 6:**
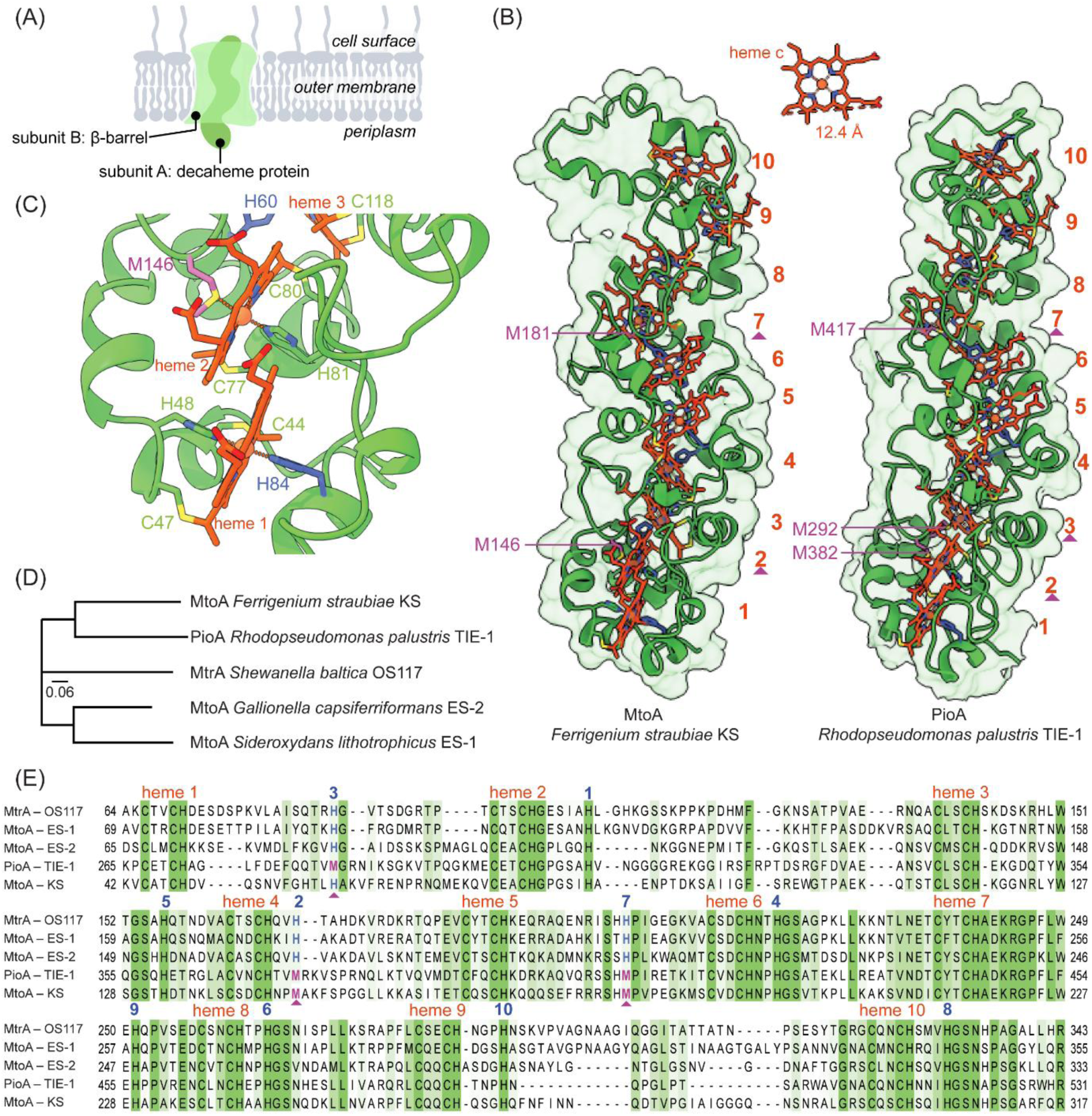
Sequence and in-situ 3D structure analysis of the Fe(II) oxidase MtoA of *F. straubiae*. (A) General architecture of the MtoAB complex based on its structurally resolved homologue MtrAB of *Shewanella baltica* OS117. (B) AlphaFold (2015) models of MtoA of *F. straubiae* and PioA of *Rhodopseudomonas palustris* TIE-1. The hemes coordinated with methionine as the sixth iron ligand are indicated by a purple rectangle triangle beneath the heme number. The hemes are counted from inside (periplasm) to the outside of the bacteria cells. (C) Coordination of iron in the first two hemes (start at N-terminus) of MtoA of *F. straubiae*. (D) Phylogenetic tree of outer membrane cytochrome (subunit A) of photoferrotrophic (PioA) and chemolithotrophic Fe(II)-oxidizers (MtoA of KS, ES-1 and ES-2), and a chemoheterotrophic Fe(III)-reducer (MtrA). (E) Sequence alignment of MtoA, MtrA and PioA, showing the typical heme *c* binding motifs [CXXCH] are highlighted above the alignment by the name and number (red) of the corresponding heme. The counts follow the order as shown in panel B. The histidine (blue) and methionine (purple, highlighted by a rectangle under the alignment) that function as the sixth iron ligand are indicated by the number of the heme to which they bind (blue number, above the alignment).

MtoA_KS_ is a multi heme protein which has 10 heme *c* binding sites, forming a wire spanning the outer membrane throughout MtoB_KS_ (Figure 6B). Interestingly, MtoA_KS_ has more similarity to PioA*_Rhodopseudomonas palustris_* _TIE-1_, which is a Mtr homologue in a photoferrotrophic Fe(II)-oxidizing bacterium, than to MtoA*_Sideroxydans lithotrophicus_* _ES-1_ of a chemolithotrophic Fe(II)-oxidizing bacterium (see phylogenetic tree, Figure 6D). Both were experimentally characterized as Fe(II) oxidases. It is important to note that three of the hemes in PioA*_Rhodopseudomonas palustris_* _TIE-1_ are likely to be His-Met (histidine-methionine) ligated while the hemes in MtoA and MtrA [of the most studied Fe-metabolizing representatives] have solely bis-His (histidine-histidine) ligated hemes (Li DB *et al*. 2020). The ligation with methionine causes higher reduction potentials for the respective hemes compared to histidine ligation. The implementations of this finding are discussed in Lit *et al*. (2020), however, overall inconclusive. His-Met coordination usually increases the Em compared to bis-His (Mao, Hauser, and Gunner 2003, Takayama *et al*. 2008). However, Em is also influenced by the surrounding proteins (H-bonds and solvent exposure), thus a bis-His heme can have a higher Em than a Met-His heme.

Inspired by this work of Lit *et al*. (2020), we performed *in-situ* analysis of the MtoA of *F. straubiae* and found that heme 2 and 7 are likely ligated with methionine (Figure 6B and C). This finding is based on sequence alignment (Figure 6E) and further supported by the AlphaFold model of MtoA_KS_ (Figure 6B).

Lit *et al*. (2020) identified hemes 2, 3 and 7 of PioA*_Rhodopseudomonas palustris_* _TIE-1_ as being most likely His-Met-ligated (Figure 6B and E). Further analysis showed that the methionine ligation of heme 3 is primarily observed in PioA homologues of purple non-sulfur bacteria belonging to the Alphaproteobacteria class, whereas His-Met ligation of heme 2 and 7 [identical to MtoA_KS_] is a conserved feature of PioA homologs from over 30 different bacterial genera. This indicates that this structural feature of MtoA_KS_ is widespread, however, it is an important factor to consider, as it influences the reduction potential of the protein.

#### 6.2.3 Computational modeling of Cyc2 from F. straubiae suggests the formation of a homopolymer

Based on amino acid sequence and de novo modeling of the proteins structure, Cyc2_KS_ has one heme *c* containing domain and a *β*-barrel domain (Figure 7A and B).

**Figure 7:**
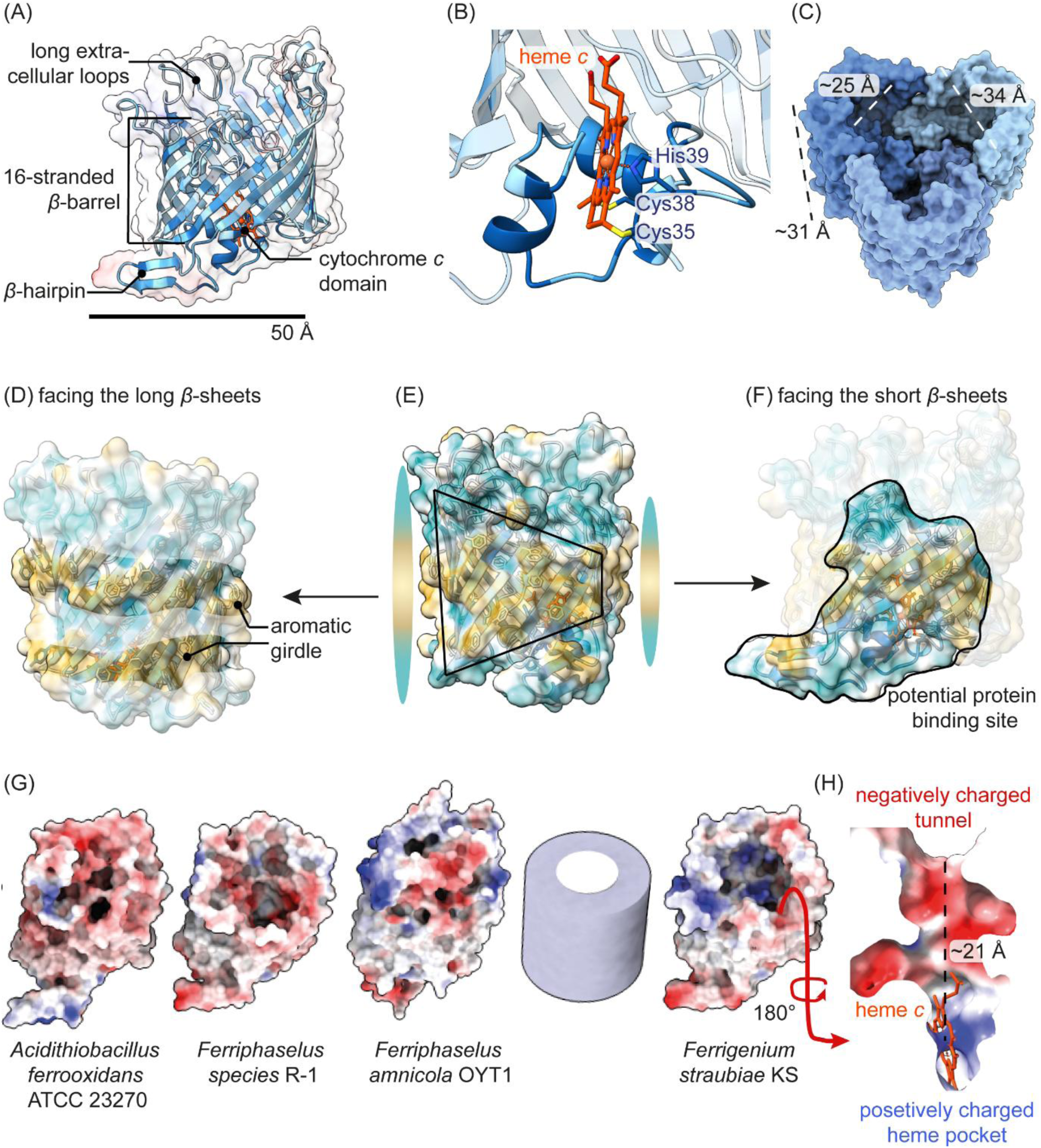
AlphaFold computational protein models of Cyc2 of *F. straubiae* KS (A-F, H) and other Fe(II)-oxidizing bacteria (G). (A) Structural features of Cyc2_KS_. The amino acids of the protein are colored by conservation (high conservation: dark blue; not-conserved: white). Scale bar represents 50 Å in the front layer of the image. (B) Cytochrome *c* domain colored by conservation. The barrel in the background is transparent, thus not colored by conservation. (C) Potential structure of Cyc2 as homotrimer. (D) The aromatic girdle, interacting with the head groups of the lipids of the membrane bilayer. (F) Potential binding surface which has less aromatic amino acids than the barrel surface shown in (D). (E) *β*-sheets differ in length forming a truncated cylinder/barrel. On the right (D) is the surface with long sheets displayed and, on the left, (F) the surface with the short sheets. All aromatic amino acids are shown in the structure (D-F) which is colored by hydrophobicity (hydrophobic surface: brown; hydrophilic surface: blue). (G) AlphaFold models of Cyc2 of different species showing a negatively charged surface at the tunnel entrance towards the heme *c* molecule. (H) negatively charged tunnel towards the heme *c* of Cyc2_KS_. The barrels are similarly tilted as the displayed cylinder, which serves as an orientation reference in the 3D space. (G) and (H) are colored by polarity (positively charged: blue, no charge: white; negatively charged: red).

The cytochrome *c* domain is very conserved while the amino acid sequence forming the *β*-barrel is less conserved (Figure 7A). Experimental work and de novo modeling on Cyc2_PV-1_ was done by Keffer *et al*. (2021). Further Jiang *et al*. (2021 and 2025) identified two iron binding sites likely involved in electron transfer in Cyc2*_Acidithiobacillus ferrooxidans_* (Keffer *et al*. 2021).

Our structure prediction using AlphaFold (Oct. 2025) indicates that Cyc2 has an acidic surface at the inside towards the heme binding pocket (Figure 7G and H). The negatively charged amino acids could either be involved in iron binding presenting an electron transfer mechanism as suggested by Jiang *et al*. (2021) or this negatively charged surface simply binds and complexes Fe(III) and thus prevents Fe(III) from precipitation before it exits the barrel. In addition, we would like to highlight the possibility of a polymeric quaternary structure of the protein as the formation of homopolymers of *β*-barrel complexes has been frequently observed (Hermansen, Linke, and Leo 2022). Interestingly the *β*-sheets on one site are shorter than on the opposing site (Figure 7E). If the β-sheets are shorter than the thickness of the phospholipid bilayer and the protein forms a homopolymer in which these sites connect, the Fe(II) could get closer to the periplasmic space than when the *β*-barrels were equally high at all sites (Figure 7C). The aromatic amino acids located at the outside of the barrel are interacting with the headgroups of the phospholipids, this area is termed the aromatic girdle (Figure 7D)(Hong *et al*. 2007). As the AlphaFold Model of Cyc2_KS_ has less aromatic amino acids at the short *β*-sheets, it might further indicate that it functions as binding site of a polymeric structure (Figure 7F).

### 6.3 Periplasmic electron acceptor after initial Fe(II) oxidation, and quinone reduction

The first electron acceptor that welcomes the electrons from the Fe(II) oxidase at the periplasmic space could be a small cytochrome *c* or copper protein such as rusticyanine (similar to *Thiobacillus ferrooxidans* (Cobley and Haddock 1975) functioning as a link and passing on the electron to a periplasmic or inner membrane redox-active enzyme. The genome of *F. straubiae* has no homologs to rusticyanine. However, it possesses many cytochrome *c* which could potentially fulfil such a function (Figure 8 and Figure 9C).

**Figure 8:**
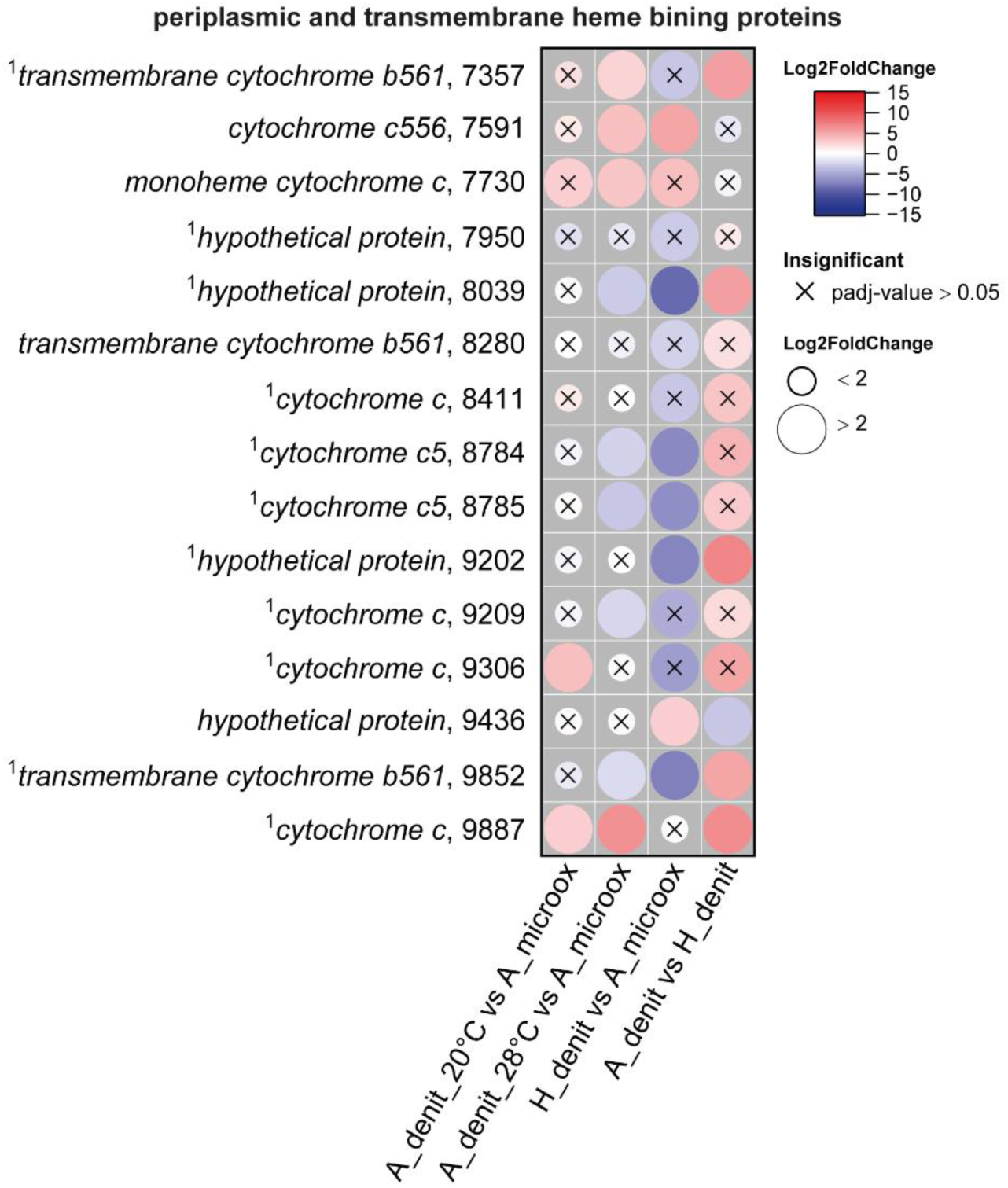
Periplasmic heme binding proteins identified by MHCScan and transmembrane cytochrome *b*561 (trans[inner]membrane) which are not discussed in separate sections. The log_2_ fold changes of normalized transcript count (Log2FoldChange) are shown as colored dots. The labels follow the structure: gene name, last four numbers of IMG Gene ID. All IMG Gene IDs share the prefix 287840XXXX. The four conditions were: autotrophic denitrifying Fe(II) oxidation at 28°C (A_denit_28°C, n=3), heterotrophic denitrification at 28°C (H_denit, n=1), autotrophic denitrifying Fe(II) oxidation at 20°C (A_denit_20°C, n=2) and autotrophic microoxical Fe(II) oxidation at 20°C (A_microox, n=2). *F. straubiae* was grown as community member of culture KS in all nitrate-containing conditions. ^1^no transcript detected in H_denit.

**Figure 9:**
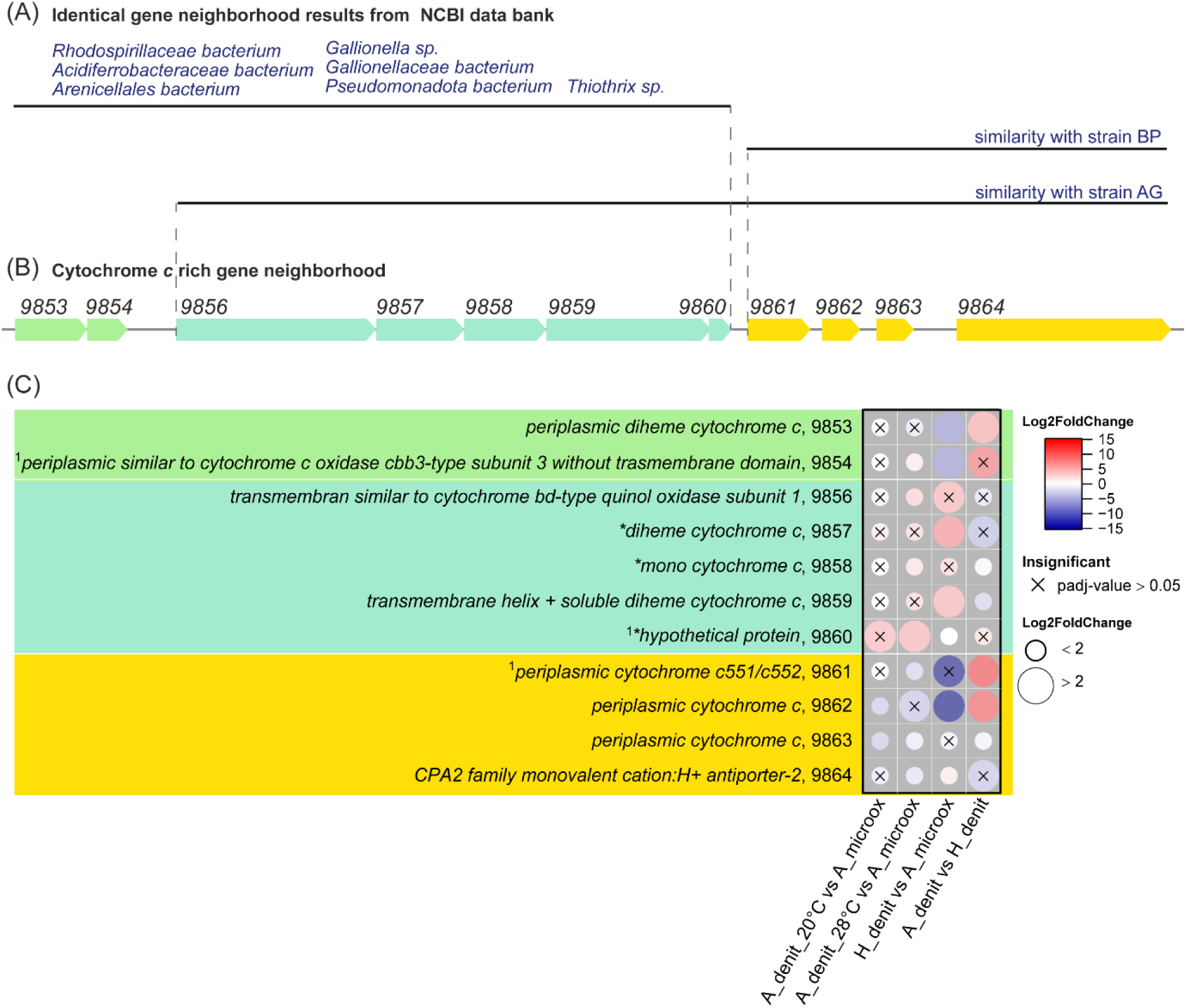
Cytochrome *c* rich gene neighborhood of unknown function. (A) Gene regions identical or similar to those in other bacteria identified by a clinker search using the NCBI database. The black line indicates matching gene patterns in a selection of taxa. The neighborhood was only partially present in the nitrate-reducing Fe(II)-oxidizing bacteria *Ca.* F. altingense strain AG and *Ca.* F. bremense strain BP. (B) Gene neighborhood of scaffold Ga0439409_04, 174669 bp – 185529 bp, of IMG data base. All IMG gene IDs have the format 287840XXXX; the corresponding last four digits are shown above each gene. (C) Gene product descriptions and the corresponding fold changes of normalized counts (Log2FoldChange) of transcripts is shown as colored dots. The labels follow the structure: gene name, last four numbers of IMG Gene ID. The four conditions were: autotrophic denitrifying Fe(II) oxidation at 28°C (A_denit_28°C, n=3), heterotrophic denitrification at 28°C (H_denit, n=1), autotrophic denitrifying Fe(II) oxidation at 20°C (A_denit_20°C, n=2) and autotrophic microaerophilic Fe(II) oxidation at 20°C (A_microox, n=2). *F. straubiae* was grown as community member of culture KS in all nitrate-containing conditions.^1^no transcript detected in H_denit. *no signal peptide was identified by SignalP 6.0, however note that cytochrome *c* biogenesis takes place in the periplasm, thus the cytochromes are expected to be located in the periplasm anyway (Stevens *et al*. 2011).

To identify an electron transport chain that could potentially transfer the electrons from the Fe(II)-oxidase via the periplasmic space to reduce quinone, we screened the *F. straubiae* genome using the software tool MHCScan. This tool searches the genome for interesting gene neighborhoods that could potentially encode multiheme cytochromes spanning the membrane and/or being localized in the periplasm. Among others, this approach identified SHP-like proteins which will be discussed in the section ’*Do SHP-like proteins in F. straubiae form quinone-reducing complexes or function as terminal oxidoreductases?*’. Further, a redox-active complex with similar architecture to the alternative complex III (quinone oxidation and reduction of cytochrome *c*) was identified and it is discussed in the section ’*Novel membrane-bound redox complex upregulated during nitrate-reducing Fe(II) oxidation*’.

The highest number of cytochrome-encoding genes was identified in the gene neighborhood discussed in the section *’Cytochrome c rich gene neighborhood of unknown function’*.

The multiheme cytochromes with the highest number of heme-binding motifs, identified using MHCScan, were MtoA (a decaheme cytochrome c; IMG Gene ID: 2878407665), a novel hexaheme cytochrome *c* (2878408594), and a nonaheme cytochrome *c* (28784098774). The first is one of the putative Fe(II) oxidases, the second appears to be specific to denitrification, and the third may be of particular interest as it could form a conductive wire spanning the periplasm to reduce quinone. This protein is discussed further in section *’A novel nonaheme cytochrome c of unknown function’*.

All additional periplasmic heme binding proteins identified by MHCScan (potential first electron acceptors) and transmembrane cytochrome *b*561 (trans[inner]membrane) of unknown function that are not discussed further are listed in Figure 8. Based on the transcription pattern, only cytochrome *c* with IMG Gene ID 2878409887 is specific for nitrate-reducing Fe(II) oxidation (autotrophic denitrification).

#### 6.3.1 Cytochrome c rich gene neighborhood of unknown function

Using MHCScan, we identified a highly interesting gene neighborhood encoding multiple redox-active proteins (Figure 9). Based on gene localization, expression patterns (Figure 9C), and co-occurrence in different bacterial genomes (analyzed using clinker), we differentiated three functional units (indicated in green, turquoise, and yellow in Figure 9B). The first functional unit comprises two genes encoding cytochrome *c* proteins. The second functional unit contains a transmembrane-spanning cytochrome (IMG Gene ID 2878409856) as well as two periplasmic cytochromes and one which has a transmembrane-spanning helix (IMG Gene ID: 2878409859). The third functional unit consists of three cytochromes followed by a gene encoding a CPA2-family monovalent cation/H⁺ antiporter.

The complete gene neighborhood encompassing all three functional units was not found in the nitrate-reducing Fe(II)-oxidizing cultures BP and AG; however, partial overlaps were observed with *Ca.* F. altingense strain AG and *Ca.* F. bremense strain BP, as indicated in Figure 9A.

Based on protein sequence computational analysis and the gene conservation among *F. straubiae* and *Ca.* F. altingense, we propose that the genes of the second fuctional unit (2878409856–2878409859) encode a membrane associated octaheme cytochrome heterotetramer (octaheme cytochrome complex). The AlphaFold model of the putative subunit encoded by gene 2878409859 shares structural similarity with the quinol oxidase subunit of cytochrome *bd*-type oxygen reductases (pdb 7NKZ). As it possesses the same conserved histidines coordinated to heme cofactors it likely binds three heme cofactors as well (sequence and structure alignment not shown). Further analysis using COFACTOR (Roy, Yang, and Zhang 2012; Zhang, Freddolino, and Zhang 2017) confirmed the presence of a quinone-binding site revealing it as a quinone-binding/-reducing subunit. Therefore, the octaheme cytochrome complex likely functions in electron transfer and may operate either as quinol oxidases or quinone reductases. Given the similar expression patterns observed under both autotrophic conditions (microaerophilic and denitrifying), this suggests that this octaheme cytochrome complex do not have a specific role in nitrate-reducing Fe(II) oxidation. Nevertheless, it is clearly involved in electron transfer processes and therefore represent interesting targets for further investigation.

In Fe(II) oxidation-related studies of Keffer *et al*. and Zhou *et al*., the gene Slit_1353 (NCBI ID) of *Sideroxydans lithotrophicus* ES-1 encodes a cytochrome and was found to be iron-responsive (Zhou N *et al*. 2022, Keffer *et al*. 2025). The *F. straubiae* cytochromes of the third functional unit (Figure 9B, marked in coral) share 40-55% similarity. The corresponding genes (IMG ID 2878409861-2878409862) were indeed upregulated in the Fe(II)-oxidizing (autotrophic) conditions (Figure 9C). However, especially the transcript of 2878409862 was very dominant in all four conditions compared to the other cytochromes (normalizes transcripts in TPM (Transcripts Per Million) listed in Table S1). This indicates that 2878409862 with 54% similarity to Slit_1353 may also play a very important role in *F. straubiae* in Fe(II) oxidation metabolism.

#### 6.3.2 A novel membrane associated nonaheme cytochrome c of unknown function

We used MHCScan to search the genome for multiheme cytochromes that could potentially span the periplasm and identified a membrane associated nonaheme cytochrome *c* (NHC; Figure 10B A; IMG ID: 28784098774). The corresponding gene is located downstream of the copper-containing nitrite reductase (nirK) gene and upstream of a Sphaeroides heme protein-like protein gene (Figure 10B). The genes in the immediate neighborhood encode a transmembrane protein (28784098775) and a [4Fe–4S] cluster-containing protein (28784098776). Analysis of these proteins predicted one transmembrane helix for the NHC and five for the transmembrane protein. As none of the three proteins contain a signal peptide for transport into the periplasm, the NHC and the transmembrane protein are likely anchored in the inner membrane, with the nonaheme domain of the NHC located in the periplasm (Figure 10A).

**Figure 10:**
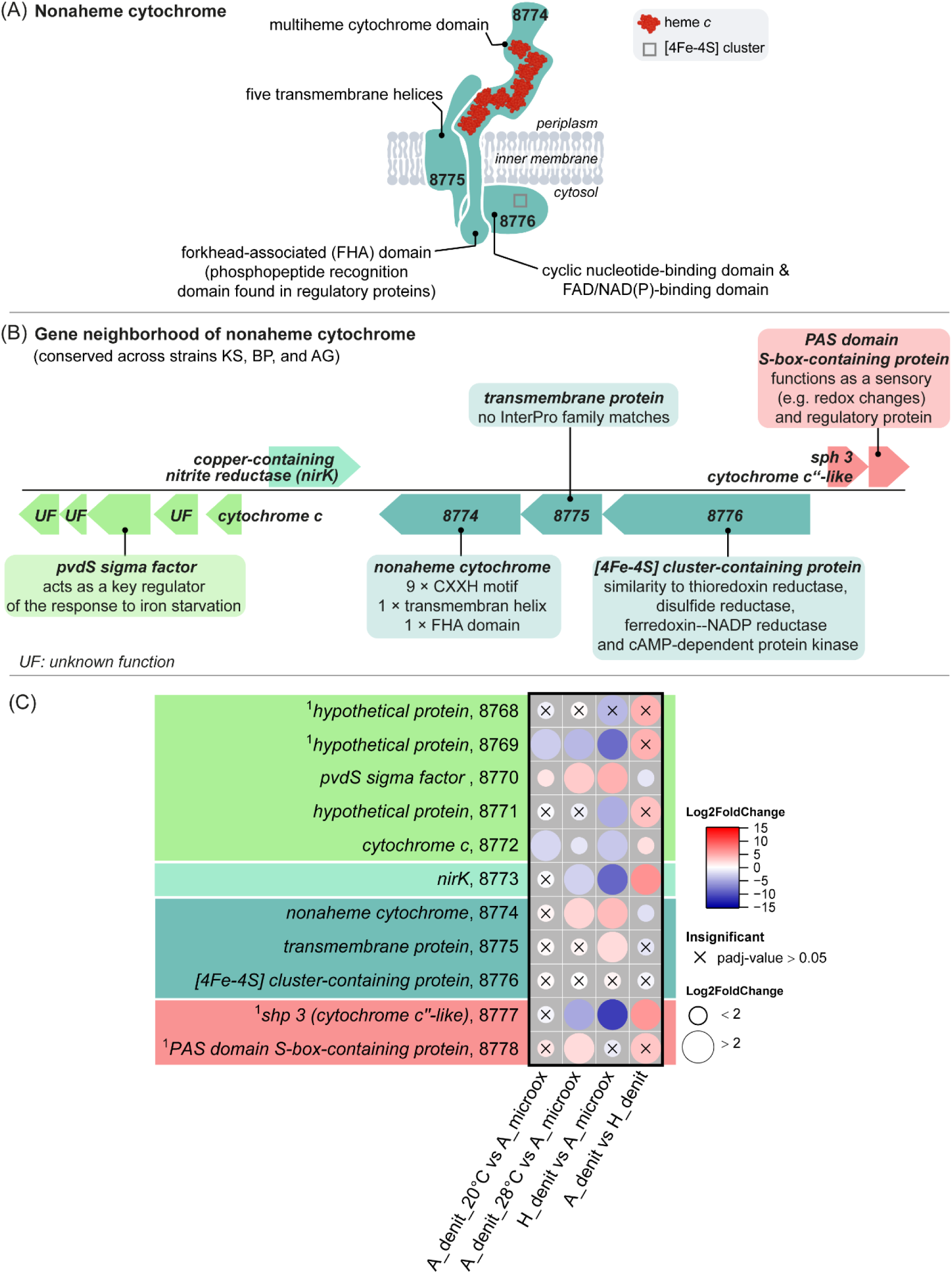
(A) Scheme of the predicted architecture of a novel nonaheme cytochrome *c* shown in putative interaction with proteins encoded in close genomic neighborhood. The overall shape of the nonaheme cytochrome *c* represents its AlphaFold model (not shown). (B) Gene neighborhood. All IMG gene IDs have the format 287840XXXX; the corresponding last four digits are shown above each gene. [Reference of PvdS sigma factor: (Imperi *et al*. 2010)] (C) Gene product descriptions and the corresponding fold changes of normalized counts (Log2FoldChange) of transcripts is shown as colored dots. The labels follow the structure: gene name, last four numbers of IMG Gene ID. The four conditions were: autotrophic denitrifying Fe(II) oxidation at 28°C (A_denit_28°C, n=3), heterotrophic denitrification at 28°C (H_denit, n=1), autotrophic denitrifying Fe(II) oxidation at 20°C (A_denit_20°C, n=2) and autotrophic microaerophilic Fe(II) oxidation at 20°C (A_microox, n=2). *F. straubiae* was grown as community member of culture KS in all nitrate-containing conditions.^1^no transcript detected in H_denit.

Structurally, the NHC resembles a nanowire that could transfer electrons from outermembrane Fe(II)-oxidases, such as MtoA and Cyc2, to the inner membrane. This is supported by the study of Tian *et al*. which demonstrated experimentally, that Fe(II) can reduce quinone under physiological conditions (Tian, Zhang, and Yuan 2024). A connection between MtoA and the NHC would create a nanowire comprising 17 hemes, making it a promising bridge for facilitating electron transfer. When connecting with Cyc2, it would result in a shorter nanowire consisting of 10 heme cofactors. As all heme-binding sites of the NHC are located outside the membrane, it remains unclear whether electrons can approach quinone closely enough to enable its reduction. However, NHC has architectural similarity to NapC/NirT-family quinol dehydrogenases such as CymA (a quinone reductase in the iron reducer Shewanella) (Marritt *et al*. 2012) and cytochrome *c* nitrite reductase subunit H (NrfH), which functions as the mediator between the quinone pool and cytochrome *c* nitrite reductase subunit A (NrfA) in *Wolinella succinogenes* (Simon *et al*. 2000). Both proteins are composed of a periplasmic tetraheme cytochrome domain and only a single transmembrane helix as well. Because of this similarity we hypothesize that he electrons from the NHC can reach the quinone pool in a similar manner as the quinol pool can reduce the tetraheme of CymA and NrfA.

One could also hypothesize that the NHC transfers electrons to the transmembrane protein (28784098775), which then reduces quinone. However, the transmembrane protein showed no matches to redox active or any other protein motifs (families, domains, or profiles) in the InterPro database. As transmembrane helices often act as binding platforms, this protein may instead stabilize partners that interact with the NHC in the periplasm and potentially connect it to the [4Fe–4S] cluster-containing protein in the cytosol. The [4Fe–4S] cluster-containing protein showed similarity to several known structures in the PDB database (Figure 10B). However, the available information is insufficient to infer its biological function.

Notably, the NHC contains a forkhead-associated (FHA) domain on the cytosolic side, which is typically associated with regulatory functions. To our knowledge, FHA domains have not previously been reported in combination with multiheme cytochromes. Sequence searches (HHpred against the PDB database) and structural comparisons (AlphaFold model analyzed using FoldSeek) identified homologues for individual domains, but not for proteins containing both domains.

While NHC is a promising candidate for linking Fe(II) oxidation to quinone reduction, the function and interaction partners of NHC and its closely encoded proteins remain unclear. The gene cluster was expressed under all tested conditions, which suggests its importance for all of the conditions tested (Figure 10C). Further, NHC is conserved across strain KS, BP and AG, highlighting it as a potential marker gene for *Ferrigenium* spp.-dominated autotrophic nitrate-reducing Fe(II)-oxidizing communities.

### 6.4 Do SHP-like proteins in *F. straubiae* form quinone-reducing complexes or function as terminal oxidoreductases?

The sphaeroides heme protein was discovered 1985 in the purple phototrophic bacterium, *Rhodobacter sphaeroides*. Based on its amino acid sequence and the protein crystal structure, it is a member of the cytochrome *c*_4_ superfamily of class I cytochromes. What was initially believed to be an unusual rare heme protein, occurs actually in many proteobacterial species. An excellent summary of the SHP findings is presented in the introduction section of Mayer *et al*. 2010, who studied the role of SHPs in photosynthetic bacteria in the family of *Rhodobacteraceae* (Meyer Terry E, Kyndt, and Cusanovich 2010).

In most species, SHP-like proteins are located in a three-gene operon that also encodes a soluble diheme cytochrome *c* (sDHC) and a membrane-spanning cytochrome *b* (CytB). The latter is sometimes extended by an extra domain which is a chimera of the diheme cytochrome *c* connected to the membrane spanning CytB (CytB-mDHC) (Figure 11). Furthermore, their gene expression seems to be regulated by a two-component system which is encoded in close gene approximate in most cases (Figure S7). Indeed *F. straubiae* has three variations of SHP-like protein gene neighborhoods: SHP-like protein gene associated with sDHC and CytB-mDHC gene (2878407668 (*shp1 1*), 2878407669 [*sDHC 1*], 2878407670 [*cytB-mDHC*]), associated with sDHC and CytB gene (2878408741 [*shp 2*], 2878408742 [*sDHC 2*] and 2878408743 [*cytB*]) and standing alone (2878408777, *shp 3*). The SHP 1 and SHP 2 likely form a complex with the sDHC and the membrane-bound CytB-mDHC and CytB, respectively (De March *et al*. 2015). The SHP1 complex was already spotted by He *et al*. (2016) as it is encoded close to the putative Fe(II)-oxidase genes *mtoAB* (860 bp up-steam of *shp 1*). They hypothesized that the complex might form the linkage of electron transfer from MtoA to quinone (He *et al*. 2016b). The role of the SHP-like protein and its complexes remains unclear, and the idea that these proteins might reduce quinones is highly speculative. However, the hypothesis that the SHP complex links MtoAB to the quinone pool has increasingly appeared in the literature, despite the fact that this connection has not yet been experimentally confirmed.

**Figure 11:**
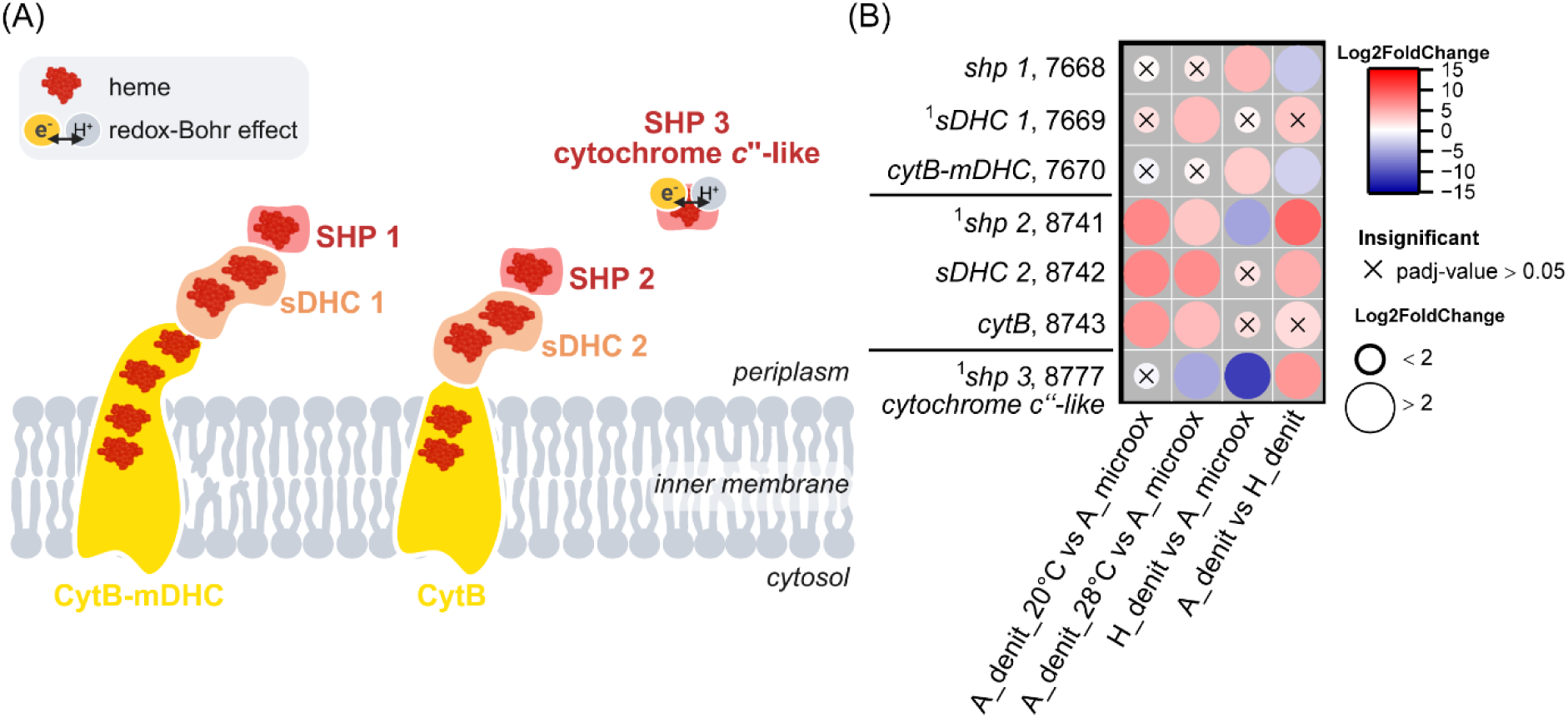
(B) Sphaeroides heme protein-like protein (SHP) hypothetical complex with soluble diheme cytochrome *c* (sDHC) and either cytochrome *b* (CytB) or cytochrome *b* extended by chimera of the diheme cytochrome *c*. Sequence analysis revealed that SHP3 is more similar to cytochrome *c*’’ than to the Sphaeroides heme protein (SHP), which is also a homologue. (B) The log_2_ fold changes of normalized transcript count (Log2FoldChange) are shown as colored dots. The labels follow the structure: gene name, last four numbers of IMG Gene ID. All IMG Gene IDs share the prefix 287840XXXX. The four conditions were: autotrophic denitrifying Fe(II) oxidation at 28°C (A_denit_28°C, n=3), heterotrophic denitrification at 28°C (H_denit, n=1), autotrophic denitrifying Fe(II) oxidation at 20°C (A_denit_20°C, n=2) and autotrophic microaerophilic Fe(II) oxidation at 20°C (A_microox, n=2). *F. straubiae* was grown as community member of culture KS in all nitrate-containing conditions. Gene abbreviation in table S7.

Based on experimental studies on the SHP complex of *Rhodobacter sphaeroides* and *Shewanella baltica*, the membrane-bound CytB reduces quinones, after which the electrons are transferred to the tightly bound sDHC, and subsequently to the SHP-like protein, which also associates tightly with the sDHC (Gibson *et al*. 2006, Di Rocco *et al*. 2011). These findings, together with conclusions from other studies on homologous systems such as cytochrome *c*’’ of *Methylophilus methylotroph*, support the idea that SHP and SHP-like proteins act as the catalytic unit of the complex and are responsible for substrate reduction (Leys *et al*. 2000, Brennan *et al*. 2001, Enguita *et al*. 2006, Li BR *et al*. 2008).

Cytochrome *c*’’ of *Methylophilus methylotroph* is the best studied structural homolog of SHP, however its biological function is thought to be distinct (Enguita *et al*. 2006). One of the α-helices of cytochrome *c*’’ undergoes movement upon reduction, whereas the SHP structure appears more rigid. Furthermore, cytochrome *c*’’ shows redox-Bohr effect behavior suggesting proton translocation as its characteristic structural feature. The redox–Bohr effect is the thermodynamic coupling between reduction of the heme and protonation of nearby ionizable amino acid side chains (e.g., histidine residues). Upon reduction of the heme iron, the axial histidine ligand detaches and becomes protonated. Thus, the redox–Bohr effect describes the coupling of a redox process with changes in the protonation state of the surrounding protein environment. The potential proton transfer pathway was well described by Enguita *et al*. 2006.

Both proteins have in common that the heme axial ligand detaches upon reduction, opening a binding site for small molecules such as O_2_, CO, CN⁻, or NO (usual exogenous heme ligands). This detachment is enabled by an unusual disulfide bridge—uncommon among cytochromes—that makes the protein more rigid and was well described by Brennan *et al*. (Figure 12A) (Brennan *et al*. 2001).

**Figure 12:**
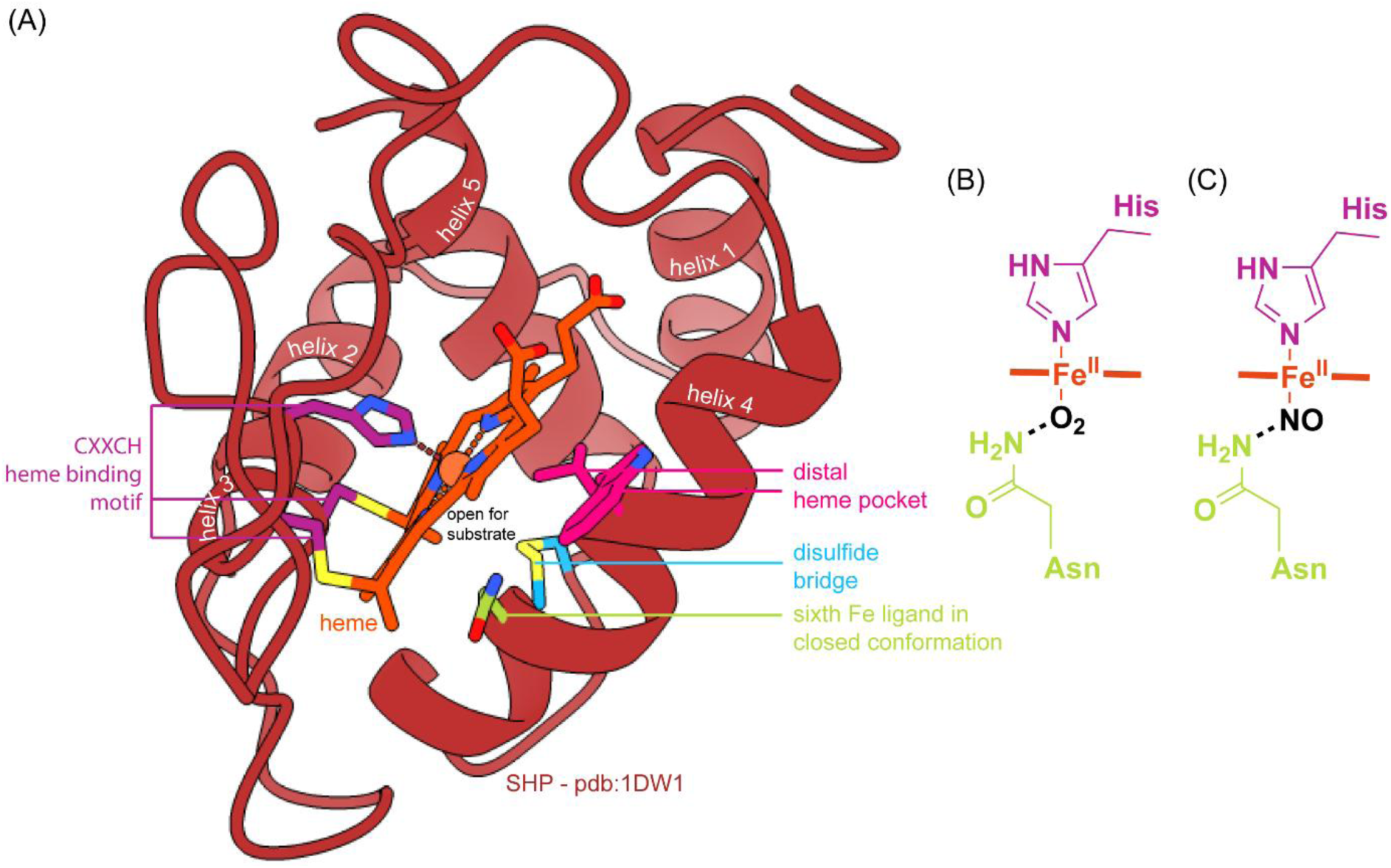
(A) Open substrate binding conformation of the oxidized SHP (*Rhodobacter sphaeroides*) cyanide complex (pdb: 1DW1) (Leys *et al*. 2000). (B) and (C) show the potential binding of a small molecular substrate such as O_2_ or NO to the Fe of heme (Li BR *et al*. 2008).

In fact, SHP has been shown to bind O_2_ (Figure 12B), CO, NO (Figure 12C), CN, azide, nitrogenous bases such as hydroxylamine, bis-tris-propane, HEPES, Tris and taurine (Meyer and Cusanovich 1985; Klarskov *et al*. 1998; Leys *et al*. 2000), and to possess nitric oxide dioxygenase activity (Li *et al*. 2008). Despite these findings, the biological function of cytochrome *c*’’ and SHP remain unknown.

To gain preliminary insight into the *F. straubiae* SHP-like protein and its complexes, we focused on analyzing SHP-like protein sequences (catalytic unit, reducing the terminal substrate) instead of examining sDHC CytB or CytB-mDHC in detail. We therefore compared the SHP-like proteins of *F. straubiae* with the three best-studied SHP/SHP-like representatives: cytochrome *c*’’ of *Methylophilus methylotrophus* ATCC 53528 (Taxon ID: 2518645588), the SHP-like protein of *Shewanella baltica* OS117 (a metal oxide-reducing bacterium; Taxon ID: 651053066), and the original SHP of *Rhodobacter sphaeroides* 2.4.1 (Taxon ID: 8067927082). In addition, we broadened our analysis to include SHP-like proteins from other well-known iron-metabolizing bacteria. These include the nitrate-reducing Fe(II)-oxidizers *Candidatus* Ferrigenium altingense AG (Taxon ID: 2860363623), *Candidatus* Ferrigenium bremense BP (Taxon ID: 2831290873), and *Thiobacillus denitrificans* ATCC 25259; the microaerophilic Fe(II)-oxidizers *Ferrigenium kumadai* An22 (Taxon ID: 2927562029), *Mariprofundus ferrooxydans* PV-1 (Taxon ID: 639879965), *Gallionella capsiferriformans* ES-2 (Taxon ID: 648028028), and *Sideroxydans lithotrophicus* ES-1 (gene Slit_1323 within the iron-responsive cluster (Zhou N *et al*. 2022, Keffer *et al*. 2025); Taxon ID: 646688985); as well as the magnetite-forming magnetotactic bacteria *Magnetospirillum magnetotacticum* and *Magnetococcus* sp. MC-1. [The genes were identified by BLAST searches of IMG genomes using 2878407668 as the seed. Because the surrounding gene neighborhoods encode CytB and sDHC, we believe that the cytochromes, although small in size, are not random hits.]

Phylogenetic analysis and gene neighborhood comparison (Figure S7, +/- 10,000 bp) of the selected SHP-like protein homologs show that two of the *F. straubiae* SHP-like proteins (**SHP 2** and **SHP 3**) cluster with homologues of the two other main Fe(II)-oxidizers (*Candidatus* Ferrigenium altingense AG and *Candidatus* Ferrigenium bremense BP) of nitrate-reducinging Fe(II)-oxidizing cultures (In Figure 13 and Figure 15, indicated by yellow box. Gene neighborhood comparison: FigureS7).

**Figure 13:**
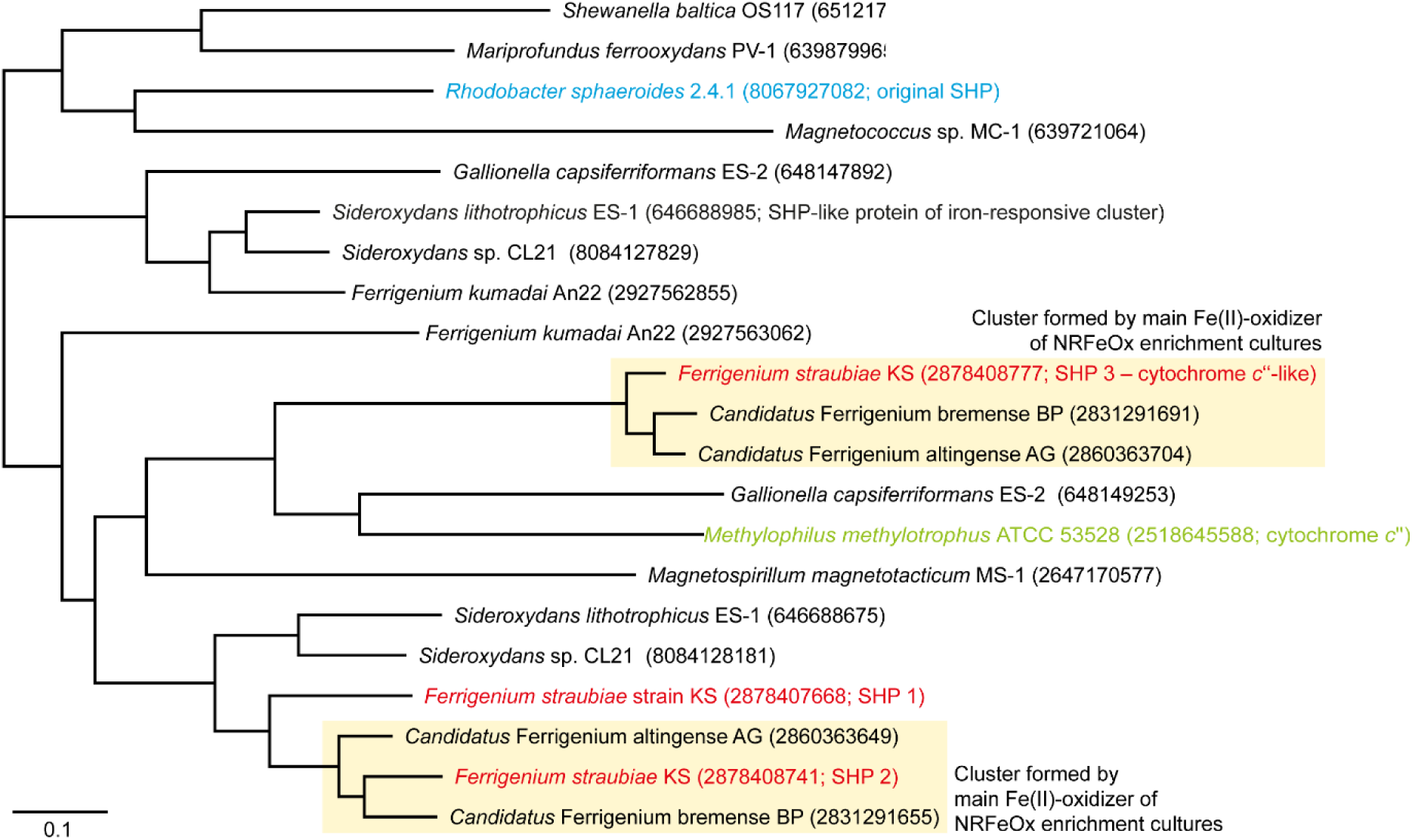
Phylogenetic tree of SHP (blue) (Leys *et al*. 2000), cytochrome *c*″ (green) (Enguita *et al*. 2006), and SHP-like proteins identified in iron-metabolizing bacteria. The three SHP paralogs of *F. straubiae* (SHP1–3) are highlighted in red. SHP-like protein from *Sideroxydans lithotrophicus* ES-1 has been found in an iron-responsive gene cluster (Zhou N *et al*. 2022).

The gene neighborhood of **SHP 2** includes components of the Tol/Pal transport system (according to the annotated by the IMG pipeline). However, the TolQ-TolR-TolA inner membrane complex shares structural similarities with the TonB-ExbB-ExbD siderophore (iron-chelator) uptake system. Therefore, it is conceivable that these proteins contribute to iron transport, and that the nearby **SHP 2** may be specifically related to Fe(II) oxidation metabolism.

The up-regulation of **SHP 2** under nitrate-reducing Fe(II)-oxidizing conditions compared to the microoxic condition was one of the most outstanding findings of the transcriptomic data (Figure 11B, Figure S4), making it a highly interesting candidate for further investigation of NRFeOx metabolism. It is remarkable how little is known about the physiological function of SHP proteins, despite the observation that **SHP 2** appears to be among the most relevant proteins in NRFeOx metabolism (Meyer Terry E, Kyndt, and Cusanovich 2010). This clearly indicates that research on SHP proteins should be resumed.

The **SHP 3** gene neighborhood contains a copper nitrite reductase (NirK), which is typically associated with nitrite reduction under microoxic conditions (Figure 10B). Therefore, although the nitrate-reducing, Fe(II)-oxidizing strains in cultures KS, BP and AG all encode this SHP-like protein, it is unlikely to play a direct role in the nitrate-reducing, Fe(II)-oxidizing pathway. Given that its expression pattern is similar to that of NirK (Figure 10C), its substrate could potentially be nitric oxide (NO), the product of nitrite reduction by NirK. It is noteworthy that **SHP 3** is not encoded next to the membrane-spanning potential quinone reductase CytB/CytB-mDHC, raising the question of what its reducing partner might be and whether it forms an SHP complex with CytB/CytB-mDHC, which is encoded at a different location in the *F. straubiae* genome and by this replacing SHP 1 or 2. On the other hand, cytochrome *c*’’ is also a ’*stand-alone’* SHP, and given its structural differences from the Sphaeroides heme protein, it may fulfill a different function that does simply not require the membrane-bound component.

**SHP 1**, which was hypothesized to act as the linking element between the Fe(II) oxidase (genetically located next to mtoAB) and the quinone pool, was not detected in *Candidatus* Ferrigenium altingense AG and *Candidatus* Ferrigenium bremense BP (both of which also lack *mtoAB*). Moreover, the association of SHP 1 with *mtoAB* appears to be unique to *F. straubiae*, as a clinker search across the entire NCBI genome database did not reveal a comparable gene cluster in other organisms. This raises the question of whether the proximity of the SHP-encoding genes to *mtoAB* in *F. straubiae* is merely coincidental, implying that SHP may not play a role in electron uptake from MtoAB and transfer to the quinone pool. Instead, this interpretation is consistent with observations suggesting that these SHP-like complexes are more likely involved in quinone oxidation rather than quinone reduction. Interestingly, the transcript of the *shp 1* and *cytB-mDHC* complex peaked out for the heterotrophic conditions while the transcript of *sDHC* was not present. Although the genes lay next to each other, it might be possible that *sDHC* is not co-expressed with *shp 1* and *cytB-mDHC* and further that the chimera mDHC is taking the function. At this stage, based on the limitations of the heterotrophic dataset, this remains highly speculative. However, to understand the metabolism of *F. straubiae* under heterotrophic conditions, this complex is certainly in the spotlight. A better understanding should be achieved through further transcriptomics and, above all, native mass spectrometry.

The SHP- like protein (IMG ID: 646688985) of strain ES-1 which was found to be located in an iron-responsive cluster (Zhou N *et al*. 2022, Keffer *et al*. 2025) did not form a cluster with any of the NRFeOx bacteria (*F. straubiae*, *Candidatus* Ferrigenium altingense AG and *Candidatus* Ferrigenium bremense BP) but was found in one cluster with homologs of strain An22 and strain ES-2. This indicates that these iron-related genes do not play a role for nitrate-reducing Fe(II) oxidation.

The sequence alignment revealed that **SHP 1** and **2** are more similar to SHP, sharing the same sixth Fe ligand (asparagine) and the same amino acids in the distal heme pocket (W and F) (Figure 14). In contrast, **SHP 3** has histidine as sixth Fe ligand as it is the case for cytochrome *c*’’. And further they have the size of the first helix in common (based of SHP 3 Alphafold model and crystal structure of cytochrome *c*’’ [pdb – 1OAE]) which is bigger compared to SHP.

**Figure 14:**
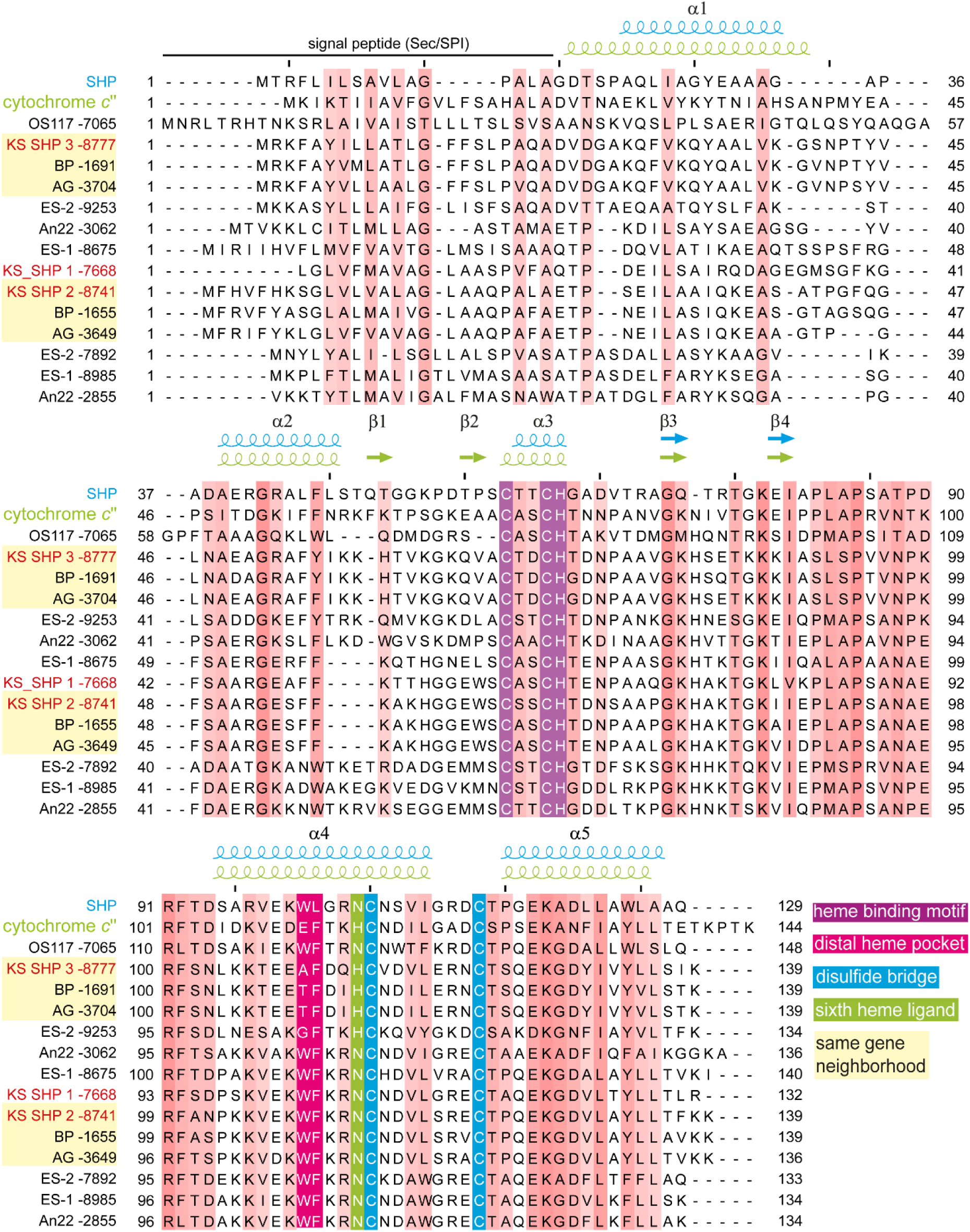
Sequence alignment of SHP-like proteins, with secondary structure motifs indicated based on the protein crystal structures of SHP (pdb – 1DW1) and cytochrome *c*’’(pdb – 1OAE). The yellow box indicates two distinct clusters where the SHP-like protein of strain KS, BP and AG share the same genomic neighborhood.

This finding suggests that **SHP 1** and **2** share the same structural mechanism with SHP protein which likely reduces its yet unknown substrate and might function as oxygenase. In contrast, **SHP 3** likely shares functional features such as the redox-Bohr effect with its homologue cytochrome *c*’’. This proposes, that its catalyzed reaction necessitates proton translocation.

To better understand which amino acid substitutions are likely involved in differences in substrate specificity and catalytic properties, we analyzed the sequence of **SHP 1** searching for evolutionary coupled amino acid pairs. We found that several residues participating in both short- and long-range evolutionary couplings within the potential substrate binding side. This suggests that substitutions within these coupled positions may drive functional divergence among SHP-like proteins which might be very diverse in function (Figure S8).

In summary, SHP and SHP-like proteins likely function as terminal oxidoreductases, but appear to reduce different small molecular substrates and thus participate in distinct metabolic pathways. Nevertheless, the presence of this protein family in diverse organisms suggests a shared physiological role. One possible common feature is that these bacteria are adapted to environments containing a variety of electron donors with relatively high reducing potentials. If so, SHP-containing microorganisms may represent particularly interesting targets for future research.

### 6.5 Classical nitrogen species reduction could be complimented by previously unrecognized carbon anhydrase and membrane bound redox complexes reducing quinone

#### 6.5.1 Classical denitrification via Nar and Nir

Under denitrifying conditions, the nitrate reductase (NarGHI) and nitrate/nitrite transporter (NarK) genes were up-regulated (Figure 15). *F. straubiae* encodes two types of nitrite reductases, two paralogs each: a cytochrome *cd*_1_ nitrite reductase (NirS) and a copper nitrite reductase (CuNir/NirK). As observed in other studies as well, NirK tends to be more dominant under microoxic conditions, whereas NirS prevails during anaerobic denitrification. Looking at the nitrate reductase (Nar) and the nitrite reductases (Nir), Nar appears to be the only enzyme in the denitrification chain directly contributes to energy conservation in the pmf. The nitrite reductase consumes electrons from cytochrome *c*, thereby positively affects the downstream electron flow that may be indirectly involved in energy conservation.

**Figure 15:**
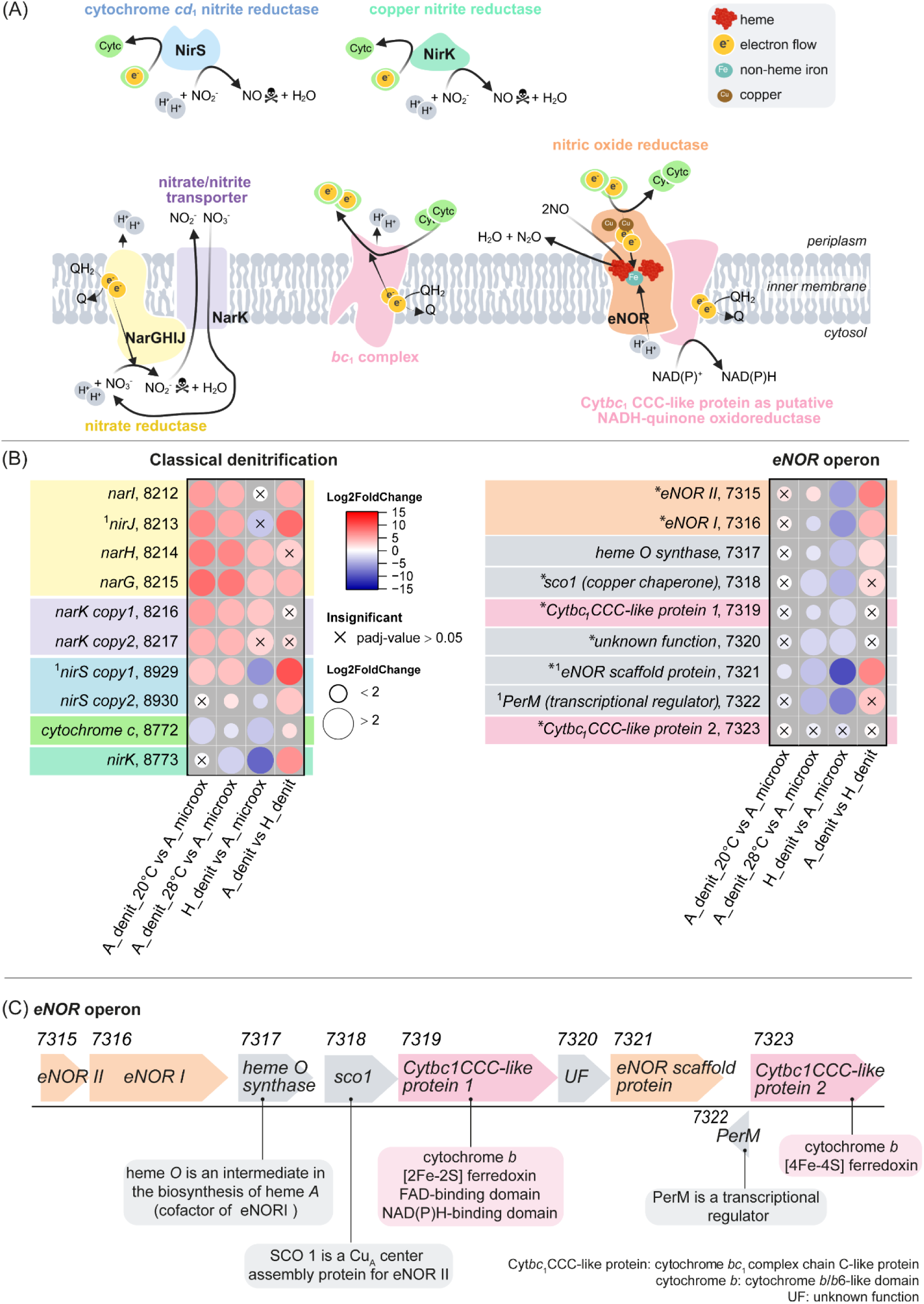
(A) Overview of known (classical) denitrification pathways in *F. straubiae* and the newly classified nitric oxide reductase (eNOR-family). Skull symbol: product is a cell toxin in too high concentrations. (B) The log_2_ fold changes of normalized transcript count (Log2FoldChange) are shown as colored dots. The labels follow the structure: gene name, last four numbers of IMG Gene ID. All IMG Gene IDs share the prefix 287840XXXX. The four conditions were: autotrophic denitrifying Fe(II) oxidation at 28°C (A_denit_28°C, n=3), heterotrophic denitrification at 28°C (H_denit, n=1), autotrophic denitrifying Fe(II) oxidation at 20°C (A_denit_20°C, n=2) and autotrophic microaerophilic Fe(II) oxidation at 20°C (A_microox, n=2). *F. straubiae* was grown as community member of culture KS in all nitrate containing conditions. We found paralogues for *nirK* and *nirS* and named them *copy1* and *copy2*. *present in the proteome of A_denit_28°C [data of Huang’s study (Huang, Straub, Blackwell *et al*. 2021)]. This analysis was conducted exclusively on the eNOR operon. ^1^no transcript detected in H_denit. (C) *eNOR* operon. Cofactors are not indicated for all proteins. Gene abbreviation in table S7.

However, the specific pathway by which electrons reach the redox-active partner of nitrite reductase under autotrophic conditions remains unclear, and therefore it is also uncertain whether nitrite reduction in this mode can support energy conservation. In the following we explain how nitrite reduction indirectly support energy conservation under heterotrophic conditions and then we discuss how nitrite reduction could still positively effects energy conservation under autotrophic conditions.

In the heterotrophic pathway, organic substrates are oxidized via the TCA cycle, generating reducing equivalents such as NADH. These electrons are transferred to NADH-quinone oxidoreductase thereby conserving energy in pmf and reducing the quinone pool (opposite direction as shown in Figure 4). Quinones then donate electrons to cytochrome *c* through the *bc*_1_ complex, again contributing to the pmf (Figure 15A). The resulting reduced cytochrome *c* serves as the redox-active partner for the nitrite reductase. Therefore, we expected the nitrite reductase to be especially up-regulated under heterotrophic conditions which allow to run the TCA [if *F. straubiae* is able to use organics from the culture]. Surprisingly, this seemed to be not the case—not even compared to the microoxic conditions. This observation may reflect a true physiological behavior for which we currently lack an explanation; alternatively, it could result from limitations in data quality, and the apparent trend may not be reliable.

Under autotrophic conditions, electrons are not donated by NADH [located in the cytosol], which would support energy conservation via transfer through NADH–quinone oxidoreductase and subsequently the *bc*_1_ complex [located in the inner membrane]. Instead, electrons originate from Fe(II) outside the cell. If electrons are transferred directly to cytochrome *c* upon entry into the periplasm, they cannot contribute to pmf generation at the inner membrane. Thus, any energetic gain from nitrite reduction must occur downstream using its product nitric oxide. Indeed, this appears to be the case, as we identified the long-standing missing nitric oxide reductase (NOR) of eNOR-family, which likely contributes to energy conservation by pumping protons into the periplasm.

#### 6.5.2 Discovery and classification of eNOR in F. straubiae!

Contrary to previous assumptions that *F. straubiae* did not have the capacity to reduce nitric oxide, we discovered two genes (2878407315 and 2878407316) that encode a nitric oxide reductase (eNOR-family)(Figure 16)(Murali *et al*. 2024). This is a significant finding, as it challenges the widely accepted hypothesis that the flanking community (2% bacteria other than *F. straubiae*) in culture KS (98% *F. straubiae*) is responsible for detoxifying NO (Huang, Straub, Blackwell *et al*. 2021).

**Figure 16:**
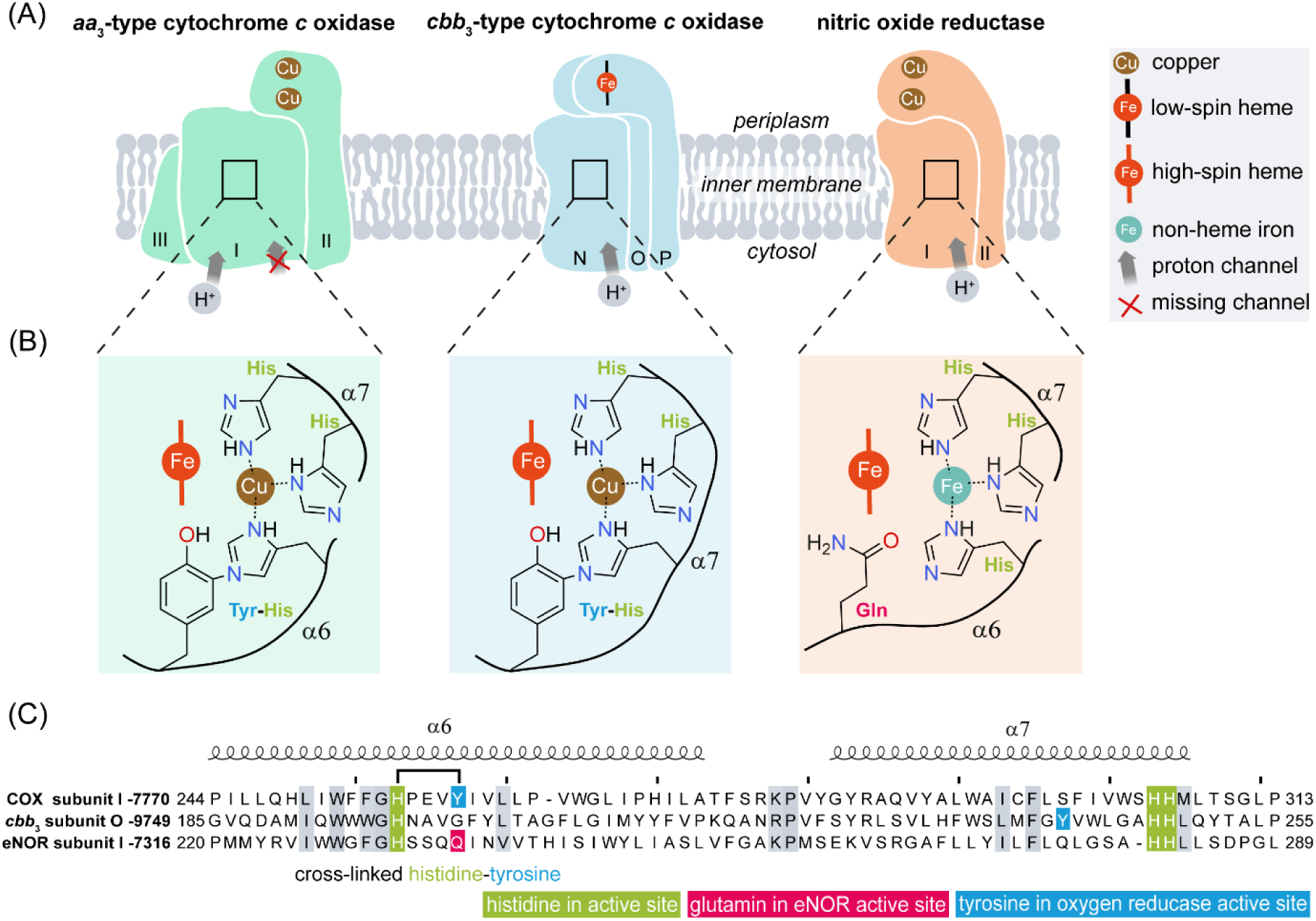
Comparison of the three heme-copper oxidoreductases (HCOs) in *F. straubiae*: cytochrome *c* oxidase (COX; A-family HCO), *cbb*_3_-type cytochrome *c* oxidase and nitric oxide reductase (eNOR-family)(IMG Gene ID: 2878407315 and 2878407316). (A) Schematic architecture showing cofactors and proton channel properties. The channel in eNOR is conserved within this family but has not yet been experimentally confirmed. This putative proton channel is highly similar to the K-proton channel of B- and A-family oxygen reductases. COX of *F. straubiae* is missing the conserved K-proton channel of the A-family. (B) Comparison of active sites. Oxygen reductases possess an active site composed of a high-spin heme, a redox-active cross-linked tyrosine, and a copper ion (Cu) ligated by three histidines. In contrast, the active sites of nitric oxide reductases consist of a high-spin heme and a non-heme iron (Fe) coordinated by three histidines and one glutamate, and lack the histidine–tyrosine cross-link. (C) Sequence alignment of the active-site regions. α: helix. Secondary structure was predicted by AlphaFold and visualized by ESPript 3.2.

**Figure 17:**
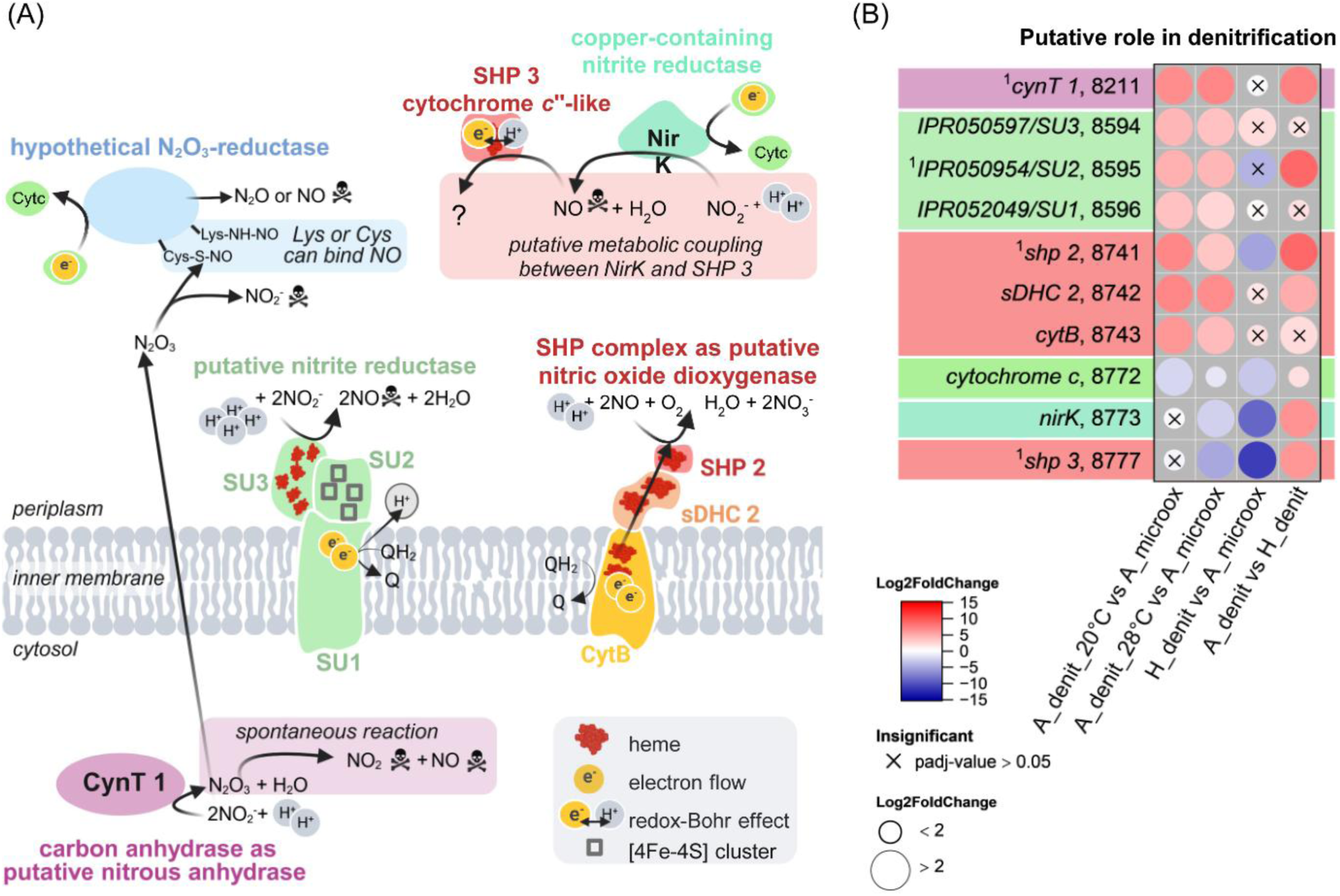
(A) Overview of novel (putative/hypothetical) denitrification pathways in *F. straubiae*. (B) The log_2_ fold changes of normalized transcript count (Log2FoldChange) are shown as colored dots. The labels follow the structure: gene name, last four numbers of IMG Gene ID. All IMG Gene IDs share the prefix 287840XXXX. The four conditions were: autotrophic denitrifying Fe(II) oxidation at 28°C (A_denit_28°C, n=3), heterotrophic denitrification at 28°C (H_denit, n=1), autotrophic denitrifying Fe(II) oxidation at 20°C (A_denit_20°C, n=2) and autotrophic microaerophilic Fe(II) oxidation at 20°C (A_microox, n=2). *F. straubiae* was grown as community member of culture KS in all nitrate containing conditions. ^1^no transcript detected in H_denit. Skull symbol: product is a cell toxin in too high concentrations. Cofactors are not indicated for all proteins. Gene abbreviation in table S7.

eNOR has two subunits: (i) eNOR subunit I, is a transmembrane protein binding to a high spin heme and a non-heme iron coordinated by three conserved histidine. Further, it possesses a K-channel that likely translocates H⁺ for active site chemistry, where N_2_O and one molecule of water are produced. By reducing the cytoplasm by two protons the eNOR is an energy conserving protein. (ii) eNOR subunit II has a transmembrane domain and a periplasmic domain with two coppers which reduces cytochrome *c*.

The *F. straubiae* eNOR operon encompasses: *eNOR II*, *eNOR I*, heme *o* synthase gene, *sco1*, and the eNOR scaffold protein gene of unknown function. This is a very common observation, however the two genes encoding proteins with partial similarity to *bc*_1_ complex chain C are rather unusual (2878407319 and 2878407323). The first gene encodes a protein encompassing a cytochrome *b*, [Fe-S] cluster, FAD-binding, and NADH-binding domain, suggesting a function as a quinol:NAD^+^ oxidoreductase. The second encodes a protein encompassing a cytochrome *b* and [Fe-S] cluster domain. Due to the close genetic proximity to eNOR, these proteins might assemble into a respiratory supercomplex were free energy from NO reduction is coupled to NAD^+^ reduction. All four proteins were present in the proteome extracted by Huang *et al*. (2021) from culture KS grown on autotrophic Fe(II)-oxidizing, denitrifying conditions (Table S6).

Most importantly, this operon is present in *F. straubiae* strain KS, *Ca.* F. altingense strain AG and *Ca.* F. bremense strain BP. If this complex truly supports the regeneration of NADH, the corresponding genes likely represent marker genes to identify *Ferrigenium* sp.-dominated autotrophic nitrate-reducing Fe(II)-oxidizing communities. Elucidating their function will be crucial for enhancing our current understanding of the underlying metabolic pathway.

Many eNOR operons encode a heme A synthase (CtaA homolog). In contrast, *F. straubiae* encodes two heme A synthases (Cox15 homologs) and thus likely does not require an additional copy within the eNOR operon. However, enzyme maturation represents a critical aspect that must be considered, as heme A biosynthesis is O_2_-dependent, whereas nitrate-reducing Fe(II) oxidation in this study was performed under anoxic conditions (Brown *et al*. 2002, Ji *et al*. 2015). This raises the question of how the required oxygen for eNOR maturation is supplied.

As mentioned previously, the proteomic data from the study of Huang *et al*. (2021) proofed that eNOR was present under Fe(II)-oxidizing anoxic conditions. Consequently, a source of oxygen must exist to enable heme A biosynthesis, the cofactor of eNOR subunit I. This may suggest that *F. straubiae* is capable of generating small amounts of oxygen, although this remains speculative.

Alternatively, the experimental conditions may not have been strictly anoxic. Indeed, observations from our laboratory and from Dr. Chao Peng (China West Normal University) indicate that culture KS does not grow under strictly anoxic conditions (personal communication). However, interpreting this is challenging, as oxygen reacts rapidly with Fe^2+^, and even trace amounts would be quickly consumed.

#### 6.5.3 Which role could a nitric oxide dioxygenase play in denitrification?

As discussed in section 6.4, the genes 2878408741–2878408743 (IMG gene ID) may encode a nitric oxide dioxygenase, while 2878408744 and 2878408745 likely represent associated regulatory components (a two-component system). The potential function of this putative nitric oxide dioxygenase is puzzling: it consumes a quinone and oxygen to produce nitrate, and it is strongly up-regulated under autotrophic denitrifying conditions. Obviously, this enzyme could serve as a NO-scavenging system, mitigating NO toxicity—but at a cost of ½ quinone per NO molecule and the uptake of one proton from the periplasm (leading to a decrease of the pmf). This reaction would regenerate nitrate for the nitrate reductase, which in turn contributes one more proton to the pmf than the dioxygenase consumes (Figure 15).

Overall, this process would basically reduce O_2_ to H_2_O, and *F. straubiae* possesses more energetically efficient pathways to achieve this. In any case the key question here is—where would the O_2_ come from? The biological function of the complex would need to be addressed by activity assays using the purified complex.

#### 6.5.4 Could one of the carbonic anhydrases be an isoenzyme functioning as nitrous anhydrase?

The gene *cynT1* (IMG IDs 2878408547), annotated as β-CA gene, was significantly upregulated under autotrophic denitrifying conditions (Figure 2: ’β-carbonic anhydrases’). This suggests a role in nitrate-reducing Fe(II) oxidation, which was not surprising based on the gene location next to the nitrate reductase, and suggests that it might even belong to the same operon and/or shares transcriptional regulators. We could not find literature on the synergy of carbonic anhydrase and nitrate reductase. Still, Andring *et al*. observed nitrite reductase and nitrous anhydrase activity for mammalian carbonic anhydrase II (α-carbonic anhydrase class II), which is structurally distinct to β-CAs (Andring, Kim, and McKenna 2020). However, both kinds are classically Zinc-carbonic anhydrases. When Andring *et al*. observed activity with nitrite, copper was bound to the carbonic-anhydrase, which was argued as cause for this observation. Despite the structural differences, it made us wonder, if the protein encoded by *cynT* 1 could also bind copper and react with nitrite. Consequently, it would function as isoenzyme of carbonic anhydrase. Further, carbonic anhydrase II was found to react with NO_2_^-^ and form N_2_O_3_ (Tsikas and Gambaryan 2021), which, in contrast to the nitrite reduction to NO, was attributed to its zinc-active site, a feature shared with the bacterial β-CA. Could the bacterial zinc-active site of β-CA also catalyze the reaction to N_2_O_3_? If so, does it dissociate into NO_2_ (cell toxin) and NO or could N_2_O_3_ be the substrate of a novel denitrification protein that reduces one of the nitrogen atoms to either NO or N_2_O? Alternatively, could this pathway even result in the formation of N_2_ and so-called dark O_2_ (oxygen production without light energy)?

Besides of CAs role in CO_2_ fixation pathways, or the speculated possibility as nitrous anhydrase, CAs are also known to function in cyanate degradation to ammonia (Supuran and Capasso 2017). Future research into the bacterial carbonic anhydrase might reveal even more novel functions, as such could be one in denitrifying nitrogen metabolism.

#### 6.5.5 Novel membrane-bound redox complex upregulated for nitrate-reducing Fe(II) oxidation

The transcripts of three adjacent genes showed some of the strongest up-regulation in the transcriptome under nitrate-reducing Fe(II)-oxidizing conditions (Figure 15C). The encoded protein complex consists of a membrane-bound subunit (SU1, Gene ID: 2878408596), an electron-transfer iron-sulfur cluster-binding protein (SU2, Gene ID: 2878408595), and a multiheme *c*-type cytochrome subunit (SU3, Gene ID: 2878408594, 6 heme binding motifs), as inferred from protein family assignments using InterProScan (Figure 18).

**Figure 18:**
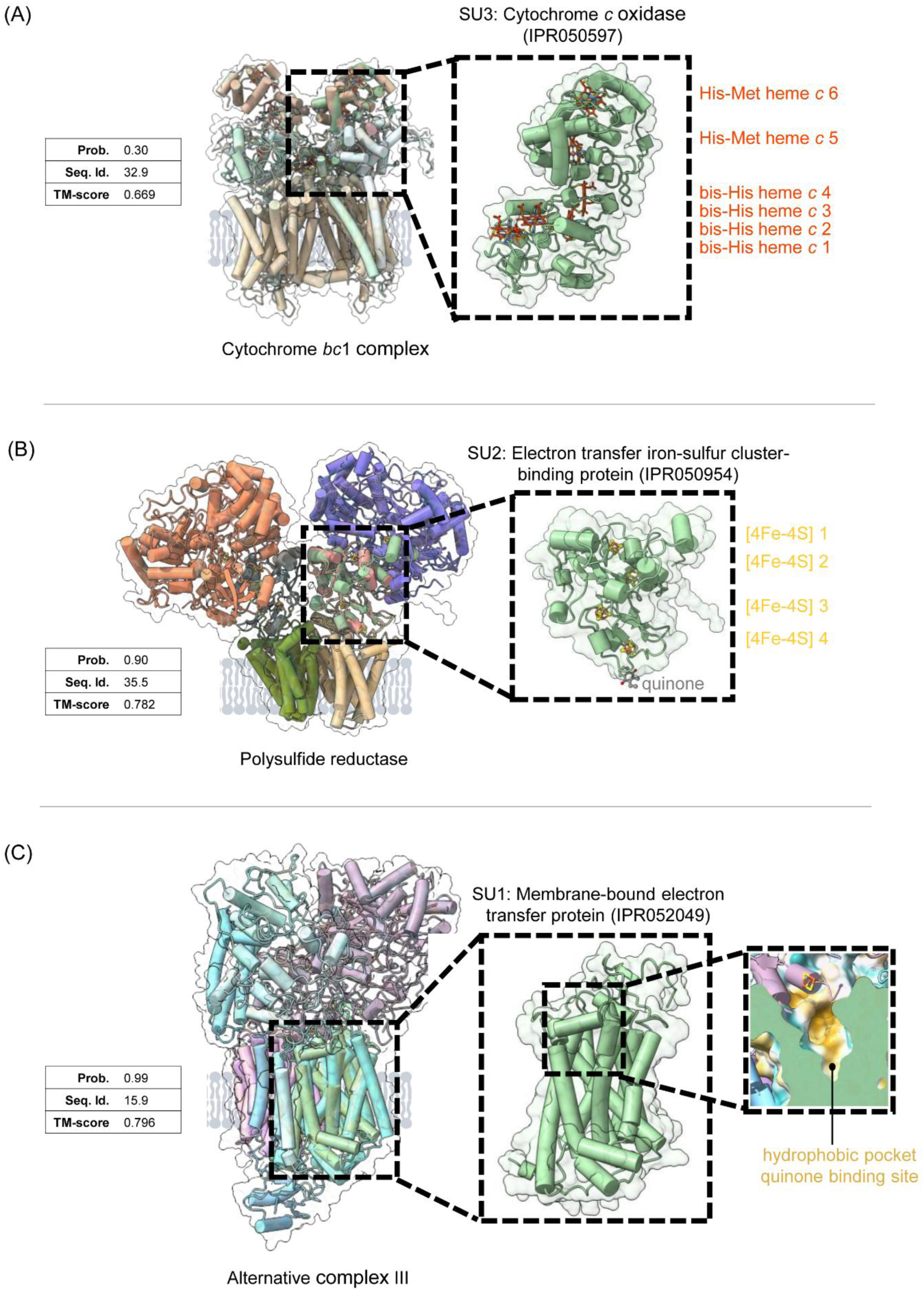
Novel membrane-bound redox complex upregulated for nitrate-reducing Fe(II) oxidation. Foldseek result showing the most similar structure hits to the Alphafold models of the proteins corresponding to Gene ID 2878408594 (A), 2878408595 (B) and 2878408596 (C) respectively. The protein family was determined by InterProScan. Prob. – Probability; Seq. Id. – Sequence Identity; TM-score – Template Modeling Score.

The AlphaFold model of SU1 reveals a putative quinone-binding pocket, similar to those found in the alternative complex III (ACIII) and polysulfide reductase (structural similarity identified using FoldSeek). SU2 aligns with the 4Fe-4S cluster-containing subunit of polysulfide reductase (inferred by FoldSeek), indicating the presence of four 4Fe-4S clusters, consistent with the feature predictions from InterProScan. SU3 contains four *c*-type hemes with axial bis-histidine ligation at the N-terminus, followed by two *c*-type hemes with His-Met axial ligands, as predicted by AlphaFold.

We propose that the overall architecture of this complex resembles that of alternative complex III, which facilitates the two-electron reduction of quinone by using the Fe-S cluster domain as an electron sink. Based on ProFunc analysis, a potential electron acceptor could be NO_2_^-^. While NO_2_^-^ becomes reduced by one electron to NO, the second electron harnessed from quinone "waits" in the Fe-S cluster for a second NO_2_^-^ to bind to the complex. However, this interpretation remains highly speculative at this stage (Figure 15A, putative nitrite reductase). Given the structural similarity of the membrane subunit to proton-translocating ACIII, we further propose a role for this complex in energy conservation via a redox-coupled proton translocation mechanism, making it one of the most compelling targets for future functional analyses.

### 6.6 Oxygen reduction by *cbb*_3_- and *aa*_3_-type cytochrome *c* oxidases

#### 6.6.1 The cbb_3_-type cytochrome c oxidase and an unconventional aa_3_-type cytochrome c missing a conserved proton pumping channel

*F. straubiae* has two types of oxygen-reducing, proton-pumping respiratory heme-copper oxidoreductase: a *cbb_3_*-type cytochrome *c* oxidase (Cbb_3_-COX) and an aa_3_-type cytochrome *c* oxidase (COX, complex IV) (architecture and active sites: Figure 16A; reaction scheme: Figure 19A). Further it does not possess a cytochrome *bd* oxidase. Having multiple oxygen reducing heme-copper oxidoreductases is typical for bacteria that need to adapt to different oxygen levels (Pitcher and Watmough 2004). While Cbb_3_-COX has a higher affinity for O_2_, allowing it to conserve energy even under low O_2_ partial pressure, the energy conservation efficiency of COX is higher under high O_2_ concentrations. Cbb_3_-COX is encoded by the classical operon structure *ccoNOQP* (gene IDs: 2878409749–2878409752), in which the gene products of *ccoN*, *ccoO*, and *ccoP* comprise the catalytic core complex (Kaplan *et al*. 2021). The COX is encoded by the operon *coxBA11C* (gene IDs 2878407768–2878407771), in which the gene products of *coxB*, *coxA*, and *coxC* comprise the catalytic core complex of COX (Esposti 2020). Interestingly, we observed that COX subunit I and COX subunit II lack the conserved K-channel of A-family heme-copper oxidoreductase. Mutations in the K-channel usually significantly impact the enzyme’s activity. The implementation of this maturation on its physiological function in *F. straubiae* COX remains unclear.

**Figure 19:**
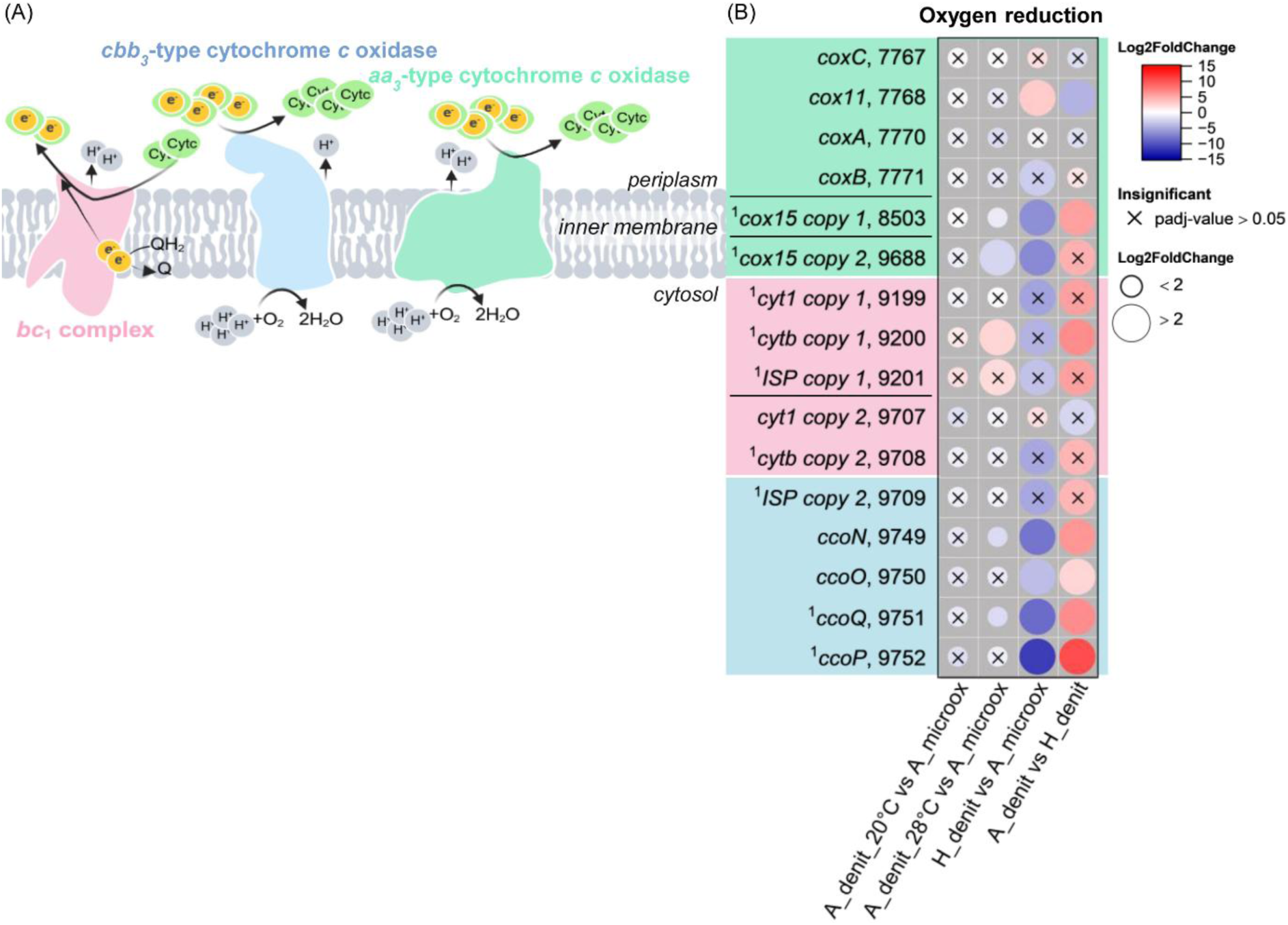
(A) Oxygen reduction pathway in *F. straubiae.* (B) The log_2_ fold changes of normalized transcript count (Log2FoldChange) are shown as colored dots. The labels follow the structure: gene name, last four numbers of IMG Gene ID. All IMG Gene IDs share the prefix 287840XXXX. The four conditions were: autotrophic denitrifying Fe(II) oxidation at 28°C (A_denit_28°C, n=3), heterotrophic denitrification at 28°C (H_denit, n=1), autotrophic denitrifying Fe(II) oxidation at 20°C (A_denit_20°C, n=2) and autotrophic microaerophilic Fe(II) oxidation at 20°C (A_microox, n=2). *F. straubiae* was grown as community member of culture KS in all nitrate-containing conditions. We found paralogues for the genes encoding *bc*_1_ complex and *coxA* and *B* and named them *copy 1* and *copy 2*. ^1^no transcript detected in H_denit. Gene abbreviation in table S7.

#### 6.6.2 Gene expression of oxygen and nitric oxide recuing heme-copper oxidoreductase in F. straubiae and its implementation of previous assumptions on the role of the flanking community in culture KS

Heme-copper oxidoreductases share a high degree of sequence homology; consequently, automated annotation pipelines often misinterpret their functions. In the case of *F. straubiae*, this has resulted in fatal errors and biased interpretations of previous ’-omics’ studies (He *et al*. 2016a, Huang, Straub, Blackwell *et al*. 2021). For example, it was erroneously assumed that the bacterium had second copies of coxA and coxB (the eNOR subunits I and II, with IMG gene IDs 2878407316 and 2878407315), which were ’surprisingly’ dominant under anoxic conditions. It also led to the erroneous conclusion that the flaking community must detoxify the cell toxin nitric oxide, as the nitric oxide reductase that it actually encoded remained unseen. One widespread assumption to explain the expression under anoxic conditions is that the flaking community might supply *F. straubiae* with oxygen (dark oxygen, formed without light and independently of phototrophic processes). Such a nitrate-driven dark oxygen production has been recently described by Zheng *et al*. (2025) (Zheng R *et al*. 2025). However, if this oxygen is not protected (for example, by a haemoglobin-like protein), it would react immediately with Fe(II) in the media and would not reach *F. straubiae*. Another widespread assumption is therefore that *F. straubiae* might possess a nitric oxide dismutase or another unknown protein capable of producing dark oxygen independently. However, these assumptions are now outdated due to the correct classification of the cox genes as eNOR genes. Nevertheless, no nitric oxide dismutase has been identified in the genome of *F. straubiae* (Becker and Kappler 2025).

We hope that this argument will disappear from the literature, as we have clarified that these genes encode the eNOR, the presence of which under anoxic conditions is not surprising. However, we would also like to state that we are not saying that oxygen formation as a secondary metabolite, [for example to facilitate heme A maturation] might not play a role within the community. Overall, it is more likely that the community’s role is in the production of a secondary metabolite than in the supply or detoxification of a primary metabolite.

The transcriptional data collected in this study and by Huang *et al*. (2021) both indicate the presence of heme-copper oxidoreductase under all three tested conditions, facilitating a rapid response by *F. straubiae* to changes in environmental oxygen and nitrate availability (normalized transcripts in TPM are listed in Table S2). Under autotrophic denitrifying conditions, transcripts of *eNORI/II* were the most abundant, followed by *ccoNOQP*, whereas the genes of the canonical *cox* operon showed the lowest expression levels. Under microoxic conditions, the expression pattern of *eNORI/II* and *ccoNOQP* was reversed, while the classical *cox* operon again remained the least expressed. The log_2_ fold change of *eNORI/II* under autotrophic denitrifying conditions was slightly higher (below two) than under microoxic conditions (Figure 15B), whereas *ccoNOQP* was slightly higher (below two) under microoxic conditions and oxygen reductases (Figure 19B).

The constitutive, low-level expression of the canonical *cox* operon under both anoxic and microoxic conditions enables *F. straubiae* to respond rapidly to a sudden shift in the environment to high oxygen availability. The dominance of *cbb*_3_-COX as an oxidase specialized for low O_2_ levels under microoxic conditions was likewise expected.

The presence of eNOR under microoxic conditions is not unusual and has been observed in anaerobic denitrifiers (Tamegai, Yamanaka, and Fukumori 1993, Tanimura and Fukumori 2000, Ji *et al*. 2015). This further supports the metabolic flexibility of *F. straubiae*. Additionally, nitric oxide (NO) can occur as a secondary metabolite; thus, eNOR may not only function as an enzyme of primary metabolism but also play a broader role in NO detoxification.

### 6.7 Iron-related genes involved in transcription regulation, iron acquisition and storage

We identified Fe-related genes involved in gene regulation (Figure 20A) as well as iron acquisition and storage (Figure 20B) using the software tool FeGenie. We found that the Fe starvation-associated sigma factor transcription regulator *pvdS* (IMG Gene ID: 2878408770) was upregulated under autotrophic denitrifying conditions compared to the autotrophic microoxic condition. This is surprising, as approximately 7.9 mM Fe(II)_total_ [2.2 mM dissolved Fe(II)] was present in the medium at the sampling time point and could have been further oxidized [evidenced by cultures which were continuously sampled and not harvested for RNA extraction] (Figure S2) (no Fe concentration was measured for the microoxic conditions). Furthermore, the expression of this Fe starvation transcription regulator was slightly higher under heterotrophic denitrifying conditions, which did not include Fe, and is therefore expected.

**Figure 20:**
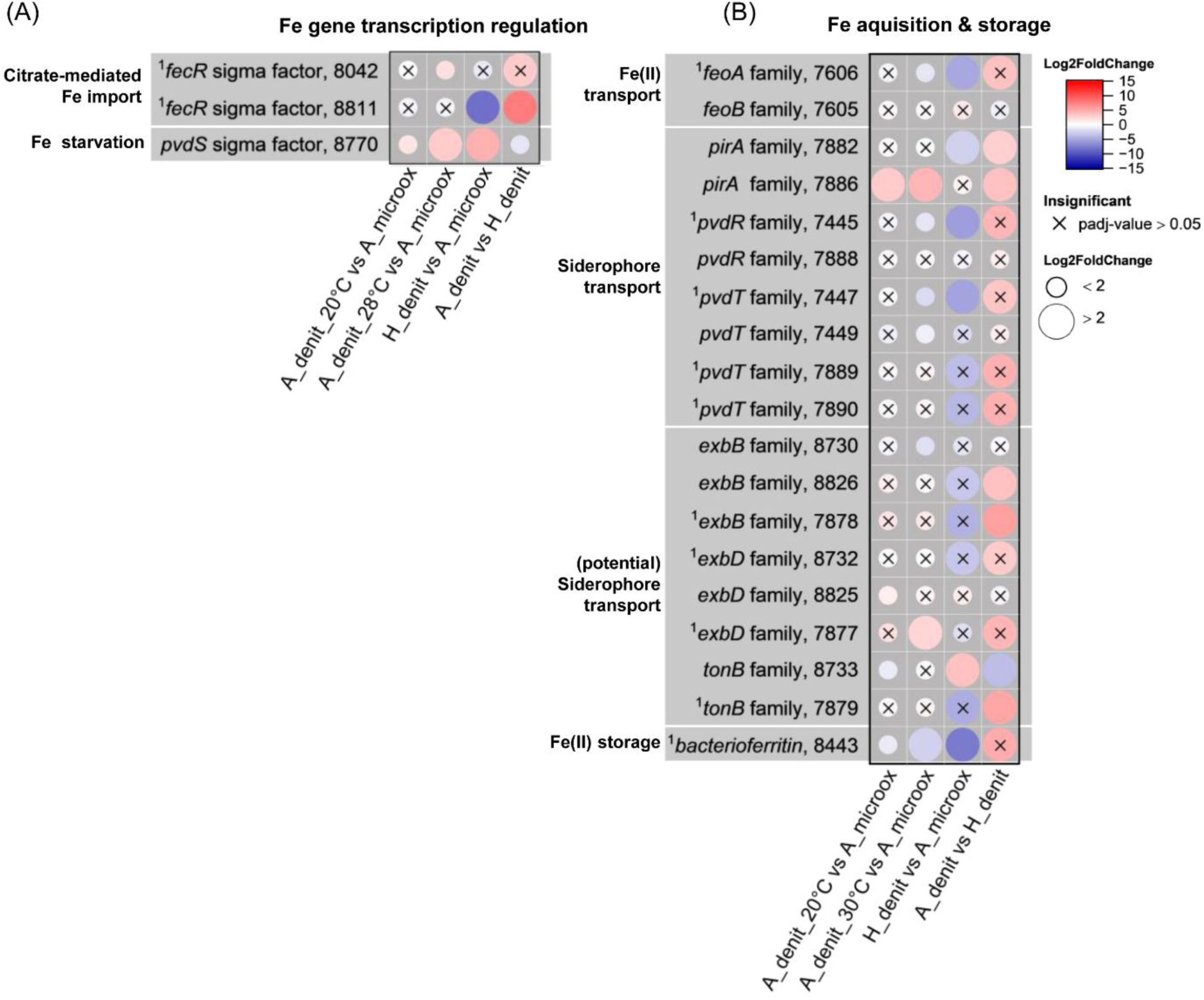
Fe-metabolism related genes identified by FeGenie in *F. straubiae*. The log_2_ fold changes of normalized transcript count (Log2FoldChange) are shown as colored dots. The labels follow the structure: gene name, last four numbers of IMG Gene ID. All IMG Gene IDs share the prefix 287840XXXX. The four conditions were: autotrophic denitrifying Fe(II) oxidation at 28°C (A_denit_28°C, n=3), heterotrophic denitrification at 28°C (H_denit, n=1), autotrophic denitrifying Fe(II) oxidation at 20°C (A_denit_20°C, n=2) and autotrophic microaerophilic Fe(II) oxidation at 20°C (A_microox, n=2). *F. straubiae* was grown as community member of culture KS in all nitrate-containing conditions. 1no transcript detected in H_denit. Gene abbreviation in table S7.

Analysis of Fe acquisition genes revealed that the siderophore receptor gene *pirA* (IMG Gene ID: 2878407886) was strongly upregulated during nitrate-reducing Fe(II) oxidation (autotrophic denitrification). This indicates a higher Fe demand for biosynthesis compared to the other respiratory pathways underlying autotrophic microaerophilic Fe(II) oxidation and heterotrophic denitrification. In contrast, bacterioferritin, which is involved in Fe(II) storage, was upregulated under autotrophic microoxic conditions. This suggests that Fe(II) released from FeS minerals may be more bioavailable than the 2.2 mM dissolved Fe(II) present under autotrophic denitrifying conditions. The more iron available the more likely it is that *F. straubiae* would invest in bacterioferritin production to store excess Fe(II).

## 7. Conclusions and outlook

In our comprehensive genome and transcriptome analysis of *F. straubiae* under autotrophic denitrifying (nitrate-reducing Fe(II) oxidation), autotrophic microoxic (microaerophilic Fe(II) oxidation), and heterotrophic denitrifying (heterotrophic denitrification) conditions, we aimed to identify research targets to shed light on the poorly understood physiology of *F. straubiae*.

The transcriptomic data indicate that the CBB cycle was most active under autotrophic conditions; however, it was also present under heterotrophic conditions, raising the question of whether *F. straubiae* can grow on acetate at all. Furthermore, we identified the CBB cycle as the sedoheptulose-7-phosphate-forming transaldolase variant, which appears to function as the initial CO_2_ fixation pathway feeding into a roTCA. Whether this reversed TCA operates as a complete cycle or terminates at citrate remains unresolved and could be addressed using the protocols described by Steffens *et al*. for tracking the roTCA in bacteria (Steffens *et al*. 2022).

In addition to the CBB cycle in combination with the reversed TCA cycle, we hypothesize the existence of an unknown CO_2_ fixation pathway feeding the glycine synthase pathway via 10-formyl-THF. Experiments using labeled CO_2_ could help to elucidate this possibility.

Under autotrophic conditions, the electrons required for CO_2_ fixation and for maintaining cellular energy homeostasis originate from Fe(II) oxidation. Among the putative Fe(II) oxidases, we showed that Cyc2 is the most dominant Fe(II) oxidase. Sequence analyses suggest that (i) Cyc2 may function as a trimer, (ii) MtoA possesses less common His/Met heme coordination, leading to higher reducing potentials compared to classical bis-His coordination, and (iii) MofA contains FN domains that may function as adhesive element like a sticky arm to facilitate interactions with Fe-bearing minerals.

It is puzzling that *F. straubiae* encodes three potential Fe(II) oxidases, and we speculate that each may serve a specific biological function. These functions could include forming a supercomplex within the respiratory chain for O_2_ reduction to H_2_O, coupling to denitrification, or contributing to quinone pool reduction to support CO_2_ fixation. In such a scenario, the bacterium may be able to harness electrons from Fe(II) species with different reducing potentials.

We screened the entire genome for cytochromes which we all addressed in this comprehensive study. Thereby we discovered (i) that SHP-like protein complexes play an important role in *F. straubiae* (IMG gene IDs: 2878407668–2878407670; 2878408741–2878408743 and 2878408777), (ii) the presence of a putative octaheme complex within a cytochrome-rich gene neighborhood encoding in total ten heme-binding proteins (IMG gene IDs: 2878409853–2878409864) and a conserved nonaheme cytochrome *c* (28784098774) that may form an electron transfer chain within the periplasmic space to link Fe(II) oxidation to quinone reduction, (iii) a potentially proton-translocating redox complex that appears to be specific for nitrate-reducing Fe(II) oxidation (IMG gene IDs: 2878408594–2878408596).

To elucidate the exact roles of these complexes, further studies will be required. These include substrate identification by mass spectrometry, protein structural analyses using cross-linking experiments, pull-down-based interaction analyses, classical single-particle cryo-electron microscopy and protein crystallography, as well as cryo-electron tomography, which may provide insights into supercomplexes spanning the periplasmic space, the spatial distribution of Fe(II) oxidases at the outer membrane, and cellular attachment to Fe(II)-bearing minerals. In addition, these complexes will need to be extracted from their native environment—ideally produced in their original host organism—to preserve complex architecture and protein-protein interactions. Such native complexes should then be analyzed using cryo-electron microscopy and native mass spectrometry.

A key aspect of our study is the recognition that the heme-copper oxidoreductases 2878407315 and 2878407316 represent a nitric oxide reductase (eNOR-family). While previous studies discussed the possibility that *F. straubiae* encodes a novel NO dismutase or reductase, eNOR was present in the genome all along, but its function remained unrecognized due to misannotation as an oxygen reductase, effectively “hiding in plain sight.” This finding not only challenges the previous assumption that nitric oxide detoxification in culture KS is performed by the flanking community members, but also identifies a denitrification step that may contribute to the proton motive force. Furthermore, eNOR was found within an operon that likely encodes a novel quinol:NAD+ oxidoreductase (2878407319).These genes may function as maker genes to identify *Ferrigenium* sp.-dominated autotrophic nitrate-reducing Fe(II) oxidizing communities.

Last but not least, we would like to highlight that the fate of N intermediates shall be not explained by canonical denitrification genes alone, while several genes are highly expressed under denitrifying conditions without a known function (Figure 15C). Such as the SHP-like protein complex encoded by 2878408741–2878408743 (IMG Gene ID) and the redox complex encoded by 2878408594–2878408596 which is likely electrogenic, conserving energy from N-species reduction by translocating protons across the membrane creating an electrochemical gradient. Further we hypothesize that the carbon anhydrase encoded by 2878408211 could function as an isoenzyme interacting with nitrite or nitric oxide. We therefore propose that nitrate reduction under Fe(II)-oxidizing conditions involves this set of previously unrecognized proteins and is more complex than the canonical denitrification pathway.

## Supporting information

Supplemental Transcriptomic Data Table

## 8. Author statements

### 8.1 Author contributions

S.B. (Conceptualization, Data curation, Formal analysis, Methodology, Supervision, Visualization, Writing – original draft, Writing –review& editing), D.S. (Software, Formal analysis, Validation, Writing –review& editing), J.H. (Bioinformatic analysis of heme-copper oxidoreductase), (A.K. (Resources, Proofreading –review& editing). S.B. conceived and planned the study. S.B. performed all wet-lab experiments. S.B. conducted all bioinformatic analyses, except for transcriptomic raw data processing and validation, which were performed by D.S., and the bioinformatic analysis of the heme-copper oxidoreductase, which was carried out by J.H.. D.S. managed data storage and availability. S.B. illustrated and wrote the manuscript with input from D.S. and J.H.. D.S. and A.K. proofread and edited the manuscript.

## 8.4 Acknowledgements

We would like to acknowledge the Core Facility Genomics, Medical Faculty, University Hospital Tübingen for performing NGS methods and the Quantitative Biology Center (QBiC) for data management and storage of raw data. We thank Lars Grimm and Dr. Ulf Lueder for synthesizing the FeS used in this study. We thank Tzu Jung Huang for carrying out the chloramphenicol step of the Lueders protocol and carrying out all test extractions. We are grateful to Prof. Dr. Clara Chan, Dr. Jessica Keffer, Dr. Arkadiy Garber, Dr. Cristina Escudero and Prof. Dr. Karl Forchhammer for their feedback on the manuscript. We acknowledge the use of AI-assisted and non-AI language tools, including OpenAI’s ChatGPT (GPT-4 and GPT-5), DeepL Pro Write (2025), and Microsoft Word Professional Plus 2019, which were used solely to improve grammar, spelling, and readability. The outputs of AI-assisted software tools were critically reviewed to ensure that language improvements did not alter the intended meaning of the original content. We thank the Max Planck Society for providing the DeepL software license. We acknowledge the use of Biorender, ChimeraX, ChemDraw, Geneious Prime, PowerPoint, Adobe Illustrator, and OriginPro v10.15 for data visualization and figure generation. We like to thank OriginLab for a learning license and their User Support which scripted the template used to generate the bubble-plots.

