## Supplemental Text for "Towards understanding nitrate-reducing Fe(II) oxidation in *Ferrigenium straubiae*: spotlight on old and new key protein candidates"

### SI-Figure 1: Cultures and intermediate steps of RNA purification for transcriptomic study

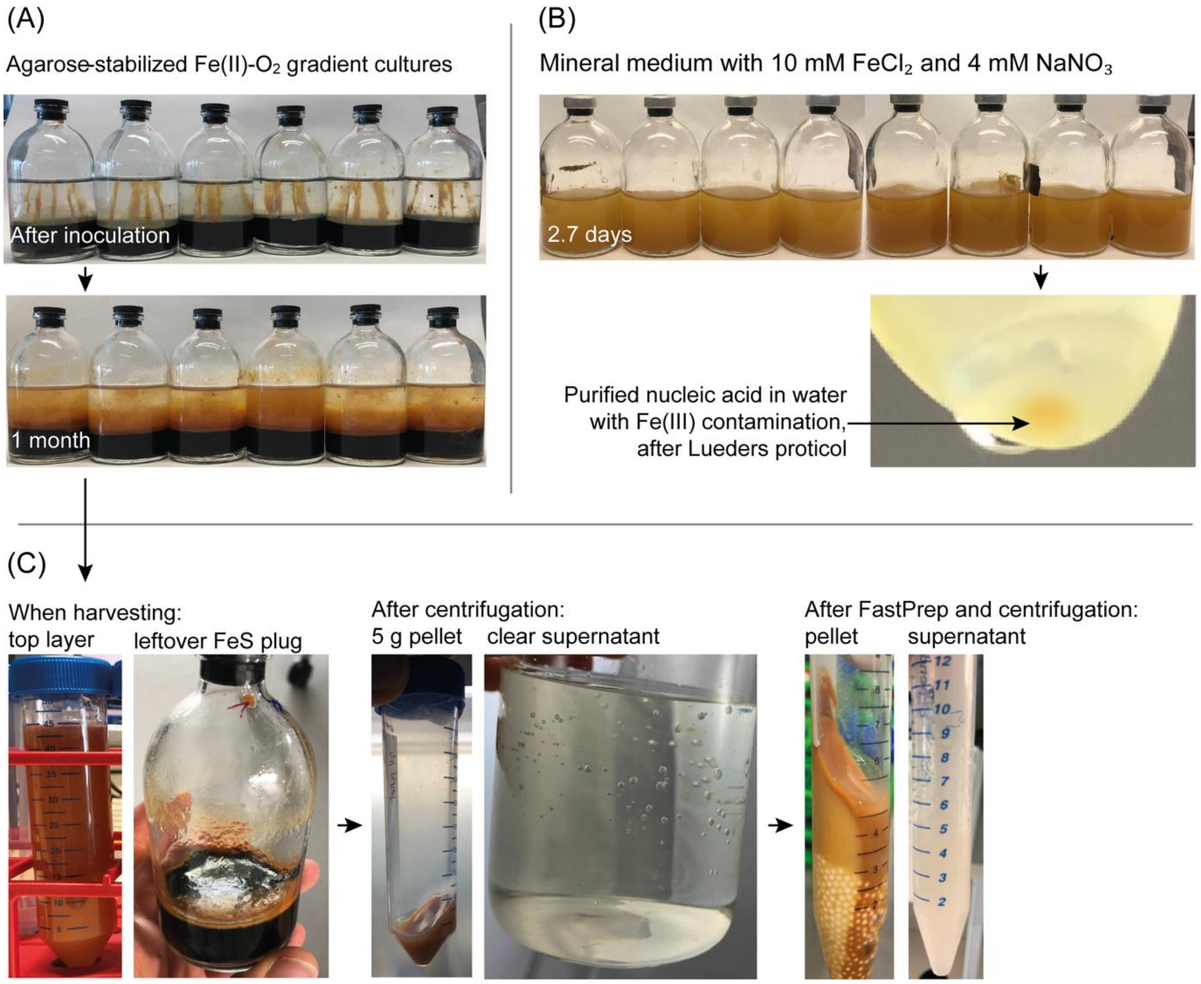

**Figure S1: (A) and (B): Iron-rich *F. straubiae* cultures used for RNA isolation. The fourth one-month-old gradient serum bottle accidentally tipped over during imaging. (C): Intermediate samples collected during nucleic acid isolation using the Lueders protocol (Lueders, Manefield, and Friedrich 2004). Arrows indicate the conditions corresponding to the intermediate samples.**

### SI-Figure 2: Nitrate-reducing Fe(II) oxidizing *F. straubiae*

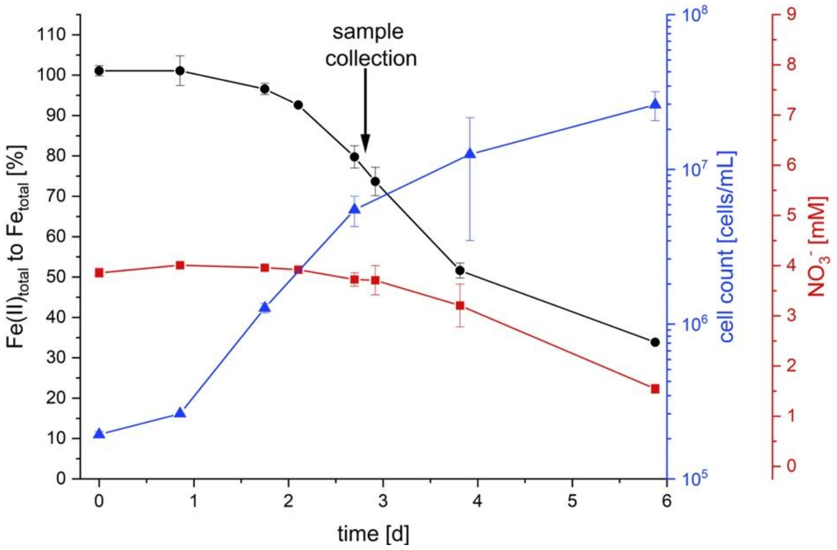

**Figure S2: Fe(II), cell and nitrate concentration of *F. straubiae* cultures in unfiltered mineral medium buffered with 22 mM bicarbonate (pH 6.4) and supplemented with 10 mM $FeCl_2$ and 4 mM $NaNO_3$. The cultures were inoculated together with the cultures that have been harvested for the transcriptomic study. The 8 bottles for RNA isolation were harvested at 2.7 days (late exponential phase), as indicated by the arrow. n =2. Nitrate and Fe(II) were analyzed as described in Becker *et al.* and the cell count was performed as described in Becker and Kappler *et al.* (Becker and Kappler 2025; Becker *et al.* 2025).**

**SI-Text 1: Cell viability check by fluorescence microscopy**

Fluorescence microscopy was performed using a Leica CTR5500 Microscope (Leica Microsystems GmbH) equipped with the Leica Application Suite X software.

Propidium iodide (PI) staining was conducted using the LIVE/DEAD™ BacLight™ Bacterial Viability Kit (Invitrogen, Thermo Fisher Scientific, Inc.) according to the manufacturer's instructions. This kit contains both PI and SYTO 9. While SYTO 9 is theoretically capable of staining both live and dead cells, in practice it does not efficiently stain viable *F. straubiae* cells and is inactive for dead cells when used within this dual-stain system.

For independent SYTO 9 staining (SYTO™ Green Fluorescent Nucleic Acid Stain SYTO 9, Invitrogen, Thermo Fisher Scientific, Inc.), a 1:4 working solution in DMSO was freshly prepared from a frozen stock. The working solution was used to stain culture samples at a 1:1000 dilution, resulting in a final staining concentration of 1.25 µM.

For SYTOX Green staining (SYTOX™ Green, Invitrogen, Thermo Fisher Scientific, Inc.), a 1:1000 working solution in Milli-Q water was prepared and stored at 4°C. The working solution was used to stain culture samples at a 1:1 dilution, resulting in a final staining concentration of 2.5 µM.

To assess cell viability, cultures were imaged under two conditions: (i) untreated samples directly after collection and (ii) samples after heat treatment at 70°C for 5 minutes. This approach was necessary because viable *F. straubiae* cells exhibit poor SYTO 9 staining, despite SYTO 9 being theoretically capable of labeling all intact and disrupted cells. Therefore, cell integrity was evaluated by comparing intact and partially disrupted cells (untreated) with killed/fully disrupted cells (heat-treated).

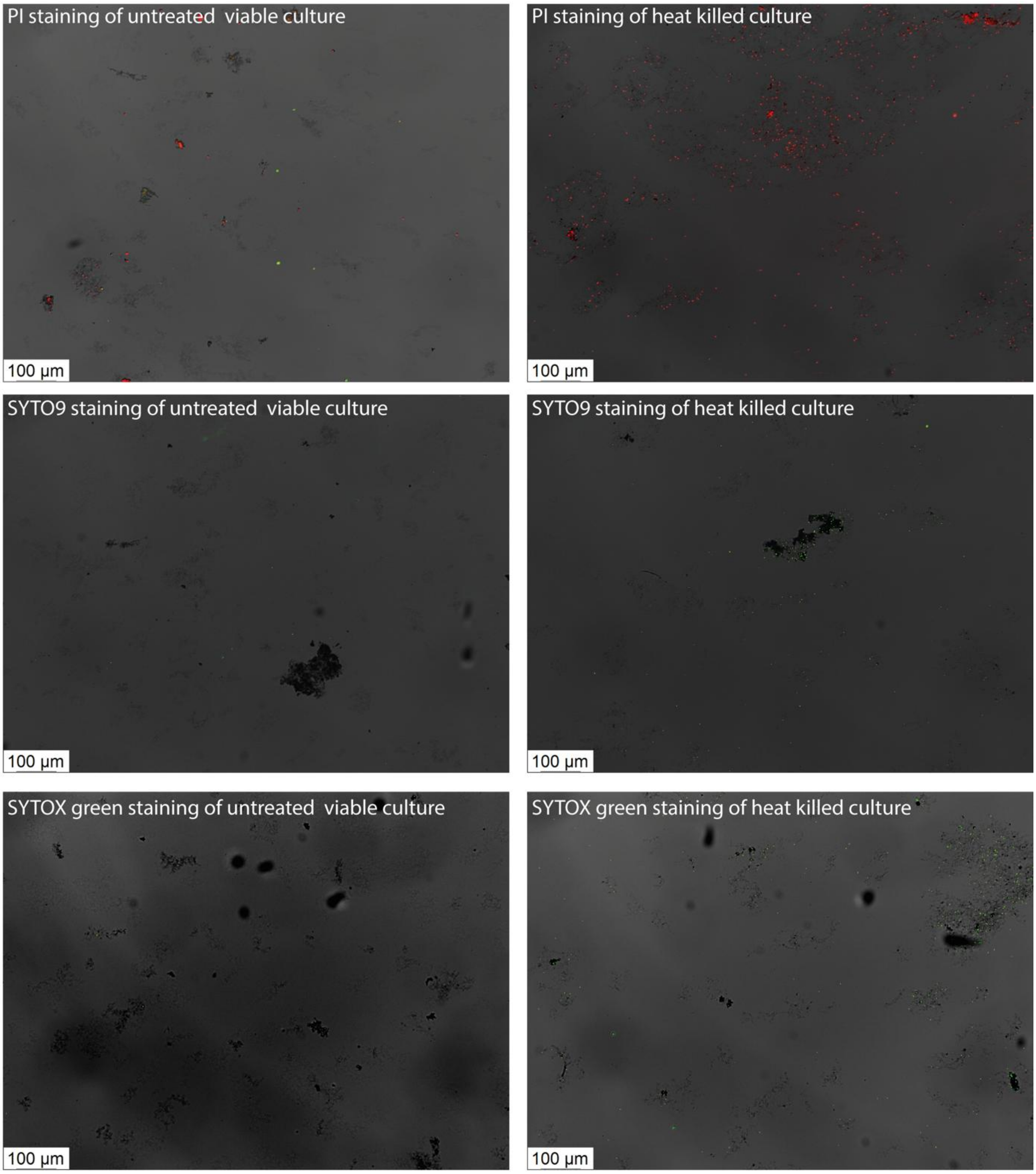

**Figure S3: Assessment of cell viability in *F. straubiae* cultures. Cultures were imaged under two conditions: (i) untreated samples immediately after collection and (ii) samples subjected to heat treatment at 70°C for 5 minutes. The figure in the left column display only dead cells in a viable *F. straubiae* culture, whereas the right column shows all cells following heat-induced cell death. The results confirm that the *F. straubiae* culture was healthy and predominantly viable.**

**SI-Figure 4: Volcano plots comparing the transcriptome of autotrophic denitrifying with autotrophic microaerophilic Fe(II) oxidation**

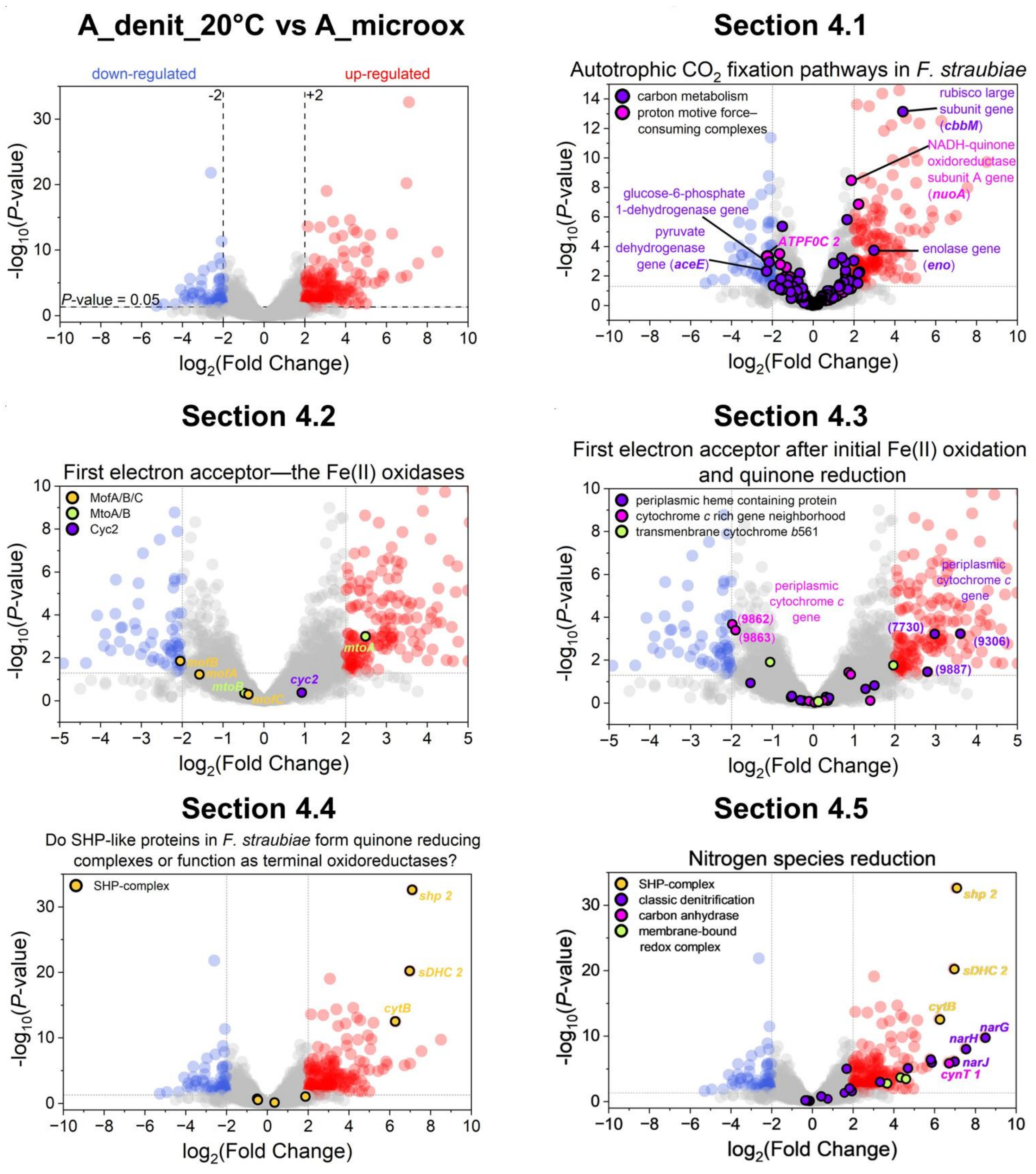

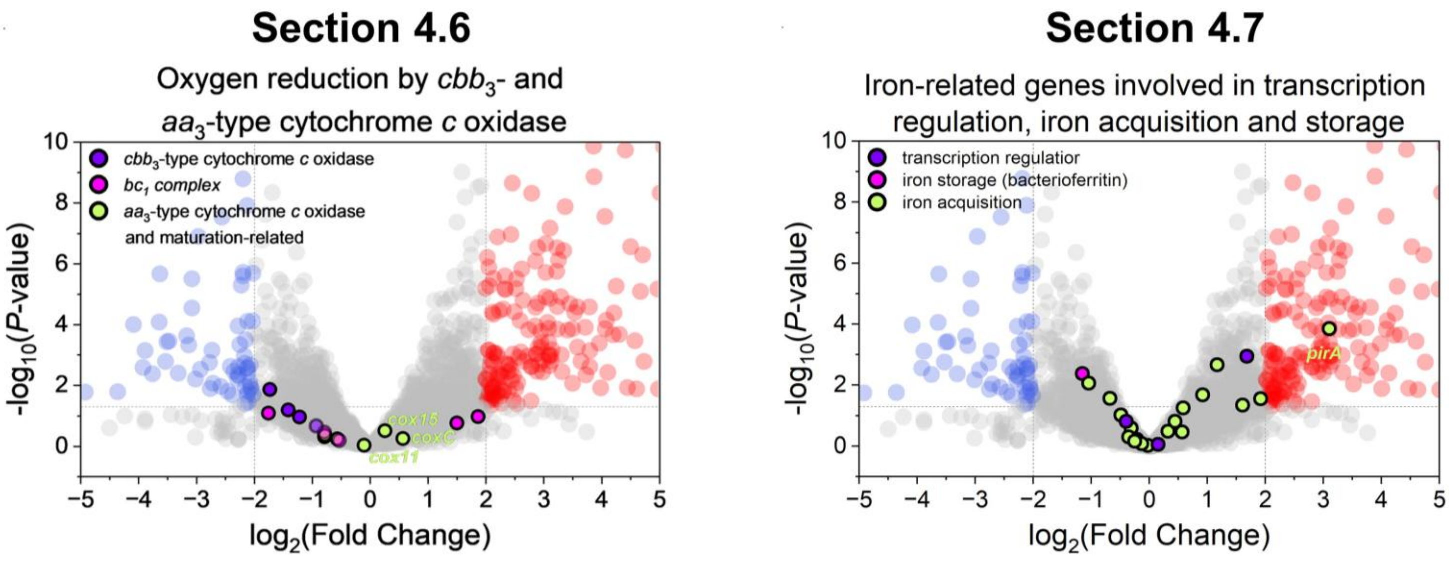

**Figure S4: Volcano plots comparing the transcriptome of autotrophic denitrifying (A_denit_20°C) with autotrophic microaerophilic (A_microox) cultivated *F. straubiae* cultures. The volcano plots serve as a table of contents, highlighting proteins that are discussed in the corresponding sections indicated by the plot titles. Please note that the title refers to the title of the section and does not necessarily describe what is shown in the plot. The plots include the genes that are the subject of the corresponding sections. All the plots show the same data in the background (blue, red, gray), while the transcripts of interest in the corresponding the sections were highlighted. The separation into individual plots and variation in axe scaling was done to avoid overlaps.**

**SI-Text 2: Computational modeling of the multicopper oxidase MofA of *F. straubiae* indicates potential role in mineral attachment**

$MofA_{KS}$ is a multicopper oxidase (MCO) with four copper binding sites (Tang *et al.* 2025) and it is homologous to a putative manganese oxidase of the Mn(II)-oxidizing bacterium *Leptothrix discophora* (Corstjens *et al.* 1997) and OmpB of *G. sulfurreducens* which possesses Fe(III) reduction activity (Mehta *et al.* 2006). It has been suggested that homologs of $OmpB_{G.\ sulfurreducens}$ and $MofA_{Leptothrix\ discophora}$ present in Fe(II)-oxidizing bacteria could potentially function as Fe(II) oxidase (He *et al.* 2017). He *et al.* (2017) considered MofA homolog of *F. straubiae* ($MofA_{KS}$) as a potential Fe(II) oxidase. However, at this stage it is very speculative. In *Leptothrix discophora mofA* forms an operon with *mofB* and *mofC*, similarly to *F. straubiae*, where homologs to *mofB* and *mofC* could be identified as well. MofB, is proposed to be a chaperone involved in the folding and activation of MofA, and MofC (G. J. Brouwers 2000) MofC is a cytochrome *c* type protein, which likely functions in concert with MofA to oxidize either Mn(II) in *Leptothrix discophora* or Fe(II) in *F. straubiae*.

While *mofA* expression was up-regulated in both microoxic and denitrifying Fe(II)-containing conditions compared to the heterotrophic condition without Fe(II), mofB and C transcripts were not even detected without Fe(II) (Table S 2). This supports its potential function as Fe(II) oxidase.

For a better understanding of $MofA_{KS}$, we performed sequence analysis and predicted the structural model. $MofA_{KS}$ has four Cu binding motifs [two HXH, HXXHXH and HCHXXXH] thus it binds to four copper ions (Gräff *et al.* 2020; Tang *et al.* 2025). Further it has two Fe(III)-binding motifs (EXXE) similar to $OmpB_{G.\ sulfurreducens}$ (Mehta *et al.* 2006).The model of $MofA_{KS}$ derived from an alphafold prediction (Abramson *et al.* 2024) shows five beta sandwich domains (Figure S5A) which are similar to the two beta sandwich domains of the extracellular subunit (MtrC) of the Fe(II) reductase complex Mtr of *Shewanella baltica* OS185 (Figure S5B) (Edwards *et al.* 2020) and the fibronectin type III (FNIII) domain (Figure S5C) (Pak *et al.* 2020). Based on this similarity the five beta sandwich domains of $MofA_{KS}$ are potentially involved in cell adhesion, and thus might be involved in attachment of MofA to the cell envelope, to iron minerals, other minerals and/or to proteins such as MofC which is a heme *c* binding protein encoded in near gene neighborhood. The 3D location of the $BSS2\text{-}3_{MofA}$ domains differ significantly among different alphafold models and a model which we derived by using I-Tasser structural prediction. Consequently, the model shown in Figure S5 only illustrates the existence of these domains. However, the true location of BSS2-5 is not reflected. There is also the possibility that the linker sequences of individual BSS domains are flexible anyway.

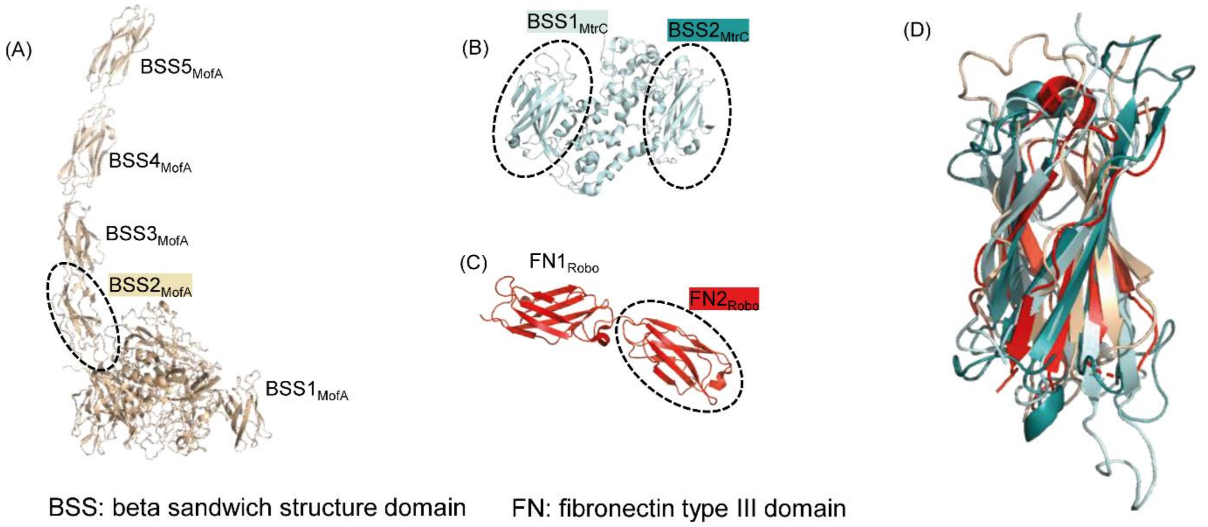

**Figure S5: Presence of beta sandwich structure domains (BSS) in metal-oxidizing MofAKS (A) and -reducing MtrC (B) proteins and their similarity to fibronectin type III (FN) domain. As an illustrative example, we present here two fibronectin type III domains from the transmembrane receptor Robo3 (C). The structures of BSS2MofA, BSS1MtrC, BSS2MtrC and FN2Robo are superposed and shown in (D). The structure of MofAKS is an alphafold model, MtrC and the FN domains of Robo3 are crystal structures (pdb ID: 6R2Q and 6POK)(Edwards *et al.* 2020; Pak *et al.* 2020). The structures were aligned by Pairwise Structure Alignment (Bittrich *et al.* 2024).**

Based on its function as either Mn or (potentially) Fe(II) oxidase, the absence of predicted transmembrane regions, and the signal peptide Sec/SPI in the MofA and C protein sequences, MofAC is likely located extracellularly [on the outside of the cell] (Viklund and Elofsson 2004; Nielsen *et al.* 2024). This raises the question of how the electron passes the outer membrane of if the electron derived from Fe(II) directly reduces $O_2$? In the latter case it would be rather involved in detoxification than energy metabolism Does $MofAC_{KS}$ potentially pass the harnessed electrons to Cyc2 or MtoAB? To answer this questions, Fe(II)-oxidation potential of $MofAC_{KS}$ needs to be studied and further its potential interaction with Cyc2, MtoA/B or other electron accepting cytochromes.

**SI-Text 3: Genomic screening for an alternative multicopper Fe(II) oxidase to MofA in *F. straubiae* could not identify a potential candidate**

We aimed to identify additional MCOs in *F. straubiae* similar to MofA that may function as potential Fe(II) oxidases. Candidate proteins were required to meet the following three criteria: (i) be identified as multicopper oxidase based on the presence of known amino acid motifs involved in copper binding (Gräff *et al.* 2020; Tothero *et al.* 2024); (ii) be upregulated under Fe(II)-oxidizing conditions; and (iii) be localized outside the cytosol.

The search for multicopper oxidase (MCO) amino acid motifs in the *F. straubiae* genome revealed 6 MCOs of unknown function, MofA, and 18 MCOs with specific protein designations assigned by the IMG genome annotation pipeline (Figure S6). If these annotations are correct, the specific functions of the 18 annotated MCOs are already known. We did not identify any novel potential Fe(II) oxidases, as the MCOs of unknown function that were specifically expressed under one or two Fe(II)-containing conditions (IMG IDs: 2878408028, 2878409595, and 2878408610) were predicted to be cytoplasmic proteins (Nielsen *et al.* 2024), and therefore did not meet the criteria for potential Fe(II) oxidases.

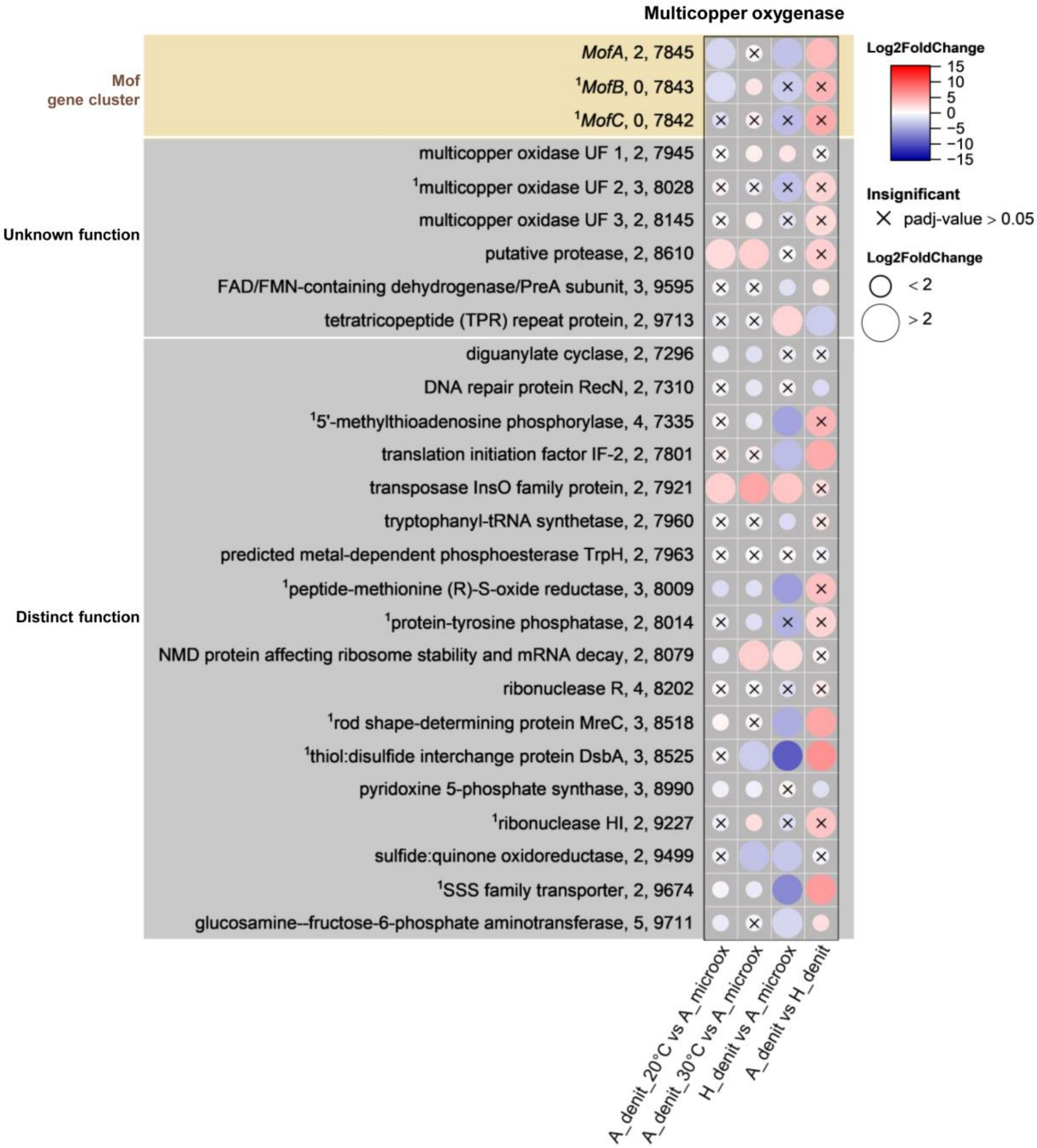

**Figure S6: Multicopper oxidases in *F. straubiae* and corresponding fold changes of normalized counts (Log2FoldChange) of transcripts. The label of the y-axis follows the following structure: gene product, number of Cu-binding amino acid motifs, last four numbers of IMG Gene ID. All IMG Gene IDs are of the structure 287840XXXX. The set of MCO motifs was published in Gräff *et al.* and a custom the script for identification of the motifs was developed initially coded by Tothero *et al.* (Gräff *et al.* 2020; Tothero *et al.* 2024). The four conditions were: autotrophic denitrifying Fe(II) oxidation at 28°C (A_denit_28°C, n=3), heterotrophic denitrification at 28°C (H_denit, n=1), autotrophic denitrifying Fe(II) oxidation at 20°C (A_denit_20°C, n=2) and autotrophic microaerophilic Fe(II) oxidation at 20°C (A_microox, n=2). *F. straubiae* was grown as community member of culture KS in all nitrate-containing conditions. [1]no transcript detected in H_denit; UF: unknown function.**

### SI-Table 1 and 2: Normalized transcripts of genes encoding putative Fe(II) oxidases and Log2FoldChange of Fe(II) oxidases within one condition compared to each other

**Table S 1: Normalized transcripts in TPM (Transcripts Per Million) of genes encoding putative Fe(II) oxidases, Mn-oxidase MofA and MofA associated proteins, in *F. straubiae*. The four investigated conditions were: autotrophic denitrifying Fe(II) oxidation at 28°C (A_denit_28°C, n=3), heterotrophic denitrification at 28°C (H_denit, n=1), autotrophic denitrifying Fe(II) oxidation at 20°C (A_denit_20°C, n=2) and autotrophic microaerophilic Fe(II) oxidation at 20°C (A_microox, n=2).**

| IMG Gene ID | Gene name | Normalized transcripts in TPM | | | | | | | |
|---|---|---|---|---|---|---|---|---|---|
| | | A_microox 1 | A_microox 2 | A_denit_20°C 1 | A_denit_20°C 2 | A_denit_28°C 1 | A_denit_28°C 2 | A_denit_28°C 3 | H_denit_28°C 1 |
| 2878409868 | *cyc2* | 2237.93 | 881.76 | 60633.36 | 480.34 | 17951.51 | 15850.5 | 14550.58 | 2122.91 |
| 2878407665 | *mtoA* | 0.29 | 0.12 | 9.17 | 3.84 | 23.37 | 53.23 | 65.57 | 0 |
| 2878407666 | *mtoB* | 3.42 | 1.88 | 11.5 | 7.42 | 20.86 | 50.49 | 64.02 | 521.21 |
| 2878407845 | *mofA* | 6.13 | 2.7 | 1.29 | 4.8 | 221.46 | 102.64 | 243.2 | 33.05 |
| 2878407843 | *mofB* | 1.69 | 0.83 | 1.21 | 1.52 | 117.55 | 82.03 | 132.21 | 0 |
| 2878407842 | *mofC* | 1.32 | 0.57 | 0.94 | 1.47 | 49.82 | 48.39 | 86.03 | 0 |

**Table S 2: Log2FoldChange of Fe(II) oxidase to each other within one condition.**

**A_micoox**

| | MtoA | MofA |
|---|---|---|
| MtoA | - | |
| MofA | 4.45 ± 0.05 | - |
| Cyc2 | 12.88 ± 0.04 | 8.43 ± 0.08 |

**A_denit_20°C**

| | MtoA | MofA |
|---|---|---|
| MtoA | - | |
| MofA | -1.25 ± 1.58 | - |
| Cyc2 | 9.83 ± 2.86 | 11.08 ± 4.44 |

**A_denit_28°C**

| | MtoA | MofA |
|---|---|---|
| MtoA | - | |
| MofA | 2.03 ± 0.94 | - |
| Cyc2 | 8.53 ± 0.76 | 6.50 ± 0.57 |

**H_denit**

| | [1]MtoA | MofA |
|---|---|---|
| [1]MtoA | - | |
| MofA | x | - |
| Cyc2 | x | 6.01 |

### SI-Text 4: *Cbb* operon

Proteins encoded adjacent to *cbbM* (RubisCO) are homologous to a LysR-type transcriptional regulator (IMG ID: 2878408539), CbbQ (IMG ID: 2878408541), and CbbO (IMG ID: 2878408542). The LysR-type regulator was recognized by the IMG databank and as such annotated in the genome (IMG Taxon ID: 2878407288). Given its position directly upstream of *cbbM*, it can be further identified as *cbbR*. This is consistent with the known function of CbbR as a LysR-type transcriptional regulator of the CBB cycle, confirming the IMG annotation. The *cbbQ* and *cbbO* genes, located downstream of *cbbM*, were initially listed in the IMG annotation, as homologues of *norQ* and *norD*, respectively. However, based on the well-established *cbbMQO* operon structure in other bacteria, as well as protein sequence identities of 71% and 35% to CbbQ and CbbO from *Acidithiobacillus ferrooxidans*, these genes can be confidently assigned as *cbbQ* and *cbbO*. Thus, *F. straubiae* possesses a *cbb* operon with the structure *cbbM-cbbQ-cbbO*, with *cbbR* located directly upstream in the opposite transcriptional orientation.

### SI-Table 3: Log2FoldChange of *Cbb* operon

**Table S3: Log2FoldChange (Log2FC) of *Cbb* operon in *F. straubiae*. The four investigated conditions were: autotrophic denitrifying Fe(II) oxidation at 28°C (A_denit_28°C, n=3), heterotrophic denitrification at 28°C (H_denit, n=1), autotrophic denitrifying Fe(II) oxidation at 20°C (A_denit_20°C, n=2) and autotrophic microaerophilic Fe(II) oxidation at 20°C (A_microox, n=2). padj – adjusted *P* value.**

| IMG Gene ID | Gene name | A_denit_20°C vs A_microox | | A_denit_28°C vs A_microox | | H_denit vs A_microox | | A_denit_28°C vs H_denit | |
|---|---|---|---|---|---|---|---|---|---|
| | | Log2FC | padj | Log2FC | padj | Log2FC | padj | Log2FC | padj |
| 2878408539 | *LysR* | 2.64 | 6.10E-05 | 2.29 | 3.85E-05 | -2.34 | 0.36 | 4.63 | 6.85E-02 |
| 2878408540 | *cbbM* | 4.39 | 1.82E-11 | 3.38 | 4.49E-09 | -0.35 | 0.72 | 3.73 | 8.73E-06 |
| 2878408541 | *cbbQ* | 2.35 | 2.04E-04 | 3.44 | 2.43E-11 | -5.22 | 0.04 | 8.66 | 7.92E-04 |
| 2878408542 | *cbbO* | 1.20 | 0.02 | 3.19 | 1.73E-17 | 2.01 | 8.68E-04 | 1.18 | 0.05 |

### SI-Figure 7: Sphaeroides heme protein (SHP)-like proteins

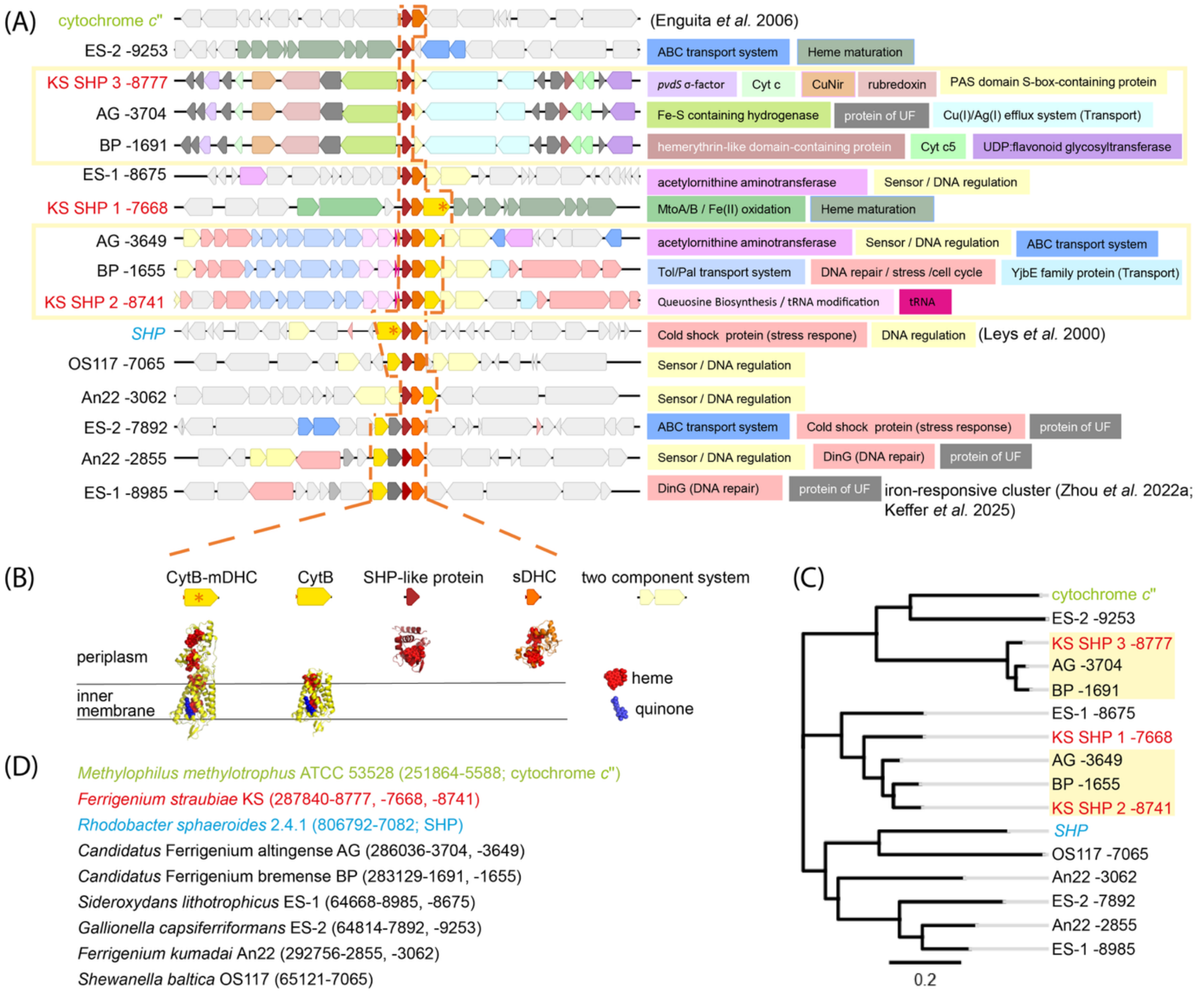

**Figure S7: (A) Gene neighborhoods (±10,000 bp) of sphaeroides heme protein (SHP)-like genes. Labels on the left indicate the strain name followed by the last four digits of the IMG gene ID of the SHP-like protein. Yellow boxes highlight genomic neighborhoods that are shared among the three main Fe(II)-oxidizing strains from autotrophic nitrate-reducing Fe(II)-oxidizing enrichment cultures. Functional annotations of neighboring genes present in at least two strains are indicated on the right and are partially grouped by matching colors. (B) AlphaFold models of representative CytB-mDHC, CytB, SHP-like protein, and sDHC structures derived from *F. straubiae* protein sequences. (C) Phylogenetic tree of SHP, cytochrome *c*'', and SHP-like proteins. (D) Full strain names and complete IMG Gene IDs.**

### SI-Figure 8: Evolutionary couplings in sphaeroides heme protein-like protein 1 (SHP 1)

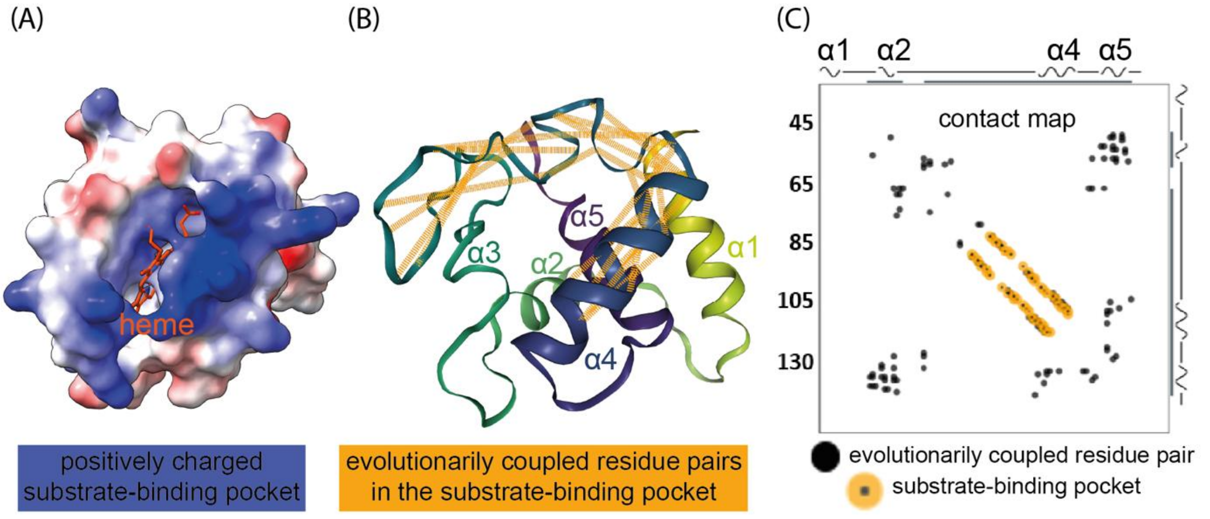

**Figure S8: (A) AlphaFold model of SHP 1 of *F. straubiae* in same orientation as structure in (B) showing evolutionary couplings within the area of potential substrate binding. (C) contact map showing evolutionary couplings from sequence covariation generated by EV couplings. Evolutionary couplings are pairwise interactions between amino acid positions in a protein sequence that coevolve due to structural or functional constraints.**

### SI-Table 4: Normalized transcripts of genes of proteins similar to cytochrome Slit_1353 of *Sideroxydans lithotrophicus* strain ES-1

**Table S4: Normalized transcripts in TPM (Transcripts Per Million) of genes encoding cytochrome in *F. straubiae* with similarity (40-55%) to the highly expressed and upregulated iron-responsive periplasmic cytochrome Slit_1353 (NCBI Gene ID) of *Sideroxydans lithotrophicus* strain ES-1 (Keffer *et al.* 2025; Zhou *et al.* 2022). The four investigated conditions were: autotrophic denitrifying Fe(II) oxidation at 28°C (A_denit_28°C, n=3), heterotrophic denitrification at 28°C (H_denit, n=1), autotrophic denitrifying Fe(II) oxidation at 20°C (A_denit_20°C, n=2) and autotrophic microaerophilic Fe(II) oxidation at 20°C (A_microox, n=2).**

| IMG Gene ID | Similarity to *Slit_1353* | Normalized transcripts in TPM | | | | | | | |
|---|---|---|---|---|---|---|---|---|---|
| | | A_microox 1 | A_microox 2 | A_denit_20°C 1 | A_denit_20°C 2 | A_denit_28°C 1 | A_denit_28°C 2 | A_denit_28°C 3 | H_denit_28°C 1 |
| 2878409861 | 40% | 111.32 | 41.76 | 875.28 | 182.88 | 885.37 | 534.15 | 592.48 | 0 |
| 2878409862 | 54% | 2304.29 | 1184.98 | 6310.35 | 812.88 | 13060.35 | 9629.58 | 9143.16 | 401.59 |
| 2878409863 | 55% | 25.62 | 10.56 | 62.89 | 8.84 | 308.63 | 224.23 | 401.86 | 1017.18 |

### SI-Table 5: Normalized transcripts of genes encoding oxygen reducing and nitric oxide reducing heme-copper oxidoreductase oxidases and their biogenesis related genes

**Table S5: Normalized transcripts in TPM (Transcripts Per Million) of genes encoding oxygen reducing and nitric oxide reducing heme-copper oxidoreductase oxidases and their biogenesis related genes in *F. straubiae*. The four investigated conditions were: autotrophic denitrifying Fe(II) oxidation at 28°C (A_denit_28°C, n=3), heterotrophic denitrification at 28°C (H_denit, n=1), autotrophic denitrifying Fe(II) oxidation at 20°C (A_denit_20°C, n=2) and autotrophic microaerophilic Fe(II) oxidation at 20°C (A_microox, n=2).**

| IMG Gene ID | Gene name | Normalized transcripts in TPM | | | | | | | |
|---|---|---|---|---|---|---|---|---|---|
| | | A_microox 1 | A_microox 2 | A_denit_20°C 1 | A_denit_20°C 2 | A_denit_28°C 1 | A_denit_28°C 2 | A_denit_28°C 3 | H_denit_28°C 1 |
| 2878407315 | *eNOR II* | 153.03 | 74.39 | 6344.74 | 244.3 | 13202.81 | 15061.76 | 10813.14 | 198.36 |
| 2878407316 | *eNOR I* | 359.32 | 185.78 | 9191.38 | 238.51 | 2493.65 | 2264.78 | 2078.26 | 315.9 |
| 2878407767 | *coxC* | 2.47 | 0.76 | 4.18 | 10.73 | 37.04 | 41.94 | 67.99 | 424.82 |
| 2878407768 | *cox11* | 5.75 | 1.98 | 3.21 | 17.15 | 34.84 | 52.97 | 70.59 | 2631.14 |
| 2878407770 | *coxA* | 8.21 | 3.52 | 2.41 | 16.81 | 30.27 | 45.55 | 61.02 | 341.21 |
| 2878407771 | *coxB* | 6 | 3.43 | 4.38 | 16.09 | 43.91 | 51.44 | 71.83 | 48.67 |
| 2878408503 | *cox15* | 13.79 | 7.45 | 143.54 | 32.84 | 140.93 | 165.73 | 177.37 | 0 |
| 2878409688 | *cox15* | 19.33 | 8.07 | 81.25 | 23.62 | 73.69 | 90.51 | 128.28 | 0 |
| 2878409749 | *ccoN* | 700.31 | 481.51 | 4221.58 | 221.01 | 6177.05 | 5246.1 | 4113.19 | 213.49 |
| 2878409750 | *ccoO* | 276.83 | 197.28 | 1982.82 | 93.14 | 3854.88 | 3799.4 | 3081.55 | 1457.27 |
| 2878409751 | *ccoQ* | 427.32 | 221.3 | 3167.58 | 140.19 | 2951.62 | 2730.9 | 2628.79 | 0 |
| 2878409752 | *ccoP* | 327.16 | 225.72 | 1555.94 | 89.64 | 5231.37 | 5730.26 | 4888.07 | 0 |

### SI-Table 6: Average expression of proteins encoded within the eNOR operon of *F. straubiae*

**Table S6 Average expression of proteins encoded within the eNOR operon of *F. straubiae*. Proteome derived from culture KS grown in Fe(II) and nitrate at 28 °C collected by Huang et al. (2026), n=1.**

| IMG Gene ID | Protein name | Average Expression |
|---|---|---|
| 2878407315 | eNOR II | 31.6 |
| 2878407316 | eNOR I | 31.6 |
| 2878407317 | heme *o* synthase | - |
| 2878407318 | Sco1 | 24.7 |
| 2878407319 | Cytbc1CCC-like protein 1 | 26.2 |
| 2878407321 | eNOR scarffold protein | 28.0 |
| 2878407322 | PerM | - |
| 2878407323 | Cytbc1CCC-like protein 2 gene | 23.9 |

### SI-Table 7: Gene abbreviation

**Table S7: Gene abbreviation with corresponding protein. Genes were listed in the dot plots.**

| Gene | Protein |
|---|---|
| *accA* | acetyl-CoA carboxylase carboxyl transferase subunit alpha |
| *accB* | acetyl-CoA carboxylase biotin carboxyl carrier protein |
| *accC* | acetyl-CoA carboxylase biotin carboxylase subunit |
| *accD* | acetyl-CoA carboxylase carboxyl transferase subunit beta |
| *aceE* | pyruvate dehydrogenase E1 component |
| *ACO 1* | aconitate hydratase |
| *ACO 2* | aconitate hydratase |
| *ATP_synth_I* | ATP synthase protein I (F-type H) |
| *ATPF0A 2* | F-type H+-transporting ATPase subunit a |
| *ATPF0B 1* | F-type H+-transporting ATPase subunit b |
| *ATPF0B 2* | F-type H+-transporting ATPase subunit b |
| *ATPF0C 1* | F-type H+-transporting ATPase subunit c |
| *ATPF0C 1* | F-type H+-transporting ATPase subunit a |
| *ATPF0C 2* | F-type H+-transporting ATPase subunit c |
| *ATPF1A 1* | F-type H+-transporting ATPase subunit alpha |
| *ATPF1A 2* | F-type H+-transporting ATPase subunit alpha |
| *ATPF1B 1* | F-type H+-transporting ATPase subunit beta |
| *ATPF1B 2* | F-type H+-transporting ATPase subunit beta |
| *ATPF1D 1* | F-type H+-transporting ATPase subunit delta |
| *ATPF1E 1* | F-type H+-transporting ATPase subunit epsilon |
| *ATPF1E 2* | F-type H+-transporting ATPase subunit epsilon |
| *ATPF1G 1* | F-type H+-transporting ATPase subunit gamma |
| *ATPF1G 2* | F-type H+-transporting ATPase subunit gamma |
| *cbbM* | ribulose-bisphosphate carboxylase large chain |
| *cbbO* | RubisCO activase CbbO |
| *cbbQ* | RubisCO activase CbbQ |
| *ccoN* | cytochrome c oxidase cbb3-type subunit 1 |
| *ccoO* | cytochrome c oxidase cbb3-type subunit 2 |
| *ccoP* | cytochrome c oxidase cbb3-type subunit 3 |
| *cox11* | cytochrome c oxidase assembly protein subunit 11 |
| *cox15* | cytochrome c oxidase assembly protein subunit 15 |
| *coxA* | cytochrome c oxidase subunit 1 |
| *coxB* | cytochrome c oxidase subunit 2 |
| *coxC* | cytochrome c oxidase subunit 3 |
| *CS* | citrate synthase |
| *cyc2* | outer membrane cytochrome c Cyc2 |
| *cynT 1* | carbonic anhydrase |
| *cynT 2* | carbonic anhydrase |
| *cynT 3* | carbonic anhydrase |
| *cyt c"* | cytochrome c" |
| *cyt1 copy 1* | ubiquinol-cytochrome c reductase cytochrome c1 subunit |
| *cyt1 copy 2* | ubiquinol-cytochrome c reductase cytochrome c1 subunit |

| | |
|---|---|
| ***cytB*** | trans membrane cytochrome b |
| ***cytb copy 1*** | ubiquinol-cytochrome c reductase cytochrome b subunit |
| ***cytb copy 2*** | ubiquinol-cytochrome c reductase cytochrome b subunit |
| ***cytB-mDHC*** | trans membrane cytochrome b + membrane diheme cytochrome c |
| ***DLAT*** | pyruvate dehydrogenase E2 component (dihydrolipoamide acetyltransferase) |
| ***DLD*** | dihydrolipoamide dehydrogenase |
| ***DLST*** | 2-oxoglutarate dehydrogenase E2 component (dihydrolipoamide succinyl transferase) |
| ***eNOR I*** | clade e nitric oxide reductase subunit 1 |
| ***eNOR II*** | clade e nitric oxide reductase subunit 2 |
| ***exbB*** | biopolymer transport protein ExbB |
| ***exbD*** | biopolymer transport protein ExbD |
| ***FBA*** | fructose-bisphosphate aldolase |
| ***FBP*** | fructose-1,6-bisphosphatase I |
| ***fdoG 1*** | formate dehydrogenase major subunit |
| ***fdoG short 2*** | formate dehydrogenase major subunit |
| ***fdoH*** | formate dehydrogenase iron-sulfur subunit |
| ***fdoI*** | formate dehydrogenase subunit gamma |
| ***fecR sigma factor*** | Ferric citrate regulatory protein |
| ***feoA*** | ferrous iron transport protein A |
| ***feoB*** | ferrous iron transport protein B |
| ***FH class I*** | fumarate hydratase class I |
| ***FH class I*** | fumarate hydratase class I |
| ***FH class II*** | fumarate hydratase class II |
| ***FH class-II*** | fumarate hydratase class II |
| ***fixJ*** | FixJ family two-component response regulator |
| ***folD*** | methylenetetrahydrofolate dehydrogenase (NADP+)/methenyltetrahydrofolate cyclohydrolase |
| ***GAPDH*** | glyceraldehyde 3-phosphate dehydrogenase |
| ***gcvH*** | glycine cleavage system H protein |
| ***gcvPA*** | glycine dehydrogenase subunit 1 |
| ***gcvPB*** | glycine dehydrogenase subunit 2 |
| ***gcvT*** | aminomethyltransferase |
| ***glyA*** | glycine hydroxymethyltransferase |
| ***IDH1*** | isocitrate dehydrogenase |
| ***ilvA*** | threonine dehydratase |
| ***ISP copy 1*** | ubiquinol-cytochrome c reductase iron-sulfur subunit |
| ***ISP copy 2*** | ubiquinol-cytochrome c reductase iron-sulfur subunit |
| ***korA*** | 2-oxoglutarate ferredoxin oxidoreductase subunit alpha |
| ***korB*** | 2-oxoglutarate ferredoxin oxidoreductase subunit beta |
| ***LysR*** | DNA-binding transcriptional LysR family regulator |
| ***LysR 1*** | LysR family cys regulon transcriptional activator |
| ***LysR 3*** | LysR family cys regulon transcriptional activator |
| ***mdh*** | malate dehydrogenase |
| ***metF*** | methylenetetrahydrofolate reductase (NADPH) |
| ***mofA*** | Mn-multicopper oxidase A |
| ***mofB*** | Chaperone MofB |

| | |
|---|---|
| ***mofC*** | Cytochrome c MofC |
| ***mtoA*** | Metal-oxidase multi-heme subunit MtoA |
| ***mtoB*** | Metal-oxidase beta-barrel subunit MtoB |
| ***narG*** | nitrate reductase alpha subunit |
| ***narH*** | nitrate reductase beta subunit |
| ***narI*** | nitrate reductase gamma subunit |
| ***narJ*** | nitrate reductase delta subunit |
| ***narK copy1*** | NNP family nitrate/nitrite transporter-like MFS transporter |
| ***narK copy2*** | NNP family nitrate/nitrite transporter-like MFS transporter |
| ***narL/fixJ*** | DNA-binding NarL/FixJ family response regulator |
| ***NhaB family antiporter*** | NhaB family Na+:H+ antiporter |
| ***NhaP-type antiporter 1*** | NhaP-type Na+/H+ or K+/H+ antiporter |
| ***NhaP-type antiporter 2*** | NhaP-type Na+/H+ or K+/H+ antiporter |
| ***nirK*** | copper-containing nitrite reductase (nirK) |
| ***nirS copy1*** | nitrite reductase (NO-forming)/hydroxylamine reductase (nirS) |
| ***nirS copy2*** | nitrite reductase (NO-forming)/hydroxylamine reductase (nirS) |
| ***NtrC family*** | DNA-binding NtrC family response regulator |
| ***nuoA*** | NADH-quinone oxidoreductase subunit A |
| ***nuoB*** | NADH-quinone oxidoreductase subunit B |
| ***nuoC*** | NADH-quinone oxidoreductase subunit C |
| ***nuoD*** | NADH-quinone oxidoreductase subunit D |
| ***nuoE*** | NADH-quinone oxidoreductase subunit E |
| ***nuoF*** | NADH-quinone oxidoreductase subunit F |
| ***nuoF*** | NADH-quinone oxidoreductase subunit F |
| ***nuoG*** | NADH-quinone oxidoreductase subunit G |
| ***nuoH*** | NADH-quinone oxidoreductase subunit H |
| ***nuoI*** | NADH-quinone oxidoreductase subunit I |
| ***nuoJ*** | NADH-quinone oxidoreductase subunit J |
| ***nuoK*** | NADH-quinone oxidoreductase subunit K |
| ***nuoL*** | NADH-quinone oxidoreductase subunit L |
| ***nuoM*** | NADH-quinone oxidoreductase subunit M |
| ***nuoN*** | NADH-quinone oxidoreductase subunit N |
| ***OGDH*** | 2-oxoglutarate dehydrogenase E1 component |
| ***ompR*** | two-component system OmpR family response regulator/two-component system response regulator QseB |
| ***PerM*** | predicted PurR-regulated permease PerM |
| ***PGK*** | phosphoglycerate kinase |
| ***pirA*** | outer membrane receptor protein involved in Fe transport |
| ***pirA*** | iron complex outermembrane receptor protein |
| ***pntA*** | NAD(P) transhydrogenase subunit alpha |
| ***pntB*** | NAD(P) transhydrogenase subunit beta |
| ***por*** | pyruvate-ferredoxin/flavodoxin oxidoreductase |
| ***ppc*** | phosphoenolpyruvate carboxylase |
| ***ppdK*** | pyruvate,orthophosphate dikinase |
| ***PRK*** | phosphoribulokinase |
| ***pvdR*** | HlyD family secretion protein |

| | |
|---|---|
| ***pvdS sigma factor*** | RNA polymerase sigma-70 factor (ECF subfamily) |
| ***pvdT*** | putative ABC transport system permease protein |
| ***pycA*** | pyruvate carboxylase subunit A |
| ***pycB*** | pyruvate carboxylase subunit B |
| ***qseC*** | two-component system sensor histidine kinase QseC |
| ***rnfA 1*** | electron transport complex protein RnfA |
| ***rnfA 2*** | electron transport complex protein RnfA |
| ***rnfB 1*** | electron transport complex protein RnfB |
| ***rnfB 2*** | electron transport complex protein RnfB |
| ***rnfC 1*** | electron transport complex protein RnfC |
| ***rnfC 2*** | electron transport complex protein RnfC |
| ***rnfD 1*** | electron transport complex protein RnfD |
| ***rnfD 2*** | electron transport complex protein RnfD |
| ***rnfE 1*** | electron transport complex protein RnfE |
| ***rnfE 2*** | electron transport complex protein RnfE |
| ***rnfG 1*** | electron transport complex protein RnfG |
| ***rnfG 2*** | electron transport complex protein RnfG |
| ***rpe*** | ribulose-phosphate 3-epimerase |
| ***rpiA*** | ribose 5-phosphate isomerase A |
| ***sdhA*** | succinate dehydrogenase / fumarate reductase flavoprotein subunit |
| ***sdhB*** | succinate dehydrogenase / fumarate reductase iron-sulfur subunit |
| ***sDHC*** | soluble diheme cytochrome c |
| ***Sco1*** | synthesis of cytochrome C oxidase 1 |
| ***sdhC*** | succinate dehydrogenase / fumarate reductase cytochrome b subunit |
| ***sdhD*** | succinate dehydrogenase / fumarate reductase membrane anchor subunit |
| ***shp*** | sphaeroides heme protein |
| ***SHP 1*** | sphaeroides heme protein-like protein |
| ***SHP 2*** | sphaeroides heme protein-like protein |
| ***SHP 3*** | sphaeroides heme protein-like protein |
| ***sucC*** | succinyl-CoA synthetase beta subunit |
| ***sucD*** | succinyl-CoA synthetase alpha subunit |
| ***TALDO1*** | transaldolase |
| ***TIP*** | triosephosphate isomerase |
| ***TKT 1*** | transketolase |
| ***TKT 2*** | transketolase |
| ***tonB*** | protein TonB |
